# Ecological and Evolutionary Characteristics of Anthropogenic Roosting Ability in Bats of the World

**DOI:** 10.1101/2023.10.15.562433

**Authors:** Briana A. Betke, Nicole L. Gottdenker, Lauren Ancel Meyers, Daniel J. Becker

## Abstract

Although the global conversion of wildlife habitat to built environments often has negative impacts on biodiversity, some wildlife have the ability to cope by living in human-made structures. However, the determinants of this adaptation on a global scale are not well understood and may signify species with unique conservation needs at the human–wildlife interface. Here, we identify the trait profile associated with anthropogenic roosting in bats globally and characterize the evolution of this phenotype using an original dataset of roosting behavior developed across 1,279 extant species. Trait-based analyses showed that anthropogenic roosting is predictable across bats and is associated with habitat generalism, small geographic ranges, small body size, temperate zone distributions, and insectivory. We identified moderate phylogenetic signal in this complex phenotype, which has undergone both gains and losses across bat evolution and for which speciation rates are lower compared to natural roosting bats.

## Introduction

Human influences on environmental conditions (e.g., climate change, urbanization, deforestation, agriculture) impact wildlife by altering community composition, resource availability, and available habitat.^1–4^ As a result, wildlife health can suffer through exposure to toxins and pathogens,^5^ vehicle collisions,^6^ predation by domestic animals,^7,8^ and competition with invasive species.^7^ Despite these risks, some wildlife species thrive in the midst of human disturbance, benefiting from anthropogenic resources such as shelter, supplementary food, and protection from predation.^9,10^ For example, urban environments provide the coyote (*Canis latrans*) with an ample diet of small mammals (e.g., rabbits, squirrels, domestic cats) and vegetation (e.g., ornamental fruits), as well as protection from apex predators.^11,12^ Researchers have associated various functional traits that facilitate the observed tolerance of some wildlife species to human disturbance.^13–21^ However, most studies do not consider the propensity to occupy anthropogenic structures when measuring tolerance (e.g., buildings, bridges, houses), often categorizing species broadly by urban presence (i.e., defined as dwellers, avoiders, or visitors) or dichotomizing species as urban or non-urban.^16,22^ Such structures serve as an increasingly critical frontier where human–wildlife conflict can occur.^23,24^ Identifying the species that inhabit anthropogenic structures and their functional traits that facilitate usage can inform more effective wildlife conservation through conflict mitigation efforts.

Wildlife use a variety of anthropogenic structures as habitat including roofs,^25–29^ crawl spaces and under floorboards,^25,30^ chimneys,^31,32^ bridges,^33–35^ road culverts,^33,34^ dumpsters,^36^ and power line pylons.^37,38^ Some species live in these structures occasionally while others rely on them as primary habitation. For instance, European stone martens (*Martes foina*) live almost exclusively in buildings, and large-spotted genets (*Genetta tigrina*) roost in roofs of houses with increasing occurrence in South African suburbs, whereas Madagascar free-tailed bats (*Otomops madagascariensis*) are obligate cave-dwellers.^28,39,40^ Wildlife that inhabit anthropogenic structures may be threatened by extirpation originating from human-wildlife conflict.^23^ Such threats include the potential transmission of zoonotic pathogens and parasites that cause disease in humans and companion animals (e.g., rabies, tuberculosis, roundworm, leptospirosis, salmonella).^41,42^ Additionally, wildlife species may be perceived as pests due to property damage,^23,43^ attacks on humans and companion animals,^44,45^ the off-putting odors of excreta near or in homes.^46,47^ Identifying structures and the species that commonly use them can help tailor conservation efforts. For example, such actions could include preserving existing structures for vulnerable wildlife, developing structures that would be more suitable for dwelling, and targeting efforts that reduce conflict with humans.

While there are many reports of wildlife use of anthropogenic structures, global patterns of this behavior are largely unknown. Most studies typically make intraspecific rather than interspecific comparisons, focus on a local or regional scale which limit data available for global biogeographic comparisons, and do not consider underlying determinants of habitat choice beyond microclimatic conditions. Although mass data collection remains cost prohibitive, meta-analysis of existing global data can help to overcome some of these limitations and robustly identify the foundational biological principles that determine anthropogenic habitat usage.

Bats are an ideal model for studying functional traits associated with living in anthropogenic structures, given their high species diversity, global geographic distribution, and the substantial research investment to understand and address conservation and public health risks associated with various bat species. The order Chiroptera constitutes 20% of all mammalian species and includes a high diversity of physical characteristics, life history, and ecology.^48^ They roost in a variety of structures owing to wide interspecific variation in roosting requirements ranging from natural (e.g., caves, foliage, rock crevices, banana leaves, tree canopy, and crevices) to anthropogenic (e.g, buildings, bridges, and mines).^49,50^ Human-made structures provide similar conditions to natural roosts that have been lost to rapid land use changes, offering protection from predation, shelter from extreme weather, and stable temperatures for improved reproductive success.^50,51^ However, some bat species suffer adverse effects when moving from natural to anthropogenic habitats, such as increased vulnerability to predation, and unstable roost microclimates,^26,52,53^ and extermination by humans who perceive them as nuisance wildlife and reservoirs of zoonotic pathogens.^54–57^ Due to these negative perceptions, bats’ ability to provide essential ecosystem services (e.g., pollination, seed dispersal, maintenance of insect populations) and bioindicator status is often overlooked.^58^ On a local scale, there is evidence that identifying anthropogenic roosts can prioritize important bat conservation sites and reduce bat–human conflict.^59–61^ Identifying global trends in this behavior may allow for application of conflict mitigation on a global scale.

Here, we compiled a novel and comprehensive dataset documenting roosting structure type of 1,279 bat species, based on historical records. We first describe geographic and taxonomic patterns in anthropogenic roosting ability. Next, we use a machine learning algorithm, boosted regression trees (BRTs), to identify ecological and evolutionary characteristics (i.e., the trait profile) of anthropogenic roosting across bats globally, leveraging data from PanTHERIA,^62^ COalesced Mammal dataBase of INtrinsic and Extrinsic traits (COMBINE),^63^ and the International Union for the Conservation of Nature (IUCN).^64,65^ BRTs side-step many challenges associated with analyzing such ecological datasets, including nonindependent data, non-randomly missing covariates, nonlinear patterns, and complex interactions, and demonstrate improved classifier ability than traditional regression methods.^66^ Finally, we characterize the phylogenetic distribution and evolution of this complex and multifaceted phenotype. This dataset provides a new window into global trends of anthropogenic roosting and traits underlying anthropogenic dwelling that may not be apparent in data reflecting only presence/absence of bats in human dominated landscapes.

## Results

### Dataset description

Of the most known diversity of bats (over 1,460 species to date)^67^, we examined 1,279 extant bat species included in the most recent mammal phylogeny,^68^ our systematic dataset included the presence or absence of anthropogenic roosting across 80% of this order (*n* = 1,042). We find that approximately 40% of extant bat species are capable of roosting in anthropogenic structures, although the occurrence of anthropogenic roosting varies markedly across bat families, biogeographic regions, and conservation status (Fig. 1). Anthropogenic roosting structures included bridges, attics, mines, and overhangs of buildings. The proportion of anthropogenic roosting species per family ranged from 0% to 100% (Fig. 1A). Several small families have 100% coverage with all species labeled as anthropogenic (i.e., Rhinopomatidae, Noctilionidae, and Megadermatidae). Nycteridae, Molossidae, and Furipteridae contain 75%, 57%, and 50% anthropogenic roosting bat species, respectively. Three families (i.e., Thyropteridae, Myzopodidae, and Craseonycteridae) have no known anthropogenic roosting species. Of the biogeographical regions (Fig. 1B), the proportion of anthropogenic bats recorded was lowest in the Oceanian region (0.09%) and highest in the Nearctic region (79%). Fifty-two percent of bat species with a conservation status of least concern were found to roost in anthropogenic structures, followed by 39% of near threatened species (Fig. 1C); the critically endangered species had the lowest proportion (14%).

**Figure 1.**
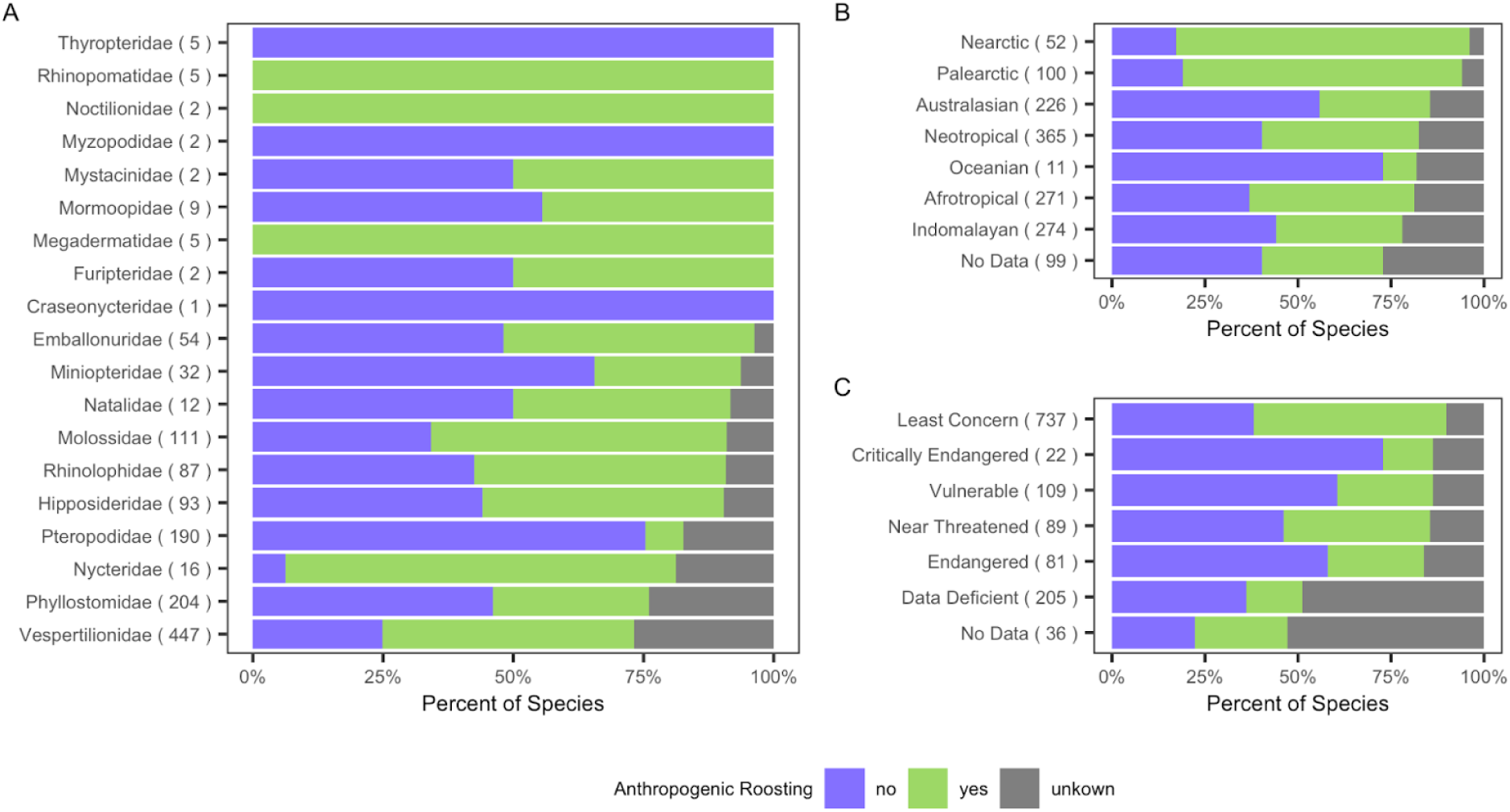
Proportion of species by roosting type across bat families (A), biogeographic region (B), and IUCN conservation status (C). The colors indicate the proportion of bats that are natural roosting (purple), anthropogenic roosting (green), and have no roosting data available (gray). The numbers located at the end of the bars show the total number of species in each category.

### Trait profile of anthropogenic roosting ability

To identify the trait profile of an anthropogenic roosting bat species, we evaluated 61 features of bat life history, geography, diet, phylogeny, and conservation status (Fig. 2) with a BRT model. Our model distinguished anthropogenic and natural roosting species with high accuracy and specificity, and moderate sensitivity (AUC = 0.95, sensitivity = 0.84, specificity = 0.89). The top features for classifying anthropogenic roosting species included geography (i.e., geographic range size, habitat breadth, mean monthly precipitation, mean monthly Actual Evapotranspiration Rate (AET), mean monthly Potential Evapotranspiration Rate (PET), maximum latitude, mean human population density), life history (i.e., litter size, adult body length, adult body mass, adult forearm length), diet (i.e., percentage of plants, fruit, invertebrates), and conservation status (Fig. 2). In contrast, taxonomic predictors had lower relative importance, with family-level traits each having less than 1% relative importance when accounting for all other traits. Importantly, citation count also had low importance, suggesting that the traits identified are not biased by data availability.

**Figure 2.**
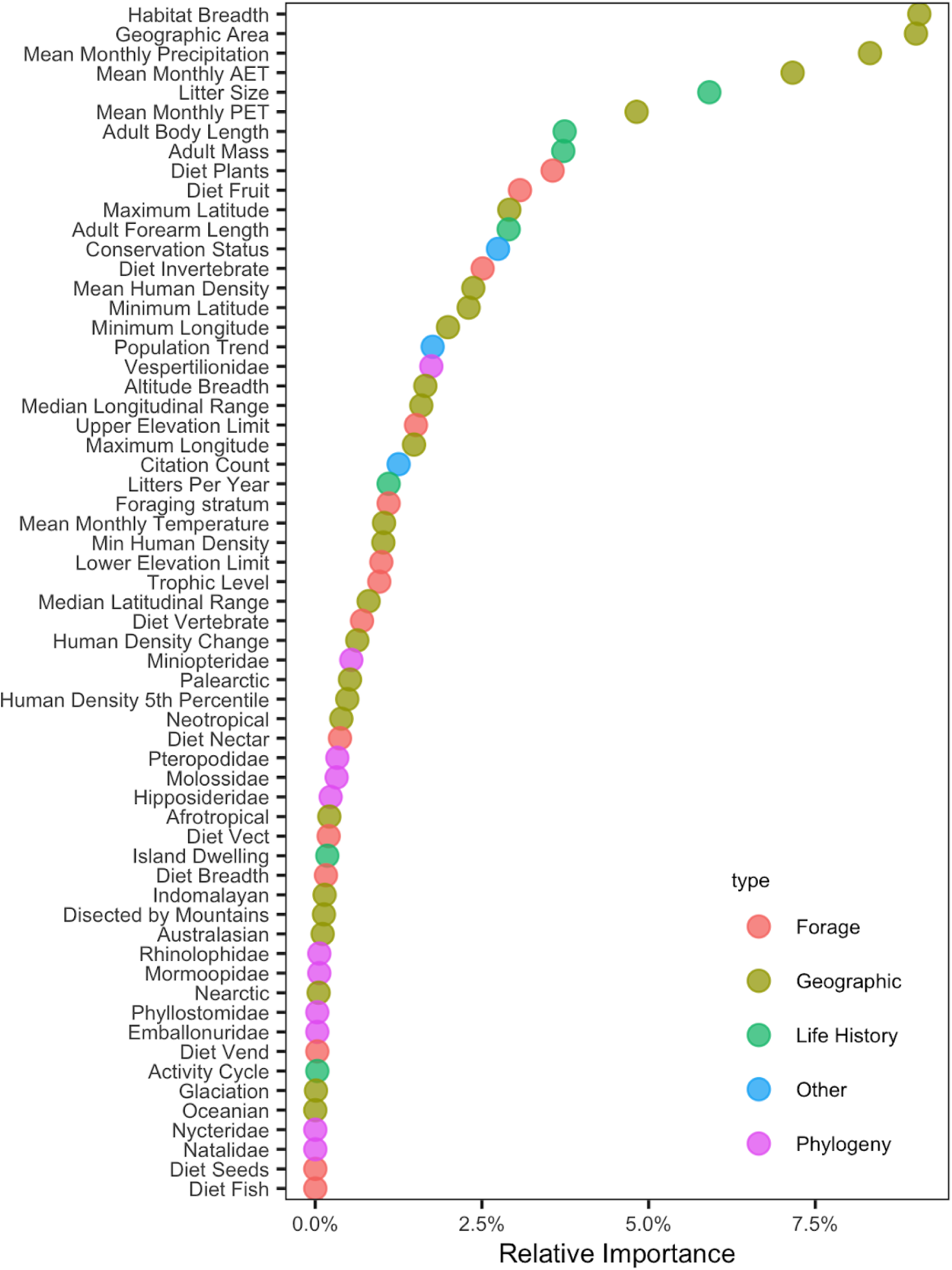
Relative importance of trait covariates from the boosted regression tree model of roosting status. Traits are ordered by ranking, with the highest at the top, and colors in the legend correspond to variable type. AET is the Actual Evapotranspiration Rate and PET is the Potential Evapotranspiration Rate.

Partial dependence plots (Fig. 3) describe anthropogenic roosting species as being habitat generalists that occupy relatively smaller geographic ranges with low monthly AET and precipitation. Anthropogenic roosting bats are also located further from the equator, showing presence at both extremes of latitude, especially in the northern hemisphere. This phenotype was further described by smaller litter sizes and smaller body size overall, as measured by adult body mass, forearm length, and body length. Additionally, the diet of an anthropogenic roosting bat consisted of a low percentage of plants, low percentage of fruit, and a high percentage of invertebrates. Of all IUCN conservation status categories, species listed as least concern were the most likely to be classified as anthropogenic roosting, followed by endangered species.

**Figure 3.**
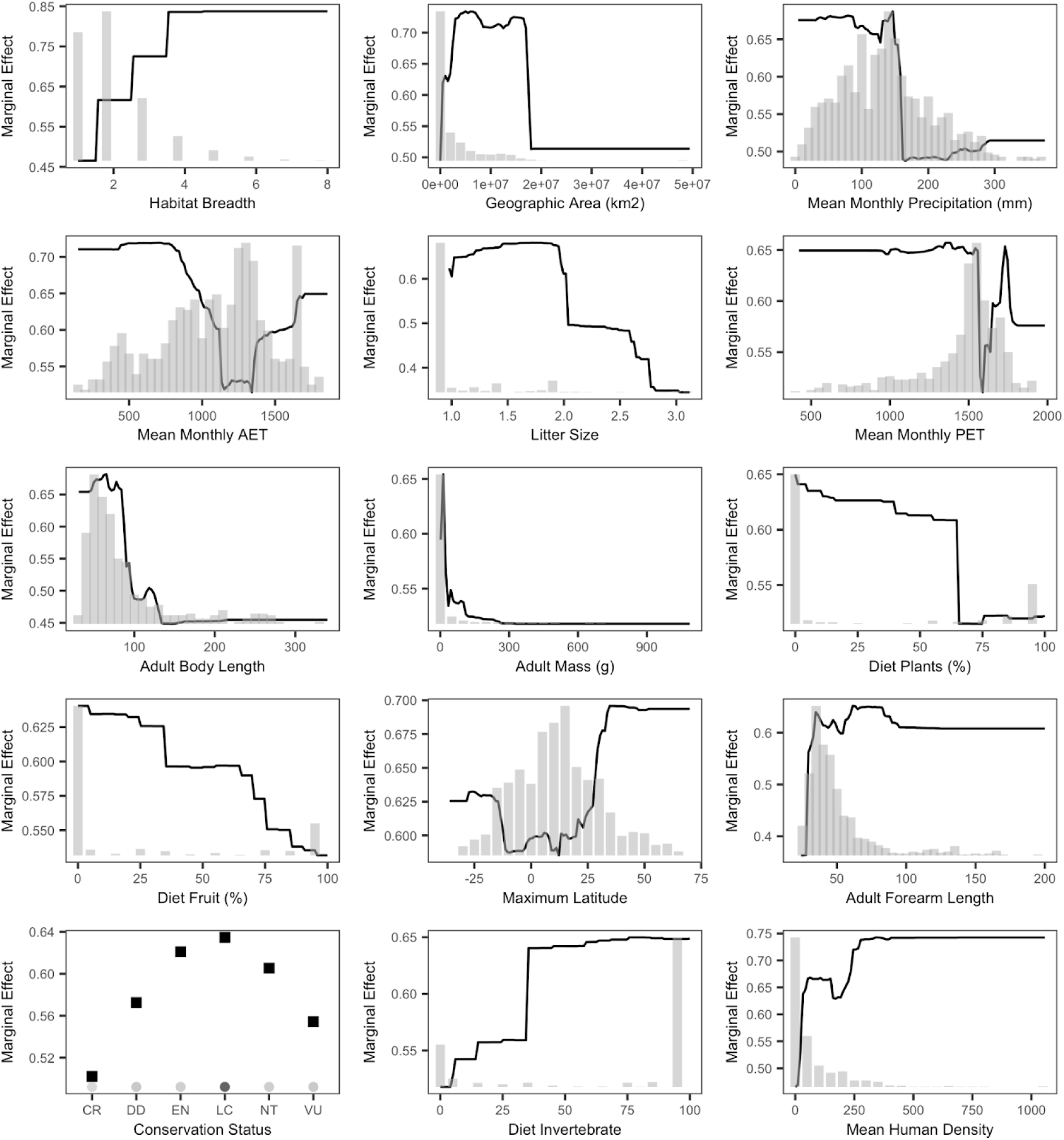
Trait profile of anthropogenic roosting bats. Partial dependence plots of the top predictors ordered by relative importance. The black line displays the marginal effect of a given variable for the prediction of roosting status. Histograms and point color show the distribution of the continuous and categorical predictors, respectively, across the 1,029 bat species. The conservation status abbreviations are as follows: Critically Endangered (CR); Data Deficient (DD), Endangered (EN), Least Concern (LC), Near Threatened (NT), Vulnerable (VU).

We assessed the sensitivity of these results to pseudoabsences, species for which adequate information on roosting status does not exist (*n* = 237). The BRT model containing pseudoabsences (*n* = 1,279 species) performed similarly to the above BRT model (*n* = 1042 species) but had lower sensitivity and higher specificity (AUC = 0.95, sensitivity = 0.80, specificity = 0.92). With the exception of citation count, variable importance between models were relatively similar (Fig. S1A), showing little variation in variable importance (Fig. S1B) and strong correlation (ρ = 0.95, *p* < 0.001). Partial dependence plots for the pseudoabsence model revealed a similar trait profile of anthropogenic roosting ability compared to the initial model (Fig. S2), suggesting inclusion of pseudoabsences did not qualitatively influence results.

### Evolution of anthropogenic roosting

As our BRT model demonstrated, anthropogenic roosting was a predictable trait on the basis of bat geography, life history, and diet. We next quantified the phylogenetic signal in this multifaceted phenotype and tested how it has evolved. Anthropogenic roosting showed low-to-moderate phylogenetic signal across bats (*D* = 0.65) and departed from both Brownian motion and phylogenetically independent models of evolutionary processes (*p* < 0.001). We also used a graph-partitioning algorithm, phylogenetic factorization, to flexibly identify clades with different propensity to roost in anthropogenic structures at various taxonomic depths.^69^

Phylogenetic factorization identified two clades of bats with markedly lower propensities to harbor anthropogenic roosting phenotypes compared to the rest of the bat phylogeny, after adjusting for differential sampling effort: the family Pteropodidae (9%) and the phyllostomid subfamily Stenodermatinae (25%; Fig. 4). Results were consistent when including pseudoabsences for the 237 understudied bat species, with similar estimates of phylogenetic signal (*D* = 0.70) and equivalent identification of both the Pteropodidae and Stenodermatinae as less likely to roost in anthropogenic structures (Table S1).

**Figure 4.**
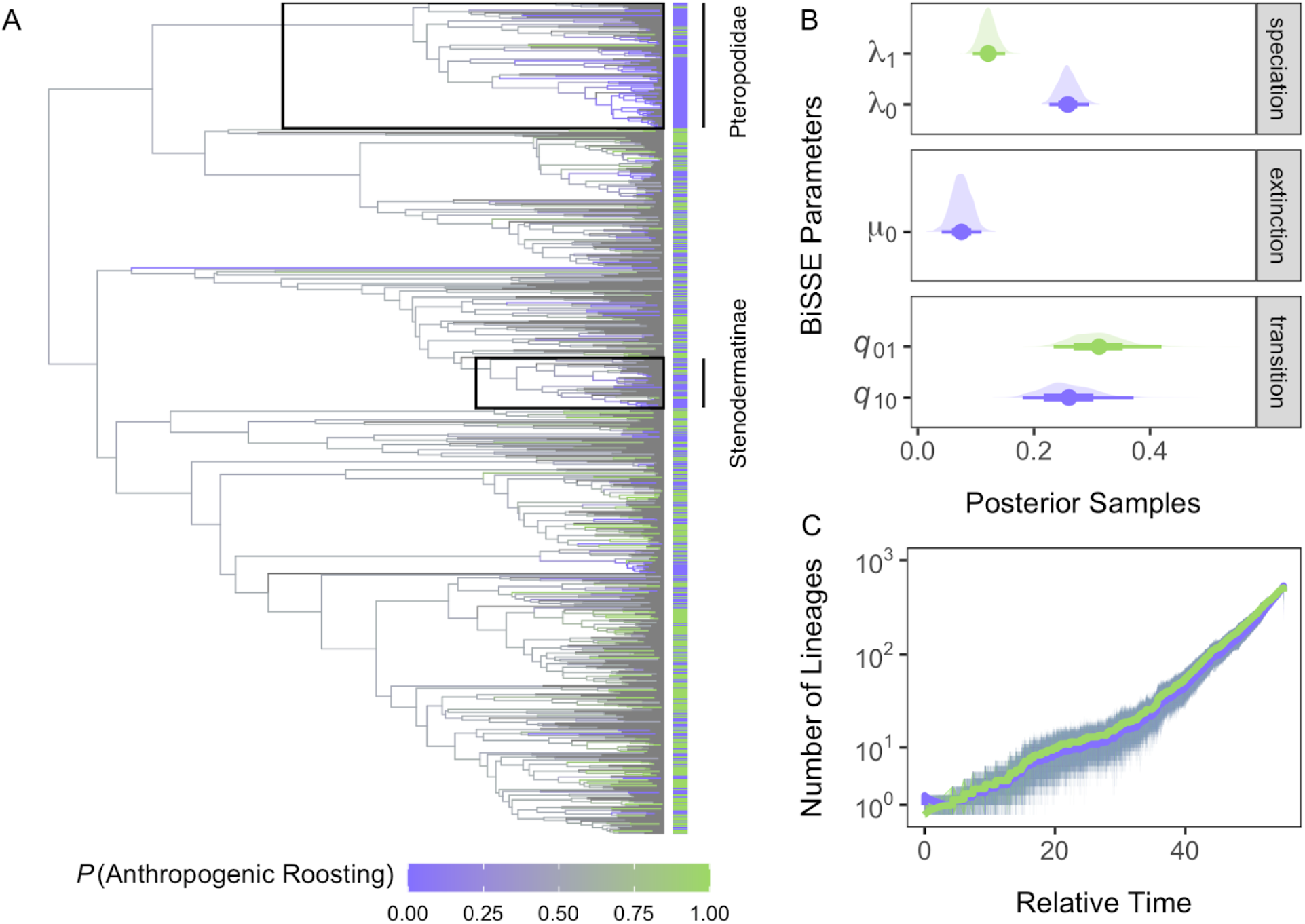
Evolution of anthropogenic roosting across bats. The extant bat phylogeny with tips colored by known roosting status (*n* = 1,029), with branches colored by posterior probabilities for anthropogenic roosting derived from stochastic character mapping of the transition matrix from the top BiSSE model. BiSSE parameters with posterior means, 66% and 95% credible intervals, and posterior densities are shown from MCMC sampling alongside individually simulated and averaged lineages through time for natural and anthropogenic roosting states.

We compared among multiple binary state speciation and extinction (BiSSE) models to assess whether the phenotypes that enable anthropogenic roosting have facilitated bat diversification.^70^ Of eight candidate models where speciation, extinction, and transition rates could vary between natural and anthropogenic roosting states, we found the most support for a constrained model with trait-dependent speciation and transitions but not extinction (λ_0_ ≠ λ_1_, μ_0_ = μ_1_, *q*_01_ ≠ *q*_10_; *w_i_* = 0.60). Models with trait-dependent extinction and irreversible models (i.e., *q*_10_ = 0) consistently received little information theoretic support (Table 1). MCMC sampling of the top BiSSE model demonstrated that speciation rates were greater in natural roosting bats than in anthropogenic roosting bats, as 95% credible intervals did not overlap (Fig. 4). Transitions from natural to anthropogenic roosting were slightly more frequent than transitions from anthropogenic to natural roosting, with less posterior support. Ancestral state reconstruction using the estimated posterior mean transition rates and stochastic character mapping further emphasized relatively more gains of natural roosting lineages across bat evolution (Fig. 4).

**Table 1.** Comparison among eight BiSSE models of trait-dependent diversification, including speciation (λ), extinction (µ), and transitions between binary states (*q*). Models are ranked by difference in Akaike information criterion from the top model (ΔAIC), the number of estimated parameters (*k*), and Akaike weights (*w_i_*).

| Model | $\Delta\text{AIC}$ | $k$ | $w_i$ |
| --- | --- | --- | --- |
| $\lambda_0 \neq \lambda_1, \mu_0 = \mu_1, q_{01} \neq q_{10}$ | 0.00 | 5 | 0.60 |
| $\lambda_0 \neq \lambda_1, \mu_0 \neq \mu_1, q_{01} \neq q_{10}$ | 0.85 | 6 | 0.40 |
| $\lambda_0 = \lambda_1, \mu_0 \neq \mu_1, q_{01} \neq q_{10}$ | 16.17 | 5 | <0.01 |
| $\lambda_0 = \lambda_1, \mu_0 = \mu_1, q_{01} = q_{10}$ | 17.64 | 3 | <0.01 |
| $\lambda_0 = \lambda_1, \mu_0 = \mu_1, q_{01} \neq q_{10}$ | 19.58 | 4 | <0.01 |
| $\lambda_0 \neq \lambda_1, \mu_0 \neq \mu_1, q_{10} = 0$ | 127.45 | 5 | <0.01 |
| $\lambda_0 = \lambda_1, \mu_0 \neq \mu_1, q_{10} = 0$ | 149.24 | 4 | <0.01 |
| $\lambda_0 \neq \lambda_1, \mu_0 = \mu_1, q_{10} = 0$ | 276.98 | 4 | <0.01 |
| $\lambda_0 = \lambda_1, \mu_0 = \mu_1, q_{10} = 0$ | 499.62 | 3 | <0.01 |

Inclusion of anthropogenic roosting pseudoabsences did not affect selection of the top BiSSE model, with similar support for trait-dependent speciation and transition rates but not extinction and no support for irreversible models (Table S2). MCMC sampling suggested similarly strong differences in speciation rates between anthropogenic and natural roosting bats, with again weak posterior support for differences in transition rates (although this analysis suggested more frequent losses of anthropogenic roosting than gains of this trait; Fig. S3).

## Discussion

As built environments expand with human population growth ^71^, the risks of human-wildlife conflict increase. In this study, we characterize the trait profile of anthropogenic roosting bats to support the detection of bat species at the human-wildlife interface and potentially mitigate associated risks. We compiled the largest curated database of anthropogenic roosting across all bat families, biogeographical realms, and conservation status, and then used the data to assess the determinants of anthropogenic roosting with fundamental ecological and evolutionary characteristics. The data are standardized to integrate seamlessly with the COMBINE, PanTHERIA and IUCN trait databases.

Our analyses suggest that anthropogenic roosting can be predicted by species traits, with such bats having broad habitat breadth but relatively small geographic ranges, occurring in temperate regions, tending to have small body size, having low reproductive output, and consuming more invertebrates than plants. We also identified moderate phylogenetic effects, with particular clades having lower propensities to roost in anthropogenic structures and that this anthropogenic roosting phenotype has undergone losses and gains throughout bat evolution.

Considering all traits included in our BRT model, geographic range was the most important predictor of anthropogenic roosting ability. Our BRT analysis suggested that species with small geographic ranges are more likely to roost in anthropogenic structures, possibly owing to these species having the smaller body sizes necessary to use these unique roosts.

Habitat breath, a common trait for determining tolerance to human disturbance in other taxa,^18,21,72–74^ was second in importance. Specifically, anthropogenic roosting ability was associated with having a broad habitat breath. Both geographic range size and habitat breadth play important roles in determining extinction, with species having smaller geographic ranges and narrower habitat breadths being more vulnerable to extinction.^75–77^ Although anthropogenic roosting bats may have small geographic ranges and small body sizes, they may be able to persist in human environments due to their ability to exist in many habitat types. This result emphasizes the likely importance of anthropogenic structures in the survival of bat species that do not fare well outside of their geographic range.

Anthropogenic roosting bats also are rarer in the tropics compared to temperate areas. This finding is consistent with the larger proportions of these bats observed in the Nearctic and Palearctic biogeographical regions in our descriptive analysis. Our model suggested that anthropogenic roosting bats occur mostly in northern temperate regions with low precipitation. Additionally, anthropogenic roosting bats occur in areas with mostly low to moderate water loss, as suggested by partial dependence plots of AET and PET. However, these plots also showed a spike in anthropogenic roosting propensity at the higher extremes of AET and PET, potentially suggesting that bat species with this trait persist in areas within the temperate region that are drier and experience greater water loss.

Although human occupancy is a prerequisite for bats to roost in anthropogenic structures, our study showed that human population density and associated metrics (i.e., minimum human population density, human density of the 5th percentile, and human population density change) are not among the top 10 predictors of the anthropogenic roosting trait profile. Mean human population density was ranked as the 15th most important factor, and anthropogenic roosting occurred at almost any human density, spanning rural to urban. This finding suggests that use of anthropogenic structures is better determined by geographic, life history, and dietary characteristics of individual species.

We also found anthropogenic bats have smaller litter sizes, smaller body mass, and smaller body length. The association between smaller litter size and anthropogenic roosting seemingly contradicts some prior studies of urban tolerance, where greater fecundity is instead associated with urban living.^16,21,73^ Our models suggest there may be reproductive pressure towards smaller litters to live in spaces provided by anthropogenic structures. Larger litter sizes of urban-tolerant species are likely an adaptive response to cope with excess mortality unique to urban spaces (e.g., vehicle collisions, human–wildlife conflict, and predation by cats).^16^ However, anthropogenic roosting bats may be less impacted by certain external sources of mortality such as vehicle collisions and predation, whereas mortality may instead result from extirpation from human-made structures. Our finding of small body size in anthropogenic roosting bats also may reflect some unique facets of bat biology. Having a relatively small body size facilitates the ability to fit in crevices of anthropogenic structures and take up less space compared to species with larger body sizes,^78^ which could buffer these bats from external sources of predation in urban habitats. Interestingly, we observed a peak effect for forearm size at small lengths, which may enhance maneuverability in cramped spaces.

Our analysis also showed that frugivorous bats are unlikely to be anthropogenic roosting. Rather, bat species that consume mostly invertebrates, typically insects, are more prone to roost in anthropogenic structures. Several reasons could explain this pattern. First, the conditions of human-dominated environments also provide resources that insectivorous bats can exploit to persist in human-made structures. Built environments attract insects through the presence of standing water (e.g., ponds, sprinklers, birdbaths), plants (e.g., gardens, ornamental/landscaping plants, crops), and light sources.^79^ Insectivorous bats have wide behavioral or phenotypic plasticity that allow for survival in anthropogenic structures. For example, insectivorous bats can adjust features of their echolocation^80^ and alter foraging activity in anthropogenically disturbed areas ^81^. Finally, anthropogenic structures may not be suitable roost sites for frugivorous bats, which tend to have larger body sizes that could physically limit use of many anthropogenic roosts. We conducted a post-hoc analysis of our trait data using phylogenetic least squares models showed that forearm length is positively correlated with fruit consumption (*β* = 0.18, p < 0.0001) and negatively with invertebrate consumption (*β* = -0.15, p < 0.001). This explanation is supported by prior evolutionary analyses, as multiple subclades of Pteropodidae (a frugivorous family) showed rapid increases in body sizes throughout their evolution.^82^

Whereas prior studies have assessed the traits underlying species propensities to use or tolerate urban habitats, few studies have assessed the evolution of these anthropogenic phenotypes. The sole prior phylogenetic assessment of bats was restricted to species endemic to Africa and focused broadly on urban occupancy rather than use of anthropogenic structures. ^83^ We identified low-to-moderate phylogenetic signal in the anthropogenic roosting phenotype, indicating that the trait profile of anthropogenic roosting consists of particular clades that contain more bat species with high or low likelihood of having anthropogenic roosting ability.

Phylogenetic factorization did not identify clades with a high propensity to roost in anthropogenic structures, whereas the family Pteropodidae and subfamily Stenodermatinae were both identified as having a significantly lower probability. This finding is consistent with the results of our BRT analysis, where Pteropodidae showed a low but non-zero relative influence. Our BRT also found the family Vespertilionidae to have low-to-moderate influence in predicting anthropogenic roosting, although this clade was not identified by our phylogenetic analysis. Together, these results suggest that vespertilionid bats possess many traits that predispose bats to roost in anthropogenic structures (e.g., small size and insectivory) but such traits are not necessarily unique to this family. BiSSE analysis suggested the traits that predispose bats to be anthropogenic roosting may have impeded bat evolution, where anthropogenic roosting showed lower rates of speciation but equivalent extinction rates in comparison to natural roosting. This result is further supported by greater evolutionary success for natural roosting as evidenced by more lineages over time in comparison to the anthropogenic roosting phenotype and slightly lower transition rates from anthropogenic roosting to natural roosting than the reverse. These findings all together are in agreement with work that associates urbanization with loss of phylogenetic diversity globally in birds^84^ and emphasizes the need to preserve diversity in the face of increasing environmental change.

Taken together, our study demonstrates that anthropogenic roosting ability is predicted by both ecological and evolutionary characteristics of bat species. Suggesting that anthropogenic roosting bats may suffer loss in phylogenetic diversity and protecting roosting sites in anthropogenic structures may promote survival of vulnerable bat species. Our analyses also show important knowledge gaps in the ecology and the anthropogenic impacts of evolution in bats.

Although our novel dataset provides a standardized baseline in bat behavior to observe changing patterns in anthropogenic roosting throughout time, the number of species for which roosting ecology is unknown (*n* = 237) highlights the lack of available ecological data and emphasizes the need for further research to improve data coverage. For example, our evolutionary analysis of anthropogenic phenotypes showed an inconsistency in transition rate between models using known roosting ecology data and pseudoabsence data. Having roosting ecology information for these unknown species could uncover undetected phylogenetic patterns and improve understanding of transition rates between natural and anthropogenic phenotypes over time.

In addition to further expanding ecological data, an interesting future direction to consider is quantifying differences in reproductive outcomes between anthropogenic roosting bats and exclusively natural roosting bats as our trait based analysis revealed reduced reproductive outcome in anthropogenic roosting bats. Such comparative analyses could further elucidate anthropogenic impacts on population dynamics and inform conservation efforts towards species for which populations may not recover well under increasing human disturbance or conflict. Another logical next step is understanding co-roosting ecology of anthropogenic roosting bats in comparison to their natural roosting counterparts. Geographic overlap due to the presence of human-made structures could allow for species interactions that were not observed before and possibly unique to anthropogenic structures. Lastly, a comprehensive dataset documenting those species that directly live alongside humans and their potential interactions could be a powerful tool for improving predictions of pathogen transmission among species when paired with host-pathogen association data. For example, living alongside humans has been associated with a greater probability of being a reservoir of zoonotic pathogens in rodents ^85^. Similar work is needed for anthropogenic roosting in bats to improve pathogen surveillance and conflict mitigation relating to risk of transmission.

## Limitations

The data used in this study is a reflection of what is available in the current literature. As it is computationally expensive to calculate the number of records of anthropogenic roosting for each bat species, some bat classifications are single accounts and many are summaries in historical records which do not describe the frequency in which they were observed in a structure. Additionally, not all sources of historical literature were in consensus about anthropogenic roosting. Although this did not occur often, this was settled by searching for any additional information available and the primary sources used by the reference, if provided. If looking further into the literature still proved inconclusive, the species was assigned NA distinction. As roost ecology data is sparse, including quantitative metrics relating to the size of the roost is not available for the majority of bat species beyond the categorization of solitary or group roosting, but would be important for understanding the reliance on anthropogenic structures as roosts.

## Author Contributions

Conceptualization, D.B., B.B.; Methodology, D.B. and B.B.; Formal Analysis, B.B.; Investigation, B.B. and D.B.; Data Curation, B.B. and D.B.; Writing–Original Draft, B.B. and D.B., Writing –Review & Editing; B.B., D.B., L.M. and N.G.; Supervision, D.B.

## Acknowledgements

B.B. and D.J.B. were supported by funding to the Viral Emergence Research Initiative (VERENA) Institute, including NSF BII 2021909 and NSF BII 2213854.

## Methods

### Anthropogenic Roosting Data

We collected roosting ecology data for 1,279 extant bat species listed in the most recent mammal phylogeny,^68^ representing most of the over 1,460 recognized species in the order Chiroptera.^67^ Owing to taxonomic revision, we reclassified species in the genus *Miniopterus* from the *Vespertilionidae* to be the sole members of the family Miniopteridae.^86^ Starting on August 25, 2021, data were compiled through a manual search of existing literature and historical records using sources such as Google Scholar, IUCN, Animal Diversity Web, and Walkers *Bats of the World* ^107^ to identify roosting status and human-made structure type (building, home, bridge, ect.) of each bat species.

Anthropogenic roosting status was recorded as a binary variable indicating if a given bat species has been recorded in any human-made structures or roosting exclusively in natural roosting structures. Classification for each species can be found in Table S3 and corresponding references to sources used to verify roosting structure are provided in the supplementary information. A code of 1 indicates the bat species roosts in structures such as mines, buildings, homes, bridges, tunnels, roofs, and attics. Bat species that roost exclusively in natural structures (palm leaves, caves, tree hollows, foliage, etc.) were coded as 0. Species without roosting information or relevant citations (*n* = 237) were coded as pseudoabsences. We coded species as pseudoabsences in the absence of roosting information in the existing literature, having relatively few search returns (e.g., as occurs for poorly studied species), or not having relevant papers with roosting information within the first 10 pages of Google Scholar when searching with the following string: “Latin binomial’” AND “roost”.^87^ If a bat genus had consistent roosting ecology (i.e., all members used either anthropogenic or natural roosts), we updated any bats of that genus with missing values to the binary status of the group. However, if there were mixed values in a single genus specifically or the majority of species were considered NA, anthropogenic roosting remained NA until assigned as a pseudoabsence.

### Bat sampling effort and ecological traits

To account for additional bias introduced into the dataset through unequal sampling of bat species in our downstream analyses, we conducted a PubMed search with the *easyPubMed* package in R to calculate a citation count for all 1,279 bat species names.^88–91^ We next assembled a trait matrix of life history, morphological, geographic, and taxonomic traits to predict anthropogenic roosting. A description of name reconciliation across datasets can be found in the supplementary information. Trait data came primarily from COMBINE with additional geographic range variables from PanTHERIA.^62,63^ Biogeographical realms were separated into binary covariates to observe the importance of presence in each realm individually. IUCN conservation status and population trends were extracted for all available bat species with the *rredlist* package.^64^ In addition, binary covariates for all 18 bat families were generated to account for taxonomic relationships and to investigate whether a given family is disproportionately associated with anthropogenic roosting.^91–93^ We calculated several descriptive statistics using these data, including data coverage and the proportion of species roosting in anthropogenic structures as stratified by bat family, IUCN status, and biogeographical realm.

### Boosted regression trees

We used BRT models, a machine learning algorithm that determines variables of importance based on an ensemble of regression trees or classification trees, to identify the trait profiles of anthropogenic roosting bats. BRTs overcome many challenges associated with analyzing ecological data via traditional regression methods (e.g., non-independent data, non-randomly missing covariates, nonlinear patterns, complex interactions), making this approach ideal for analyzing bat trait data.^66^ After removing predictors with no variance or missing values for over 30% of bat species, our BRTs included a total 61 predictor variables (Table S4). Models were trained with the *gbm* R package, with anthropogenic roosting as a Bernoulli distributed response variable.^94^ We did not perform out-of-sample predictions, as the roosting data is well represented across the bat taxonomy, and our intention was to identify the trait profile of anthropogenic roosting bats rather than prediction.^95^ To mitigate the probability of overfitting, a grid search was used for hyperparameter tuning to determine the optimal values for the number of trees, learning rate (i.e., shrinkage), and interaction depth prior to fitting models. We identified an optimal learning rate of 0.0005, interaction depth of 4, and a maximum number of trees of 20,000 (Fig. S4). We derived several metrics of model performance with the *ROCR* R package (AUC, sensitivity, and specificity).^96^ Additionally, all models were ten-fold cross validated. We determined the influence of each predictor on roost status by variable importance, the proportion of improvement in model fit that is attributed to each predictor. We also created partial dependence plots to visualize the marginal effects of individual predictor variables on roost status while controlling for all other predictors.We also assessed the influence of pseudoabsences on BRT performance.

### Phylogenetic analyses

We used the above bat phylogeny for analyses of phylogenetic signal in anthropogenic roosting and the evolution of this trait across bats. In all analyses, we focused on known anthropogenic roosting status and assessed sensitivity of results after including pseudoabsences (*n* = 237).

We first used the *caper* package to calculate *D* in binary anthropogenic roosting, where a value of 1 indicates a phylogenetically random trait distribution and a value of 0 indicates phylogenetic clustering under a Brownian motion model of evolution.^97^ Significant departure from either model was quantified with a randomization test (1,000 permutations). However, because traits may also arise under a punctuated equilibrium model of evolution, we also used a graph-partitioning algorithm, phylogenetic factorization, to flexibly identify clades with different propensity to roost in anthropogenic structures at various taxonomic depths.^69^ Phylogenetic factorization partitions a phylogeny by iteratively identifying edges in a tree that maximize an objective function contrasting species separated by each edge; in this case, we used generalized linear models (binary anthropogenic roosting status) that also adjusted for citation counts.^98^ We used the *phylofactor* package for this analysis and determined the number of phylogenetic factors to retain using Holm’s sequentially rejective 5% cut-off for the family-wise error rate.^99^

To reconstruct the evolution of anthropogenic roosting across the bat phylogeny, we first used the *diversitree* package to fit candidate BiSSE models of trait-dependent diversification using maximum likelihood.^100^ We used Akaike’s information criterion (AIC)^101^ and Akaike weights (*w_i_*) to compare among a full BiSSE model (where speciation, extinction, and transition rates can vary between natural and anthropogenic roosting states; λ_0_ ≠ λ_1_, μ_0_ ≠ μ_1_, *q*_01_ ≠ *q*_10_) and eight constrained models, including an all-rates-different model with trait-independent speciation and/or extinction (λ_0_ = λ_1_ and/or μ_0_ = μ_1_, *q*_01_ ≠ *q*_10_), an irreversible model (λ_0_ ≠ λ_1_, μ_0_ ≠ μ_1_, *q*_10_ = 0) and variants with trait-independent speciation and/or extinction (λ_0_ = λ_1_ and/or μ_0_ = μ_1_, *q*_10_ = 0), and an equal rates model (λ_0_ = λ_1_, μ_0_ = μ_1_, *q*_01_ = *q*_10_). We then used the most competitive BiSSE model (i.e., ΔAIC < 2 and parsimony) and a Markov chain Monte Carlo (MCMC) sampling approach to estimate the posterior distribution of each parameter.^102^ MCMC used an exponential prior, maximum likelihood estimates from BiSSE for starting values, and slice sampling, with the tuning parameter *w* set as the range of each BiSSE parameter via an initial chain of 100 steps. Final chains were run for 2,000 steps with the first 20% discarded as burn-in. We used the *posterior* and *ggdist* packages to summarize resulting posterior distributions. The posterior means of transition rates were lastly used to perform stochastic character mapping (*n* = 1,000 simulations) over the bat phylogeny with the *phytools* package.^103,104^ We fixed the basal node to originate with natural roosting, given that the earliest bat fossils predate *Homo sapiens* by over 50 millions years.^105^ Stochastic character maps were used to reconstruct natural and anthropogenic roosting lineages through time and to show mean posterior probabilities of anthropogenic roosting across the bat phylogeny.^106^

