## Supplemental Information for "Ecological and Evolutionary Characteristics of Anthropogenic Roosting Ability in Bats of the World"

**Taxonomic reconciliation of trait datasets**

We manually matched select bat species across trait datasets to be consistent with the most recent phylogeny used for the collection of roosting ecology data (Upham et al. 2019). As the trait datasets were published in different years, many species were reverted to previous genus designations or updated to current names when necessary. For the PanTHERIA dataset, this included updating *Dermanura* from their former *Artibeus* classifications, switching *Hypsugo* species to *Pipistrellus*, and specific species of *Triaenops* to *Paratriaenops* (i.e., *Triaenops auritus* to *Paratriaenops auritus*, *Triaenops furculus* to *Paratriaenops furculus*). Species that were recently reclassified to another genus were also updated in the PanTHERIA binomial names to match our phylogenetic backbone (i.e., *Murina grisea* to *Hapoila grisea*, *Lissonycteris angolensis* to *Myonycteris angolensis*, *Eptesicus matroka* to *Neoromicia matroka, Hsunycteris thomasi* to *Lonchophylla thomasi*). Similar genus changes were made to COMBINE and IUCN binomial names to match with the phylogenetic backbone used in this analysis, apart from the switch to *Dermanura* to *Artibeus,* and species that needed to be changed to older names as these datasets reflect post 2019 taxonomic changes. This includes switching *Paremballonura* back to *Emballonura* (i.e., *Paremballonura atrata* to *Emballonura atrata*, *Paremballonura tiavato* to *Emballonura tiavato*), several *Macronycteris* species to *Hipposideros* (i.e., *M. commersoni*, *M. gigas*, *M. thomensis*, *M. vittatus*), *Lonchophylla* to *Hsunycteris* (i.e., *L. cadenai*, *L. pattoni*), *Gardnerycteris* to *Mimon* (i.e., *G. crenulatum*, *G. koepckeae*), *Baedon* to *Rhogeessa* (i.e., *B. alleni*, *B. gracillis*), *Austronomus* to *Tadarida* (i.e., *A. australis*, *A. Kuboriensis*), *Vampyriscus* to *Vampyressa* (i.e., *V. bidens*, *V. brocki*, *V. nymphea*), and changes to select species including: *Scotonycteris ophiodon* to *Casinycteris ophiodon*, *Pteropus leucopterus* to *Desmalopex leucopterus*, *Rhyneptesicus nasutus* to *Eptesicus nasutus*, *Hypsugo affinis* to *Falsistrellus affinis*, *Lyroderma lyra* to *Megaderma lyra*, *Ozimops loriae* to *Mormopterus loriae*, *Mormopterus kalinowskii* to *Nyctinomops kalinowskii*, *Perimyotis subflavus* to *Pipistrellus subflavus*, *Boneia bidens* to *Rousettus bidens*, *Chaerephon jobimena* to *Tadarida jobimena*. Additionally, species that were present in our dataset but were previously a subspecies, or closely related, to those in the trait datasets were synonymized. Across all trait datasets, the following species were considered to be synonyms: *Dermanura incomitatus* and *Dermanura watsoni*, *Harpiocephalus mordax* and *Harpiocephalus harpia*, *Hsunycteris thomasi* and *Lonchophylla thomasi*, *Lophostoma aequatorialis* to *Lophostoma occidentalis*, *Lophostoma yasuni* and *Lophostoma carrikeri*, *Miniopterus fuliginosus* and *Miniopterus schreibersii*, *Molossus barnesi* and *Molossus coibensis*, *Myotis flavus* and *Myotis formosus*, *Myotis midastactus* and *Myotis simus*, *Natalus saturatus* and *Natalus stramineus*, *Paracoelops megalotis* and *Hipposideros Pomona*, *Pipistrellus deserti* and *Pipistrellus kuhlii*, *Pteropus argentatus* and *Pteropus chrysoproctus*, *Pteropus yapensis* and *Pteropus pelewensis*, *Triaenops menamena* and *Triaenops rufus*, *Rhinolophus chaseni* and *Rhinolophus borneensis*, *Triaenops rufus* and *Triaenops persicus*, *Myotis aelleni* and *Myotis chiloensis, Myotis hajastanicus* and *Myotis aurascens*. For the COMBINE and IUCN trait datasets, a few additional species were synonymized (*Natalus* *lanatus* and *Natalus mexicanus*, *Nyctophilus timoriensis* and *Nyctophilus corbeni*). Additionally, there were synonyms specific to the PanTHERIA dataset which were: *Carollia sowelli* and *Carollia brevicauda*, *Natalus mexicanus* and *Natalus stramineus, Myotis abei* and *Myotis petax", Myotis ricketti* and *Myotis pilosus, Pteropus insularis* and *Pteropus pelagicus, Pteropus insularis* and *Pteropus pelagicus*, *Sturnira thomasi* and *Sturnira angeli*). Minor discrepancies in names were also corrected as needed throughout datasets (i.e., *Anoura carishina* to *Anoura canishina*, *Chiroderma vizzotoi* to *Chiroderma vizottoi*, *Dermanura azteca* to *Dermanura aztecus*, *Dermanura cinerea* to *Dermanura cinereus*, *Dermanura glauca* to *Dermanura glaucus*, *Dermanura gnoma* to *Dermanura gnomus*, *Dermanura rosenbergi* to *Dermanura rosenbergii*, *Dermanura tolteca* to *Dermanura toltecus*, *Diclidurus isabella* to *Diclidurus isabellus*, *Murina lorelieae* to *Murina loreliae*, *Neoromicia brunneus* to *Neoromicia brunnea*, *Neoromicia nanus* to *Neoromicia nana*, *Neoromicia somalicus* to *Neoromicia somalica*)

**Optimizing parameters for Boosted Regression Tree Models**

For the main text model of 1,042 bat species with known anthropogenic classification data and the pseudoabsence model where unknown species are assumed to be natural roosting ($n$ = 1,279), a grid search was conducted to select optimal parameters. We selected three interaction depths (2, 3, 4) and three learning rates (0.01, 0.001, 0.0005). In combination, we tested these interaction depts and learning rates with an assortment of initial trees (5000, 10000, 15000, 20000, 25000). Combinations of low number initial trees (i.e., 5000 and 1000) and small learning rates (0.0005) were removed, resulting in 39 parameterizations for each model. These parameters were run through BRTs with the *gbm* package (Greenwell et al. 2020). To assess performance, metrics were derived with the *ROCR* package (AUC, sensitivity, and specificity) (Sing et al. 2005) and compared to select the parameters for the final models. Combinations with the highest performance values were considered. Visualizations of performance metrics are shown in Figure S4. For the pseudoabsence model, we identified an optimal learning rate of 0.001, interaction depth of 4, and a maximum number of trees of 25000. This option had the second highest AUC while optimizing sensitivity, at a very small cost to specificity.

**Figures**

**Figure S1. Relative importance comparisons of BRT models with and without pseudoabsences.** (A) Relative importance of the initial model colored black and pseudoabsence model colored orange. (B) The correlation relative importance between BRT models.

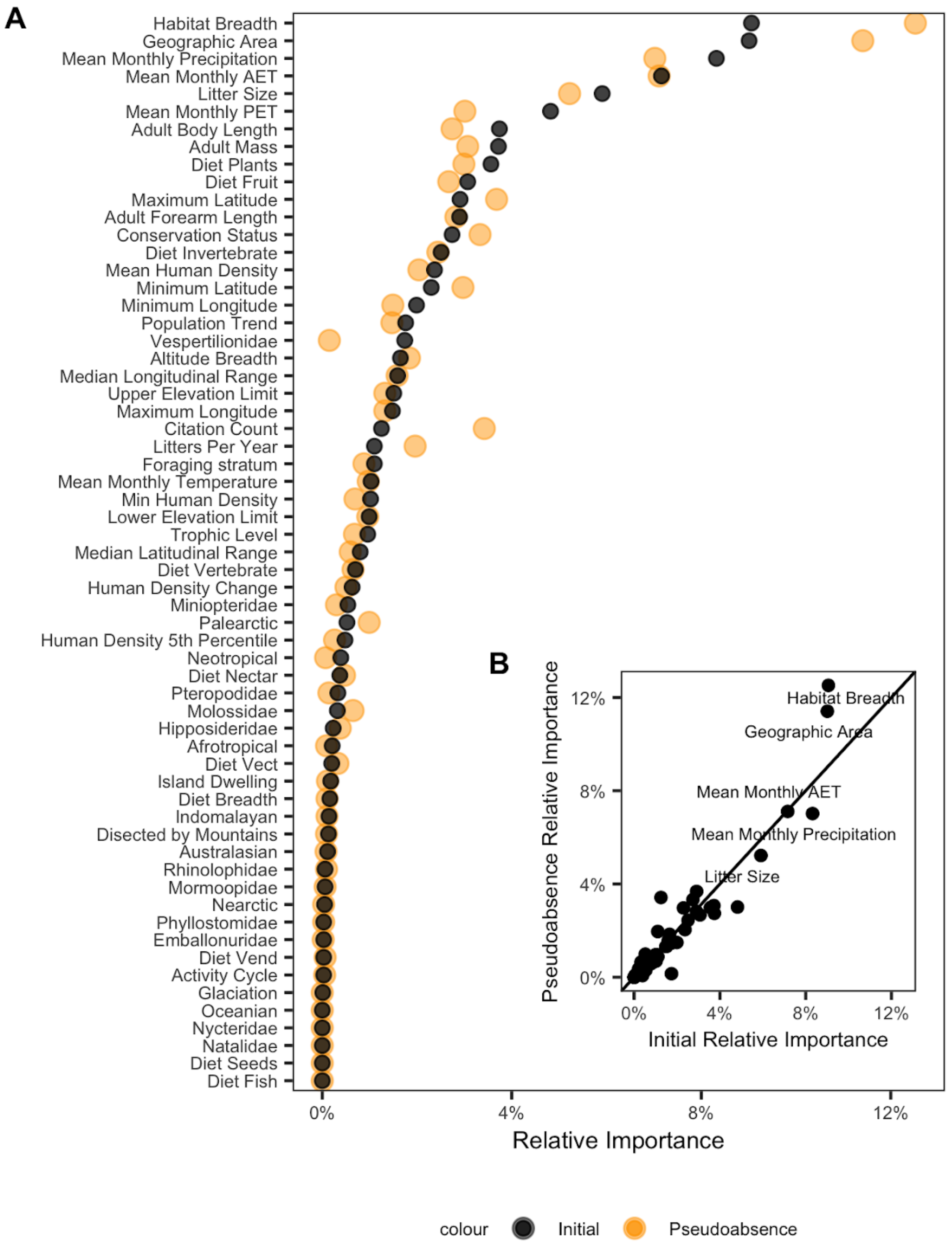

**Figure S2. Partial dependence plots of the top 15 predictors ordered by relative importance for pseudoabsecnce model (*n* = 1,279).** The black line displays the marginal effect of a given variable for predicting roosting status. Histograms and point colors show the distribution of the continuous and categorical predictors, respectively.

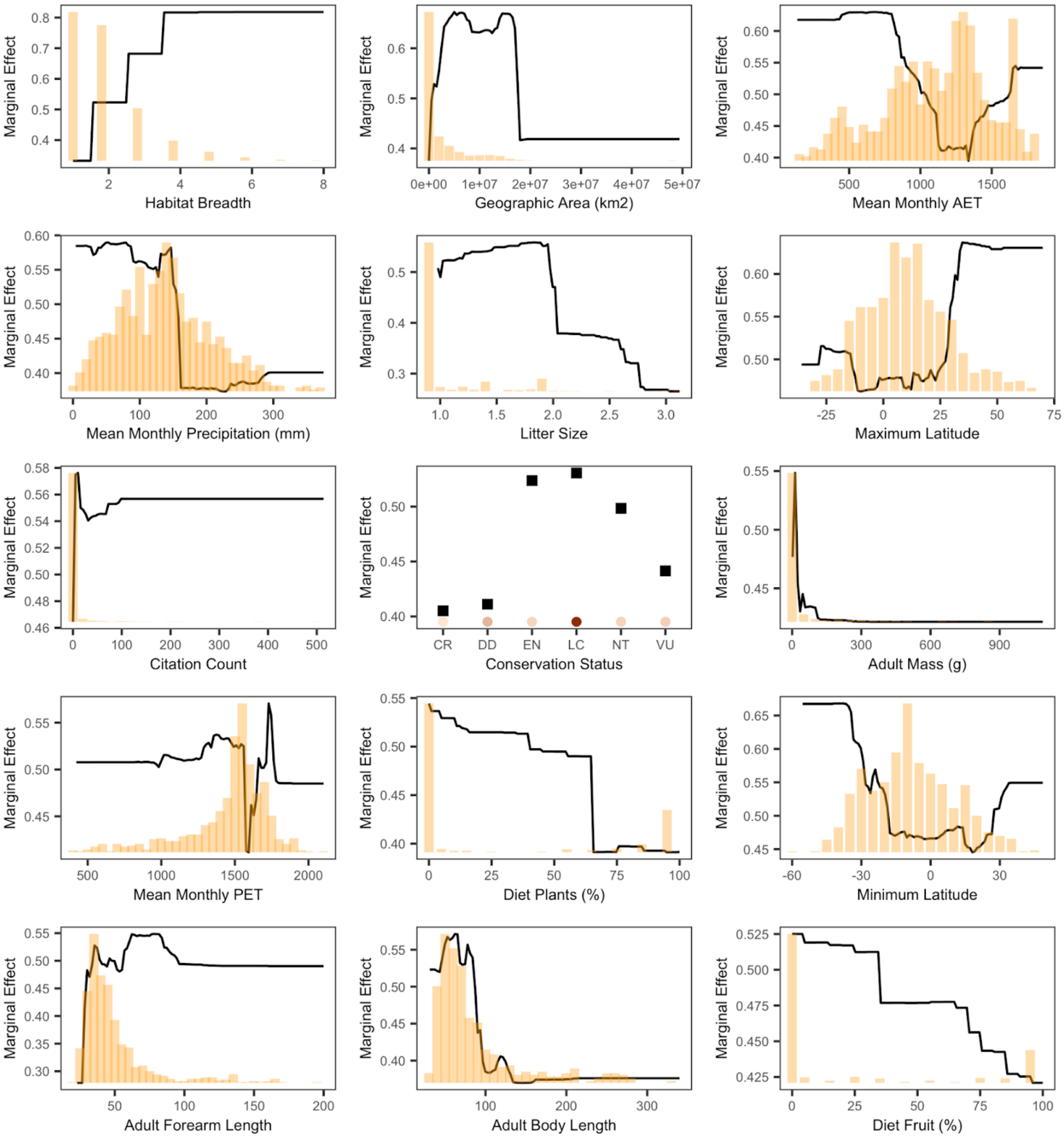

**Figure S3. Parameters from the top BiSSE model when including pseudoabsences (*n* = 1,279 species).** Shown are posterior medians, 66% and 95% credible intervals, and posterior densities from MCMC sampling.

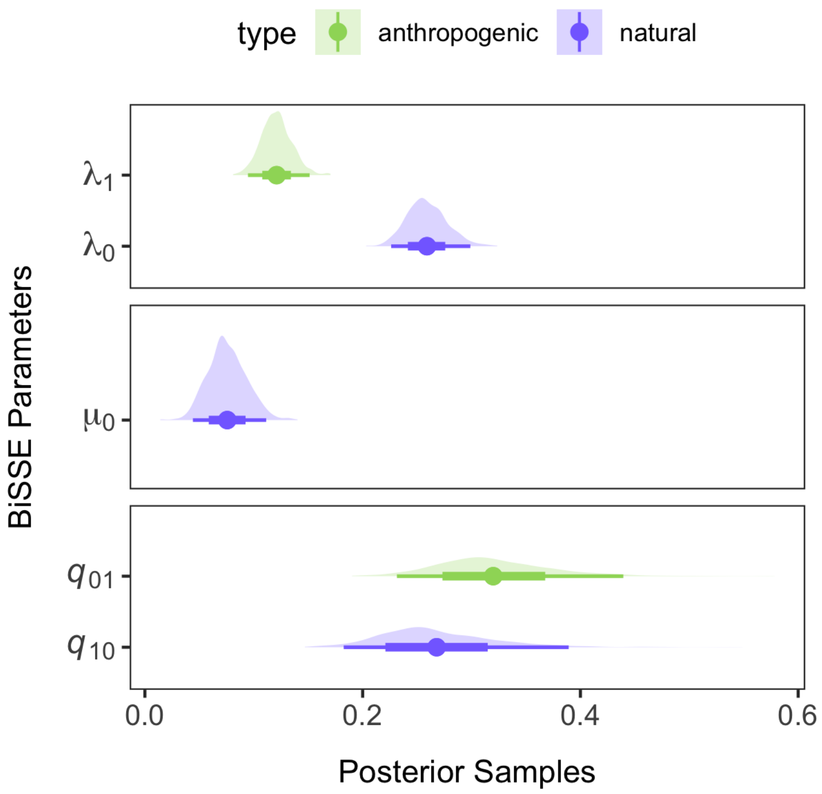

**Figure S4. Performance of parameters in the tuning grid for BRT models.** Plotted is AUC by learning rate and number of trees for both models, colored by interaction depth.

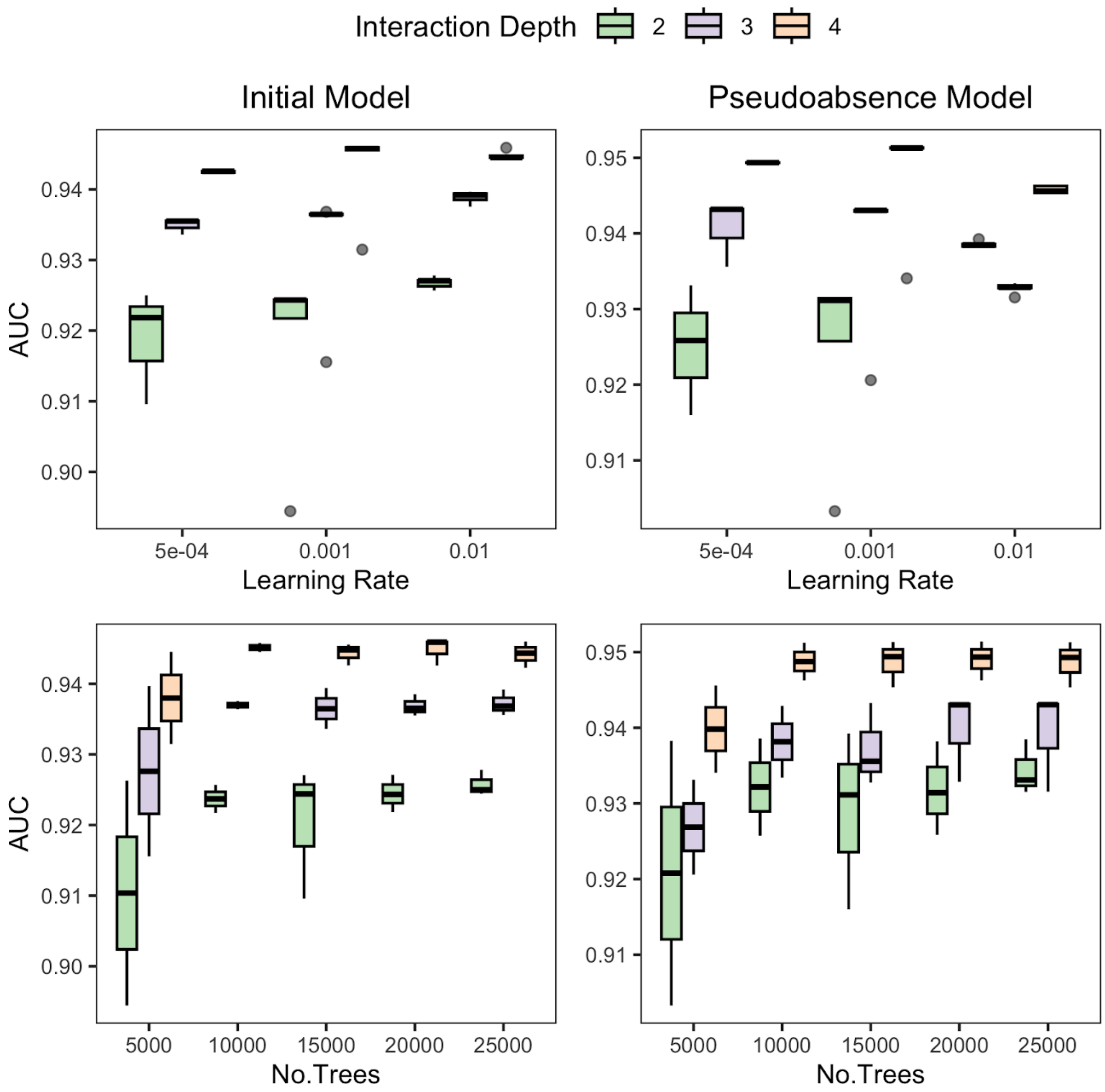

**Tables**

**Table S1. Phylogenetic factorization of anthropogenic roosting status for the primary analysis (*i*, *n* = 1,042) and the dataset with pseudoabsences (*ii*, *n* = 1,279).** The table shows the number of retained clades after a 5% family-wise error rate, taxa corresponding to those clades, number of species per clade, and mean predicted probabilities for the clade compared to the paraphyletic remainder.

| Model | Factor | Taxa | Tips | Clade | Other |
|---|---|---|---|---|---|
| (*i*) | 1 | Pteropodidae | 157 | 0.09 | 0.56 |
| | 2 | *Ectophylla, Ardops, Phyllops, Stenoderma, Pygoderma, Sphaeronycteris, Centurio, Artibeus, Dermanura, Enchisthenes, Vampyressa, Mesophylla, Vampyrodes, Platyrrhinus, Chiroderma, Uroderma, Sturnira* | 63 | 0.25 | 0.50 |
| (*ii*) | 1 | Pteropodidae | 190 | 0.07 | 0.45 |
| | 2 | *Ectophylla, Ariteus, Ardops, Phyllops, Stenoderma, Pygoderma, Ametrida, Sphaeronycteris, Centurio, Artibeus, Dermanura, Enchisthenes, Vampyressa, Mesophylla, Vampyrodes, Platyrrhinus, Chiroderma, Uroderma, Sturnira* | 92 | 0.17 | 0.42 |

**Table S2. Comparison among eight BiSSE models of trait-dependent diversification, including speciation (λ), extinction (μ), and transitions between binary states (*q*) for the pseudoabsence dataset (*n* = 1,279).** Models are ranked by difference in AIC from the top model (ΔAIC), the number of estimated parameters (*k*), and Akaike weights ($w_i$).

| **Model** | **ΔAIC** | ***k*** | **$w_i$** |
|---|---|---|---|
| $\lambda_0 \neq \lambda_1$, $\mu_0 = \mu_1$, $q_{01} \neq q_{10}$ | 0.00 | 5 | 0.51 |
| $\lambda_0 \neq \lambda_1$, $\mu_0 \neq \mu_1$, $q_{01} \neq q_{10}$ | 0.80 | 6 | 0.34 |
| $\lambda_0 = \lambda_1$, $\mu_0 = \mu_1$, $q_{01} \neq q_{10}$ | 3.26 | 4 | 0.10 |
| $\lambda_0 = \lambda_1$, $\mu_0 \neq \mu_1$, $q_{01} \neq q_{10}$ | 4.90 | 5 | 0.04 |
| $\lambda_0 = \lambda_1$, $\mu_0 = \mu_1$, $q_{01} = q_{10}$ | 50.12 | 3 | <0.01 |
| $\lambda_0 \neq \lambda_1$, $\mu_0 \neq \mu_1$, $q_{10} = 0$ | 115.26 | 5 | <0.01 |
| $\lambda_0 = \lambda_1$, $\mu_0 \neq \mu_1$, $q_{10} = 0$ | 136.97 | 4 | <0.01 |
| $\lambda_0 \neq \lambda_1$, $\mu_0 = \mu_1$, $q_{10} = 0$ | 296.39 | 4 | <0.01 |
| $\lambda_0 = \lambda_1$, $\mu_0 = \mu_1$, $q_{10} = 0$ | 593.64 | 3 | <0.01 |

**Table S3. Bat species roosting status (n = 1042) accompanied by corresponding references.**

| Species | Family | Status | Type of human-made structure | Reference |
|---|---|---|---|---|
| Acerodon celebensis | Pteropodidae | 0 | NA | 1 |
| Acerodon humilis | Pteropodidae | 0 | NA | 1 |
| Acerodon jubatus | Pteropodidae | 0 | NA | 1 |
| Acerodon leucotis | Pteropodidae | 0 | NA | 1 |
| Acerodon mackloti | Pteropodidae | 0 | NA | 1 |
| Aethalops aequalis | Pteropodidae | 0 | NA | 2 |
| Amorphochilus schnablii | Furipteridae | 1 | abandoned buildings | 1 |
| Anoura canishina | Phyllostomidae | 0 | NA | 1 |
| Anoura caudifer | Phyllostomidae | 0 | NA | 1 |
| Anoura cultrata | Phyllostomidae | 1 | tunnels | 1,3 |
| Anoura fistulata | Phyllostomidae | 0 | NA | 4 |
| Anoura geoffroyi | Phyllostomidae | 0 | NA | 1 |
| Anoura latidens | Phyllostomidae | 0 | NA | 1,5 |
| Anoura luismanueli | Phyllostomidae | 0 | NA | 6 |
| Antrozous pallidus | Vespertilionidae | 1 | mines, buildings, bridges | 1,7,8 |
| Aproteles bulmerae | Pteropodidae | 0 | NA | 9,10 |
| Ardops nichollsi | Phyllostomidae | 0 | NA | 1 |
| Arielulus societatis | Vespertilionidae | 0 | NA | 11 |
| Artibeus amplus | Phyllostomidae | 0 | NA | 12 |
| Artibeus fimbriatus | Phyllostomidae | 0 | NA | 13 |
| Artibeus fraterculus | Phyllostomidae | 1 | bridges, churches, houses, mines | 14 |
| Artibeus hirsutus | Phyllostomidae | 1 | abandoned mines, buildings | 15 |
| Artibeus inopinatus | Phyllostomidae | 1 | unoccupied houses | 15 |
| Artibeus jamaicensis | Phyllostomidae | 1 | houses and buildings | 1 |
| Artibeus lituratus | Phyllostomidae | 1 | buildings | 16 |
| Artibeus obscurus | Phyllostomidae | 0 | NA | 17,18 |
| Artibeus planirostris | Phyllostomidae | 0 | NA | 19 |
| Asellia arabica | Hipposideridae | 0 | NA | 20 |
| Asellia patrizii | Hipposideridae | 1 | buildings | 21,22 |
| Asellia tridens | Hipposideridae | 1 | temples, mines, open-wells, underground irrigation tunnels and old tombs and buildings | 23 |
| Aselliscus stoliczkanus | Hipposideridae | 0 | NA | 1 |
| Aselliscus tricuspidatus | Hipposideridae | 1 | Tunnels | 24 |
| Balantiopteryx infusca | Emballonuridae | 1 | culverts | 25 |
| Balantiopteryx io | Emballonuridae | 0 | NA | 26 |
| Balantiopteryx plicata | Emballonuridae | 1 | buildings and bridges | 27 |

| | | | | |
|---|---|---|---|---|
| Balionycteris maculata | Pteropodidae | 0 | NA | 1 |
| Barbastella barbastellus | Vespertilionidae | 1 | buildings | 1 |
| Barbastella beijingensis | Vespertilionidae | 1 | old buildings, tunnel | 1,28 |
| Barbastella leucomelas | Vespertilionidae | 1 | old buildings, mines, tunnels | 1 |
| Bauerus dubiaquercus | Vespertilionidae | 1 | building | 29 |
| Brachyphylla cavernarum | Phyllostomidae | 1 | building and well | 1,30 |
| Brachyphylla nana | Phyllostomidae | 0 | NA | 31,32 |
| Cardioderma cor | Megadermatidae | 1 | abandoned buildings, culvert | 33,34 |
| Carollia brevicauda | Phyllostomidae | 1 | concrete bridge | 35 |
| Carollia castanea | Phyllostomidae | 0 | NA | 36,37 |
| Carollia manu | Phyllostomidae | 1 | tunnels, road culverts, buildings | 38, 1 |
| Carollia perspicillata | Phyllostomidae | 1 | tunnels, road culverts, buildings | 39, 1 |
| Carollia sowelli | Phyllostomidae | 1 | houses | 35, 40, 41 |
| Carollia subrufa | Phyllostomidae | 1 | empty wells, culverts, hollow trees and buildings | 42, 1 |
| Casinycteris argynnis | Pteropodidae | 0 | NA | 1 |
| Casinycteris ophiodon | Pteropodidae | 0 | NA | 43 |
| Centronycteris centralis | Emballonuridae | 0 | NA | 45 |
| Centronycteris maximiliani | Emballonuridae | 0 | NA | 46 |
| Centurio senex | Phyllostomidae | 0 | NA | 1 |
| Chaerephon aloysiisabaudiae | Molossidae | 0 | NA | 47 |
| Chaerephon ansorgei | Molossidae | 1 | abandoned mines, roofs of buildings, expansion joints of buildings | 48 |
| Chaerephon atsinanana | Molossidae | 1 | roofs of houses, schools, and churches | 49, 50 |
| Chaerephon bemmeleni | Molossidae | 0 | NA | 51, 41 |
| Chaerephon bivittatus | Molossidae | 1 | old mines | 52 |
| Chaerephon bregullae | Molossidae | 0 | NA | 53, 54 |
| Chaerephon chapini | Molossidae | 1 | houses | 55 |
| Chaerephon gallagheri | Molossidae | 0 | NA | 56, 1 |
| Chaerephon jobensis | Molossidae | 1 | buildings, bridges and jetties | 57, 58 |
| Chaerephon major | Molossidae | 1 | houses | 59, 1 |
| Chaerephon nigeriae | Molossidae | 1 | roofs and eaves of house, bungalows, | 60, 1 |
| Chaerephon plicatus | Molossidae | 1 | disused buildings and temples | 61 |
| Chaerephon pumilus | Molossidae | 1 | thatched and corrugated iron roofs | 62 |
| Chaerephon russatus | Molossidae | 0 | NA | 63 |
| Chaerephon solomonis | Molossidae | 0 | NA | 64 |
| Chalinolobus dwyeri | Vespertilionidae | 1 | Mine tunnels | 65 |

| | | | | |
|---|---|---|---|---|
| Chalinolobus gouldii | Vespertilionidae | 1 | buildings and bat boxes | 66, 67, 1 |
| Chalinolobus morio | Vespertilionidae | 1 | schoolhouse, church | 68, 69, 70 |
| Chalinolobus neocaledonicus | Vespertilionidae | 1 | roofs | 71, 1 |
| Chalinolobus nigrogriseus | Vespertilionidae | 1 | buildings | 72, 1 |
| Chalinolobus picatus | Vespertilionidae | 1 | abandoned buildings | 73, 1 |
| Chalinolobus tuberculatus | Vespertilionidae | 0 | NA | 74, 75 |
| Cheiromeles parvidens | Molossidae | 0 | NA | 76, 1 |
| Cheiromeles torquatus | Molossidae | 1 | buildings | 77 |
| Chilonatalus micropus | Natalidae | 0 | NA | 78 |
| Chilonatalus tumidifrons | Natalidae | 0 | NA | 78 |
| Chiroderma salvini | Phyllostomidae | 1 | houses | 79 |
| Chiroderma trinitatum | Phyllostomidae | 0 | NA | 80 |
| Chiroderma villosum | Phyllostomidae | 1 | buildings, room in building | 81, 82, 80 |
| Chironax melanocephalus | Pteropodidae | 0 | NA | 83 |
| Choeroniscus minor | Phyllostomidae | 0 | NA | 84 |
| Choeronycteris mexicana | Phyllostomidae | 1 | buildings,tunnels | 1 |
| Chrotopterus auritus | Phyllostomidae | 1 | mines, abandoned buildings, mayan ruins | 85 |
| Cistugo lesueuri | Vespertilionidae | 1 | under concrete bridge | 86 |
| Cloeotis percivali | Hipposideridae | 1 | mine tunnels, dam body corridors | 87 |
| Coelops frithii | Hipposideridae | 1 | man-made tunnel, abandoned mining cave | 88 |
| Coelops hirsutus | Hipposideridae | 0 | NA | 89 |
| Coelops robinsoni | Hipposideridae | 0 | NA | 89 |
| Coleura afra | Emballonuridae | 1 | house, cellars | 90, 1 |
| Coleura kibomalandy | Emballonuridae | 0 | NA | 91 |
| Coleura seychellensis | Emballonuridae | 0 | NA | 92 |
| Cormura brevirostris | Emballonuridae | 1 | concrete bridges | 93 |
| Corynorhinus mexicanus | Vespertilionidae | 1 | abandoned buildings, mine shaft | 94, 95 |
| Corynorhinus rafinesquii | Vespertilionidae | 1 | copper mine, empty boiler, wells, attics, bridges | 96 |
| Corynorhinus townsendii | Vespertilionidae | 1 | abandoned buildings, abandoned mines | 97, 96 |
| Craseonycteris thonglongyai | Craseonycteridae | 0 | NA | 98 |
| Cynomops abrasus | Molossidae | 0 | NA | 99 |
| Cynomops greenhalli | Molossidae | 1 | buildings | 100 |
| Cynomops mexicanus | Molossidae | 1 | buildings | 101 |
| Cynomops planirostris | Molossidae | 1 | buildings | 102 |
| Cynopterus brachyotis | Pteropodidae | 0 | NA | 103, 1 |
| Cynopterus horsfieldii | Pteropodidae | 0 | NA | 104, 1 |

| | | | | |
|---|---|---|---|---|
| Cynopterus sphinx | Pteropodidae | 1 | under the eaves of houses, buildings | 105, 1 |
| Cynopterus titthaecheilus | Pteropodidae | 1 | old building | 106 |
| Cyttarops alecto | Emballonuridae | 0 | NA | 107 |
| Dermanura anderseni | Phyllostomidae | 0 | NA | 108 |
| Dermanura aztecus | Phyllostomidae | 1 | abandoned mine | 109 |
| Dermanura bogotensis | Phyllostomidae | 0 | NA | 110 |
| Dermanura cinereus | Phyllostomidae | 0 | NA | 41, 108 |
| Dermanura glaucus | Phyllostomidae | 0 | NA | 111, 108 |
| Dermanura gnomus | Phyllostomidae | 0 | NA | 108 |
| Dermanura incomitatus | Phyllostomidae | 0 | NA | 115 |
| Dermanura phaeotis | Phyllostomidae | 0 | NA | 112 |
| Dermanura rosenbergii | Phyllostomidae | 0 | NA | 113 |
| Dermanura toltecus | Phyllostomidae | 1 | buildings | 114 |
| Dermanura watsoni | Phyllostomidae | 0 | NA | 115 |
| Desmalopex leucopterus | Pteropodidae | 0 | NA | 116 |
| Desmodus rotundus | Phyllostomidae | 0 | NA | 117 |
| Diaemus youngi | Phyllostomidae | 0 | NA | 118 |
| Diclidurus albus | Emballonuridae | 0 | NA | 119 |
| Diclidurus ingens | Emballonuridae | 1 | ceiling of house | 120 |
| Diclidurus scutatus | Emballonuridae | 0 | NA | 121 |
| Diphylla ecaudata | Phyllostomidae | 1 | mines, houses | 122, 1 |
| Dobsonia anderseni | Pteropodidae | 1 | man-made tunnels | 123, 1 |
| Dobsonia beauforti | Pteropodidae | 0 | NA | 83, 1 |
| Dobsonia chapmani | Pteropodidae | 0 | NA | 83, 1 |
| Dobsonia crenulata | Pteropodidae | 0 | NA | 83, 1 |
| Dobsonia emersa | Pteropodidae | 0 | NA | 83, 1 |
| Dobsonia exoleta | Pteropodidae | 0 | NA | 83, 1 |
| Dobsonia inermis | Pteropodidae | 0 | NA | 83, 1 |
| Dobsonia minor | Pteropodidae | 0 | NA | 83, 1 |
| Dobsonia moluccensis | Pteropodidae | 1 | sinkholes, boulder piles, old mines, disused buildings | 124 |
| Dobsonia pannietensis | Pteropodidae | 0 | NA | 125, 1 |
| Dobsonia peronii | Pteropodidae | 0 | NA | 126, 1 |
| Dobsonia praedatrix | Pteropodidae | 0 | NA | 127, 1 |
| Dobsonia viridis | Pteropodidae | 0 | NA | 128, 1 |
| Dryadonycteris capixaba | Phyllostomidae | 0 | NA | 1 |
| Dyacopterus brooksi | Pteropodidae | 0 | NA | 1 |
| Dyacopterus rickarti | Pteropodidae | 0 | NA | 1 |

| | | | | |
|---|---|---|---|---|
| Dyacopterus spadiceus | Pteropodidae | 0 | NA | 129, 1 |
| Ectophylla alba | Phyllostomidae | 0 | NA | 130 |
| Eidolon dupreanum | Pteropodidae | 0 | NA | 131 |
| Eidolon helvum | Pteropodidae | 0 | NA | 132, 133 |
| Emballonura alecto | Emballonuridae | 1 | human-made tunnels | 134, 135 |
| Emballonura atrata | Emballonuridae | 0 | NA | 136 |
| Emballonura beccarii | Emballonuridae | 0 | NA | 137 |
| Emballonura dianae | Emballonuridae | 0 | NA | 138 |
| Emballonura furax | Emballonuridae | 1 | mining tunnels | 139 |
| Emballonura monticola | Emballonuridae | 1 | tables and buttresses | 140 |
| Emballonura raffrayana | Emballonuridae | 0 | NA | 141 |
| Emballonura semicaudata | Emballonuridae | 0 | NA | 142 |
| Emballonura serii | Emballonuridae | 0 | NA | 143 |
| Emballonura tiavato | Emballonuridae | 0 | NA | 144 |
| Enchisthenes hartii | Phyllostomidae | 0 | NA | 145 |
| Eonycteris major | Pteropodidae | 0 | NA | 1 |
| Eonycteris robusta | Pteropodidae | 0 | NA | 1 |
| Eonycteris spelaea | Pteropodidae | 0 | NA | 1 |
| Epomophorus crypturus | Pteropodidae | 0 | NA | 83 |
| Epomophorus gambianus | Pteropodidae | 0 | NA | 146 |
| Epomophorus labiatus | Pteropodidae | 0 | NA | 83 |
| Epomophorus wahlbergi | Pteropodidae | 1 | Thatch of open sheds | 83 |
| Epomops buettikoferi | Pteropodidae | 0 | NA | 1 |
| Epomops dobsonii | Pteropodidae | 0 | NA | 1 |
| Epomops franqueti | Pteropodidae | 0 | NA | 83 |
| Eptesicus bottae | Vespertilionidae | 1 | buildlings, tombs, and ruins | 147, 148 |
| Eptesicus brasiliensis | Vespertilionidae | 1 | houses | 149 |
| Eptesicus furinalis | Vespertilionidae | 1 | within walls or floors, behind window shutters, between beams of porch | 150 |
| Eptesicus fuscus | Vespertilionidae | 1 | walls, boxed-in eaves, barnes, attics, houses, churches | 1, 151 |
| Eptesicus gobiensis | Vespertilionidae | 1 | buildings, ancient salt mines | 152, 153 |
| Eptesicus guadeloupensis | Vespertilionidae | 0 | NA | 154 |
| Eptesicus hottentotus | Vespertilionidae | 1 | mines, buildings | 154, 132 |
| Eptesicus innoxius | Vespertilionidae | 1 | attics and roofs of uninhabited buildings | 155, 154 |
| Eptesicus isabellinus | Vespertilionidae | 1 | bridges, buildings | 154 |
| Eptesicus japonensis | Vespertilionidae | 1 | buildings | 154 |
| Eptesicus lobatus | Vespertilionidae | 1 | window sills, window frames, and ceilings | 156 |

| | | | | |
|---|---|---|---|---|
| Eptesicus nasutus | Vespertilionidae | 1 | crevices of walls and behind stones of ruined buildings | 157 |
| Eptesicus nilssonii | Vespertilionidae | 1 | houses, cellar | 158 |
| Eptesicus pachyomus | Vespertilionidae | 1 | buildings | 159, 160 |
| Eptesicus pachyotis | Vespertilionidae | 0 | NA | 161, 154 |
| Eptesicus serotinus | Vespertilionidae | 1 | attics, voids behind sheathing and shutters, ventilation shafts of buildings or joints in bridges | 162 |
| Erophylla bombifrons | Phyllostomidae | 0 | NA | 1 |
| Erophylla sezekorni | Phyllostomidae | 0 | NA | 1 |
| Euderma maculatum | Vespertilionidae | 0 | NA | 154 |
| Eudiscoderma thongareeae | Vespertilionidae | 1 | house | 163 |
| Eudiscopus denticulus | Vespertilionidae | 0 | NA | 164 |
| Eumops auripendulus | Molossidae | 1 | corrugated iron roof, attic | 165 |
| Eumops bonariensis | Molossidae | 1 | iron roof, under bridges | 99 |
| Eumops chiribaya | Molossidae | 1 | buildings | 1 |
| Eumops dabbenei | Molossidae | 1 | house | 166 |
| Eumops ferox | Molossidae | 1 | under roof shingles | 167 |
| Eumops floridanus | Molossidae | 1 | under shingles of spanish-like tile | 96 |
| Eumops glaucinus | Molossidae | 1 | under shingles of spanish-like tile | 168 |
| Eumops hansae | Molossidae | 0 | NA | 169 |
| Eumops maurus | Molossidae | 1 | 15 story apartment | 170 |
| Eumops patagonicus | Molossidae | 1 | buildings | 1, 99 |
| Eumops perotis | Molossidae | 1 | buildings | 171 |
| Eumops trumbulli | Molossidae | 1 | buildings, roofs | 99 |
| Eumops underwoodi | Molossidae | 0 | NA | 96 |
| Eumops wilsoni | Molossidae | 1 | church | 99 |
| Falsistrellus affinis | Vespertilionidae | 1 | roofs of buildings | 172 |
| Falsistrellus mackenziei | Vespertilionidae | 0 | NA | 154 |
| Falsistrellus petersi | Vespertilionidae | 1 | Eaves of house | 154 |
| Falsistrellus tasmaniensis | Vespertilionidae | 1 | buildings | 154 |
| Furipterus horrens | Furipteridae | 0 | NA | 1, 173 |
| Glauconycteris alboguttata | Vespertilionidae | 0 | NA | 1 |
| Glauconycteris argentata | Vespertilionidae | 0 | NA | 154 |
| Glauconycteris beatrix | Vespertilionidae | 0 | NA | 174 |
| Glauconycteris egeria | Vespertilionidae | 0 | NA | 1 |
| Glauconycteris gleni | Vespertilionidae | 0 | NA | 1 |
| Glauconycteris humeralis | Vespertilionidae | 0 | NA | 175 |
| Glauconycteris kenyacola | Vespertilionidae | 0 | NA | 1 |

| | | | | |
|---|---|---|---|---|
| Glauconycteris poensis | Vespertilionidae | 0 | NA | 176 |
| Glauconycteris superba | Vespertilionidae | 0 | NA | 1 |
| Glauconycteris variegata | Vespertilionidae | 1 | thatched roofs of abandoned huts | 177 |
| Glischropus aquilus | Vespertilionidae | 0 | NA | 1 |
| Glischropus bucephalus | Vespertilionidae | 0 | NA | 1, 154 |
| Glischropus javanus | Vespertilionidae | 0 | NA | 1 |
| Glischropus tylopus | Vespertilionidae | 0 | NA | 1 |
| Glossophaga commissarisi | Phyllostomidae | 1 | houses | 41, 1 |
| Glossophaga leachii | Phyllostomidae | 1 | buildings and culverts | 41, 1 |
| Glossophaga longirostris | Phyllostomidae | 1 | buildings, houses, culverts, tunnels | 41, 178 |
| Glossophaga morenoi | Phyllostomidae | 1 | culverts, wells and buildings | 41 |
| Glossophaga soricina | Phyllostomidae | 1 | tunnels, abandoned mines, culverts, underbridges, buildings | 179 |
| Glyphonycteris daviesi | Phyllostomidae | 0 | NA | 41 |
| Glyphonycteris sylvestris | Phyllostomidae | 0 | NA | 41 |
| Haplonycteris fischeri | Pteropodidae | 0 | NA | 180 |
| Harpiocephalus harpia | Vespertilionidae | 0 | NA | 1 |
| Harpiocephalus mordax | Vespertilionidae | 0 | NA | 1 |
| Harpyionycteris whiteheadi | Pteropodidae | 0 | NA | 83 |
| Hesperoptenus blanfordi | Vespertilionidae | 0 | NA | 1 |
| Hesperoptenus doriae | Vespertilionidae | 0 | NA | 1, 181 |
| Hesperoptenus gaskelli | Vespertilionidae | 0 | NA | 1 |
| Hesperoptenus tickelli | Vespertilionidae | 0 | NA | 182 |
| Hesperoptenus tomesi | Vespertilionidae | 0 | NA | 1 |
| Hipposideros abae | Hipposideridae | 0 | NA | 183 |
| Hipposideros alongensis | Hipposideridae | 0 | NA | 184 |
| Hipposideros armiger | Hipposideridae | 1 | lofts of houses, verandahs of old houses, old temples | 185 |
| Hipposideros ater | Hipposideridae | 1 | buildings, mines, tunnels, culverts, wells | 186 |
| Hipposideros beatus | Hipposideridae | 1 | road culverts | 187 |
| Hipposideros bicolor | Hipposideridae | 0 | NA | 188 |
| Hipposideros breviceps | Hipposideridae | 0 | NA | 189 |
| Hipposideros caffer | Hipposideridae | 1 | buildings | 190 |
| Hipposideros calcaratus | Hipposideridae | 1 | tunnels | 191 |
| Hipposideros cervinus | Hipposideridae | 1 | abandoned mines | 192 |
| Hipposideros cineraceus | Hipposideridae | 0 | NA | 193 |
| Hipposideros commersoni | Hipposideridae | 0 | NA | 194 |
| Hipposideros coronatus | Hipposideridae | 0 | NA | 194 |
| Hipposideros corynophyllus | Hipposideridae | 0 | NA | 194 |

| | | | | |
|---|---|---|---|---|
| Hipposideros coxi | Hipposideridae | 0 | NA | 194 |
| Hipposideros curtus | Hipposideridae | 1 | house | 195 |
| Hipposideros cyclops | Hipposideridae | 1 | house, church belfry, disused mine | 196 |
| Hipposideros demissus | Hipposideridae | 0 | NA | 194 |
| Hipposideros diadema | Hipposideridae | 1 | buildings and underground chambers | 197 |
| Hipposideros dinops | Hipposideridae | 0 | NA | 194 |
| Hipposideros durgadasi | Hipposideridae | 0 | NA | 198 |
| Hipposideros dyacorum | Hipposideridae | 0 | NA | 194 |
| Hipposideros edwardshilli | Hipposideridae | 0 | NA | 194 |
| Hipposideros einnaythu | Hipposideridae | 1 | house roof | 194 |
| Hipposideros fuliginosus | Hipposideridae | 0 | NA | 194 |
| Hipposideros fulvus | Hipposideridae | 1 | old buildings, cellars, forts, old wells | 199 |
| Hipposideros galeritus | Hipposideridae | 1 | old mines, culverts and crevices in old buildings, caves, among large boulders, overhanging ledges, tunnels, dungeons, forts, temples and churches | 200 |
| Hipposideros gigas | Hipposideridae | 0 | NA | 194 |
| Hipposideros griffini | Hipposideridae | 0 | old mines | 194 |
| Hipposideros halophyllus | Hipposideridae | 0 | NA | 194 |
| Hipposideros hypophyllus | Hipposideridae | 0 | NA | 194 |
| Hipposideros inexpectatus | Hipposideridae | 0 | NA | 201 |
| Hipposideros inornatus | Hipposideridae | 1 | disused mines | 194 |
| Hipposideros jonesi | Hipposideridae | 1 | disused mines | 194 |
| Hipposideros lamottei | Hipposideridae | 1 | mine adits | 194 |
| Hipposideros lankadiva | Hipposideridae | 1 | old disused tunnels, old temples, old forts, dark deep channels under dam sites and cellars under old buildings | 202 |
| Hipposideros larvatus | Hipposideridae | 1 | mineshafts, pagodas, and buildings | 203 |
| Hipposideros lekaguli | Hipposideridae | 0 | NA | 194 |
| Hipposideros lylei | Hipposideridae | 0 | NA | 194 |
| Hipposideros macrobullatus | Hipposideridae | 0 | NA | 194 |
| Hipposideros madurae | Hipposideridae | 0 | NA | 194 |
| Hipposideros maggietaylorae | Hipposideridae | 1 | mines, tunnels | 194 |
| Hipposideros marisae | Hipposideridae | 1 | disused mine adits | 194 |
| Hipposideros megalotis | Hipposideridae | 1 | house | 204 |
| Hipposideros muscinus | Hipposideridae | 0 | NA | 194 |
| Hipposideros obscurus | Hipposideridae | 1 | mine shaft | 194 |
| Hipposideros papua | Hipposideridae | 0 | NA | 194 |
| Hipposideros pelingensis | Hipposideridae | 0 | NA | 194 |
| Hipposideros pendelburyi | Hipposideridae | 0 | NA | 194 |

| | | | | |
|---|---|---|---|---|
| Hipposideros pomona | Hipposideridae | 1 | disused wells | 205 |
| Hipposideros pratti | Hipposideridae | 0 | NA | 194 |
| Hipposideros pygmaeus | Hipposideridae | 0 | NA | 194 |
| Hipposideros ridleyi | Hipposideridae | 1 | drainage pipes, old houses | 194 |
| Hipposideros rotalis | Hipposideridae | 0 | NA | 194 |
| Hipposideros ruber | Hipposideridae | 1 | old mines, culverts under roads, abandoned houses | 194 |
| Hipposideros scutinares | Hipposideridae | 0 | NA | 194 |
| Hipposideros semoni | Hipposideridae | 1 | mines, old buildings, culverts under roads | 194 |
| Hipposideros sorenseni | Hipposideridae | 0 | NA | 194 |
| Hipposideros speoris | Hipposideridae | 1 | underground cellars, old forts, palaces, under bridges, old disused buildings, temples, tunnels | 194 |
| Hipposideros stenotis | Hipposideridae | 1 | road culverts, old mines | 206 |
| Hipposideros sumbae | Hipposideridae | 1 | roofs of houses | 194 |
| Hipposideros thomensis | Hipposideridae | 1 | old water mine | 207 |
| Hipposideros turpis | Hipposideridae | 1 | abandoned mines and bomb shelters | 194 |
| Hipposideros vittatus | Hipposideridae | 1 | under the eaves of buildings | 132 |
| Hipposideros wollastoni | Hipposideridae | 0 | NA | 194 |
| Histiotus laephotis | Vespertilionidae | 1 | wooden storage barn | 154 |
| Histiotus macrotus | Vespertilionidae | 1 | mines, houses | 1, 154 |
| Histiotus magellanicus | Vespertilionidae | 0 | NA | 208 |
| Histiotus montanus | Vespertilionidae | 1 | houses | 1, 209 |
| Histiotus velatus | Vespertilionidae | 1 | churches and roofs | 1, 154 |
| Hsunycteris thomasi | Phyllostomidae | 0 | NA | 210 |
| Hylonycteris underwoodi | Phyllostomidae | 1 | tunnel | 1 |
| Hypsignathus monstrosus | Pteropodidae | 0 | NA | 1 |
| Hypsugo anchietae | Vespertilionidae | 0 | NA | 211 |
| Hypsugo dolichodon | Vespertilionidae | 0 | NA | 154, 212 |
| Ia io | Vespertilionidae | 0 | NA | 1, 213 |
| Idionycteris phyllotis | Vespertilionidae | 1 | mine shaft | 154 |
| Kerivoula africana | Vespertilionidae | 0 | NA | 214 |
| Kerivoula argentata | Vespertilionidae | 1 | under eaves of huts, on walls sheltered by eaves of a rondavel | 154 |
| Kerivoula cuprosa | Vespertilionidae | 0 | NA | 154 |
| Kerivoula hardwickii | Vespertilionidae | 1 | houses | 154 |
| Kerivoula lanosa | Vespertilionidae | 0 | NA | 154 |
| Kerivoula muscina | Vespertilionidae | 0 | NA | 154 |
| Kerivoula pellucida | Vespertilionidae | 0 | NA | 154, 215 |
| Kerivoula picta | Vespertilionidae | 1 | huts and buildings | 1 |

| | | | | |
|---|---|---|---|---|
| Kerivoula whiteheadi | Vespertilionidae | 0 | NA | 154 |
| Laephotis botswanae | Vespertilionidae | 0 | NA | 154 |
| Laephotis namibensis | Vespertilionidae | 0 | NA | 154 |
| Laephotis wintoni | Vespertilionidae | 0 | NA | 216 |
| Lampronycteris brachyotis | Phyllostomidae | 1 | mines and old buildings | 217 |
| Lasionycteris noctivagans | Vespertilionidae | 1 | buildings, under loose boards of buildings | 1 |
| Lasiurus atratus | Vespertilionidae | 0 | NA | 218 |
| Lasiurus blossevillii | Vespertilionidae | 1 | buildings | 154 |
| Lasiurus borealis | Vespertilionidae | 0 | NA | 154 |
| Lasiurus cinereus | Vespertilionidae | 1 | overhangs of buildings | 219 |
| Lasiurus degelidus | Vespertilionidae | 0 | NA | 220 |
| Lasiurus ega | Vespertilionidae | 1 | crevices in buildings | 154 |
| Lasiurus insularis | Vespertilionidae | 0 | NA | 221 |
| Lasiurus intermedius | Vespertilionidae | 0 | NA | 222 |
| Lasiurus minor | Vespertilionidae | 0 | NA | 223 |
| Lasiurus pfeifferi | Vespertilionidae | 0 | NA | 154 |
| Lasiurus seminolus | Vespertilionidae | 0 | NA | 132 |
| Lasiurus varius | Vespertilionidae | 0 | NA | 224 |
| Lasiurus xanthinus | Vespertilionidae | 0 | NA | 225, 226 |
| Latidens salimalii | Pteropodidae | 0 | NA | 83 |
| Lavia frons | Megadermatidae | 1 | buildings | 1 |
| Leptonycteris curasoae | Phyllostomidae | 0 | mine shafts | 227, 228 |
| Leptonycteris nivalis | Phyllostomidae | 1 | mines, culverts, unoccupied buildings | 96 |
| Leptonycteris yerbabuenae | Phyllostomidae | 0 | culverts, mines, buildings | 83 |
| Lichonycteris obscura | Phyllostomidae | 0 | NA | 229 |
| Lionycteris spurrelli | Phyllostomidae | 0 | culvert | 41 |
| Lonchophylla bokermanni | Phyllostomidae | 1 | mines | 230 |
| Lonchophylla concava | Phyllostomidae | 1 | tunnels | 41 |
| Lonchophylla dekeyseri | Phyllostomidae | 1 | tunnels | 41 |
| Lonchophylla fornicata | Phyllostomidae | 1 | railroad tunnel | 231 |
| Lonchophylla handleyi | Phyllostomidae | 0 | NA | 41 |
| Lonchophylla hesperia | Phyllostomidae | 1 | house | 41 |
| Lonchophylla mordax | Phyllostomidae | 0 | NA | 41 |
| Lonchophylla orienticollina | Phyllostomidae | 0 | NA | 41 |
| Lonchophylla robusta | Phyllostomidae | 1 | cave-like anthropogenic structures | 41 |
| Lonchophylla thomasi | Phyllostomidae | 0 | NA | 210 |
| Lonchorhina aurita | Phyllostomidae | 1 | mine tunnels | 41 |

| | | | | |
|---|---|---|---|---|
| Lonchorhina fernandezi | Phyllostomidae | 0 | NA | 210 |
| Lonchorhina inusitata | Phyllostomidae | 0 | NA | 232 |
| Lonchorhina marinkellei | Phyllostomidae | 0 | NA | 233 |
| Lonchorhina orinocensis | Phyllostomidae | 0 | NA | 41 |
| Lophostoma aequatorialis | Phyllostomidae | 0 | NA | 41 |
| Lophostoma brasiliense | Phyllostomidae | 0 | NA | 41 |
| Lophostoma carrikeri | Phyllostomidae | 0 | NA | 41 |
| Lophostoma evotis | Phyllostomidae | 1 | Building | 234 |
| Lophostoma kalkoae | Phyllostomidae | 0 | NA | 235 |
| Lophostoma silvicolum | Phyllostomidae | 0 | NA | 236 |
| Lophostoma yasuni | Phyllostomidae | 0 | NA | 41 |
| Macroderma gigas | Megadermatidae | 1 | old mines | 237 |
| Macroglossus minimus | Pteropodidae | 1 | under roofs | 83 |
| Macroglossus sobrinus | Pteropodidae | 1 | roofs | 83 |
| Macrophyllum macrophyllum | Phyllostomidae | 1 | wet tunnels, under bridges, and abandoned buildings | 238 |
| Macrotus californicus | Phyllostomidae | 1 | abandoned mines | 96 |
| Macrotus waterhousii | Phyllostomidae | 1 | mine tunnels and old buildings | 239 |
| Megaderma lyra | Megadermatidae | 1 | temple | 240, 1 |
| Megaderma spasma | Megadermatidae | 1 | buildings | 241, 1 |
| Megaerops kusnotoi | Pteropodidae | 0 | NA | 242 |
| Megaerops wetmorei | Pteropodidae | 0 | NA | 243 |
| Megaloglossus azagnyi | Pteropodidae | 1 | inside human habitations | 83 |
| Megaloglossus woermanni | Pteropodidae | 1 | native hut | 1 |
| Melonycteris fardoulisi | Pteropodidae | 0 | NA | 244 |
| Melonycteris melanops | Pteropodidae | 0 | NA | 83 |
| Melonycteris woodfordi | Pteropodidae | 0 | NA | 83 |
| Mesophylla macconnelli | Phyllostomidae | 0 | NA | 41 |
| Micronycteris brosseti | Phyllostomidae | 0 | NA | 41 |
| Micronycteris buriri | Phyllostomidae | 0 | NA | 41 |
| Micronycteris hirsuta | Phyllostomidae | 1 | buildings and under bridges | 41 |
| Micronycteris megalotis | Phyllostomidae | 1 | houses and under bridges | 41 |
| Micronycteris microtis | Phyllostomidae | 1 | mines, buildings, and culverts | 41 |
| Micronycteris minuta | Phyllostomidae | 1 | mines | 246 |
| Micronycteris sanborni | Phyllostomidae | 0 | NA | 41 |
| Micronycteris schmidtorum | Phyllostomidae | 1 | human constructions | 247 |
| Micropteropus intermedius | Pteropodidae | 0 | NA | 83, 1 |

| | | | | |
|---|---|---|---|---|
| Micropteropus pusillus | Pteropodidae | 0 | NA | 248, 1 |
| Mimetillus moloneyi | Vespertilionidae | 1 | houses | 154, 1 |
| Mimon bennettii | Phyllostomidae | 1 | highway culverts | 1, 249 |
| Mimon cozumelae | Phyllostomidae | 1 | mines, culverts | 41, 1 |
| Mimon crenulatum | Phyllostomidae | 0 | abandoned buildings | 41, 1 |
| Miniopterus aelleni | Miniopteridae | 0 | NA | 250, 1 |
| Miniopterus ambohitrensis | Miniopteridae | 0 | NA | 251, 1 |
| Miniopterus australis | Miniopteridae | 1 | abandoned mines, tunnels, water drains, buildings | 252, 1 |
| Miniopterus brachytragos | Miniopteridae | 0 | NA | 253, 1 |
| Miniopterus egeri | Miniopteridae | 0 | NA | 254, 1 |
| Miniopterus fraterculus | Miniopteridae | 1 | railroad tunnels and disused mines | 132, 1 |
| Miniopterus fuliginosus | Miniopteridae | 0 | abandoned mines | 255, 256 |
| Miniopterus fuscus | Miniopteridae | 1 | mines | 257, 1 |
| Miniopterus gleni | Miniopteridae | 0 | NA | 258, 1 |
| Miniopterus griffithsi | Miniopteridae | 0 | NA | 259, 1 |
| Miniopterus griveaudi | Miniopteridae | 0 | NA | 260, 1 |
| Miniopterus inflatus | Miniopteridae | 1 | mines | 132, 1 |
| Miniopterus macrocneme | Miniopteridae | 0 | NA | 261, 1 |
| Miniopterus maghrebensis | Miniopteridae | 0 | NA | 262, 1 |
| Miniopterus magnater | Miniopteridae | 0 | NA | 263,1 |
| Miniopterus mahafaliensis | Miniopteridae | 0 | NA | 264, 1 |
| Miniopterus majori | Miniopteridae | 0 | NA | 265, 1 |
| Miniopterus manavi | Miniopteridae | 0 | NA | 1 |
| Miniopterus medius | Miniopteridae | 0 | NA | 266, 1 |
| Miniopterus minor | Miniopteridae | 0 | NA | 267, 1 |
| Miniopterus mossambicus | Miniopteridae | 1 | old mine | 268, 1 |
| Miniopterus natalensis | Miniopteridae | 1 | disused mines | 289, 1 |
| Miniopterus newtoni | Miniopteridae | 0 | tunnels, mines | 270, 1 |
| Miniopterus paululus | Miniopteridae | 0 | NA | 271, 1 |
| Miniopterus petersoni | Miniopteridae | 0 | NA | 272, 1 |
| Miniopterus pusillus | Miniopteridae | 1 | culverts | 273, 1 |
| Miniopterus robustior | Miniopteridae | 0 | NA | 274, 1 |
| Miniopterus schreibersii | Miniopteridae | 1 | mines and tunnels | 275, 1 |
| Miniopterus sororculus | Miniopteridae | 1 | building | 276, 1 |
| Miniopterus tristis | Miniopteridae | 0 | NA | 277, 1 |
| Molossops aequatorianus | Molossidae | 0 | NA | 278 |

| | | | | |
|---|---|---|---|---|
| Molossops mattogrossensis | Molossidae | 0 | NA | 99 |
| Molossops neglectus | Molossidae | 1 | hanging on the wall of a house | 279 |
| Molossops temminckii | Molossidae | 1 | Attics and roofs made of overlapping palm logs | 280, 281 |
| Molossus aztecus | Molossidae | 1 | houses | 99 |
| Molossus barnesi | Molossidae | 1 | roofs | 282 |
| Molossus coibensis | Molossidae | 1 | roofs | 282 |
| Molossus currentium | Molossidae | 1 | building, 2nd story porch | 283, 99 |
| Molossus molossus | Molossidae | 1 | culverts, tunnels, bridges, attics | 96 |
| Molossus pretiosus | Molossidae | 1 | buildings, roof dwellings, church roof | 99, 284 |
| Molossus rufus | Molossidae | 1 | buildings | 99 |
| Molossus sinaloae | Molossidae | 1 | houses | 285 |
| Monophyllus plethodon | Phyllostomidae | 0 | NA | 41, 1 |
| Monophyllus redmani | Phyllostomidae | 0 | NA | 41, 1 |
| Mops bakarii | Molossidae | 1 | buildings, attics, hospital building | 99, 1 |
| Mops brachypterus | Molossidae | 1 | buildings | 99, 1 |
| Mops condylurus | Molossidae | 1 | attics, joints of bridges | 99, 1 |
| Mops congicus | Molossidae | 0 | NA | 99, 1 |
| Mops demonstrator | Molossidae | 0 | NA | 99, 1 |
| Mops leucostigma | Molossidae | 1 | buildings | 99, 1 |
| Mops midas | Molossidae | 1 | attics, joints of concrete bridges, and between cement walls or bricks | 99, 1 |
| Mops mops | Molossidae | 0 | NA | 99, 1 |
| Mops nanulus | Molossidae | 1 | thatched houses and sheds | 99, 1 |
| Mops niangarae | Molossidae | 0 | NA | 99, 1 |
| Mops niveiventer | Molossidae | 1 | buildings | 99, 1 |
| Mops spurrelli | Molossidae | 0 | NA | 99 |
| Mops thersites | Molossidae | 1 | under roof, culverts, and drains | 99 |
| Mops trevori | Molossidae | 0 | NA | 99 |
| Mormoops blainvillei | Mormoopidae | 0 | NA | 286 |
| Mormoops megalophylla | Mormoopidae | 1 | mines, tunnels | 286, 1 |
| Mormopterus acetabulosus | Molossidae | 0 | NA | 99 |
| Mormopterus beccarii | Molossidae | 1 | houses | 99,1 |
| Mormopterus eleryi | Molossidae | 0 | NA | 99 |
| Mormopterus francoismoutoui | Molossidae | 1 | buildings | 287 |
| Mormopterus jugularis | Molossidae | 1 | buildings, metal or ceramic roof tiles, attic trusses | 288 |
| Mormopterus kitcheneri | Molossidae | 1 | buildings | 99 |

| | | | | |
|---|---|---|---|---|
| Mormopterus loriae | Molossidae | 0 | NA | 99 |
| Mormopterus minutus | Molossidae | 1 | urban buildings | 99 |
| Mormopterus norfolkensis | Molossidae | 1 | under metal caps of wooden power poles, buildings, roostboxes | 99 |
| Mormopterus phrudus | Molossidae | 0 | NA | 99 |
| Mormopterus planiceps | Molossidae | 1 | buildings | 99 |
| Mosia nigrescens | Emballonuridae | 1 | roofs of houses | 144 |
| Murina aenea | Vespertilionidae | 0 | NA | 289 |
| Murina aurata | Vespertilionidae | 0 | NA | 154 |
| Murina balaensis | Vespertilionidae | 0 | NA | 290 |
| Murina bicolor | Vespertilionidae | 1 | abandoned bunker | 154 |
| Murina cyclotis | Vespertilionidae | 0 | NA | 154 |
| Murina eleryi | Vespertilionidae | 0 | NA | 291 |
| Murina florium | Vespertilionidae | 1 | disused buildings | 154 |
| Murina hilgendorfi | Vespertilionidae | 1 | houses, abandoned tunnels, and bat boxes, flying squirrel nest boxes | 154 |
| Murina huttoni | Vespertilionidae | 0 | NA | 154 |
| Murina leucogaster | Vespertilionidae | 1 | houses | 154 |
| Murina rozendaali | Vespertilionidae | 0 | NA | 292 |
| Murina ryukyuana | Vespertilionidae | 0 | NA | 293 |
| Murina suilla | Vespertilionidae | 0 | NA | 154 |
| Murina tenebrosa | Vespertilionidae | 1 | abandoned mine | 154 |
| Murina tubinaris | Vespertilionidae | 0 | NA | 154 |
| Murina ussuriensis | Vespertilionidae | 0 | abandoned mines, buildings | 154 |
| Musonycteris harrisoni | Phyllostomidae | 1 | culvert | 154 |
| Myonycteris angolensis | Pteropodidae | 1 | old mine adits | 83 |
| Myonycteris torquata | Pteropodidae | 0 | NA | 83 |
| Myopterus daubentonii | Molossidae | 0 | NA | 99 |
| Myopterus whitleyi | Molossidae | 1 | roof of rubber packing shed | 99 |
| Myotis adversus | Vespertilionidae | 0 | railway tunnel | 154 |
| Myotis aelleni | Vespertilionidae | 1 | abandoned houses, abandoned mines, crevices in walls, attic spaces, or be found under tiles or other roofing materials | 324, 301 |
| Myotis albescens | Vespertilionidae | 1 | attics, under roofs, under palm logs, roofs of palm thatched huts, outer walls of buildings | 234 |
| Myotis alcathoe | Vespertilionidae | 0 | NA | 154 |
| Myotis altarium | Vespertilionidae | 0 | NA | 154 |
| Myotis atacamensis | Vespertilionidae | 1 | churches, old houses, small drainage tunnels | 154 |
| Myotis ater | Vespertilionidae | 1 | village houses | 154 |
| Myotis aurascens | Vespertilionidae | 1 | construction joints of bridges | 295, 296, 297 |

| | | | | |
|---|---|---|---|---|
| Myotis auriculus | Vespertilionidae | 1 | buildings and mines | 96 |
| Myotis austroriparius | Vespertilionidae | 1 | attics, barns, bridges, and mines | 154 |
| Myotis badius | Vespertilionidae | 0 | NA | 154 |
| Myotis bechsteinii | Vespertilionidae | 1 | buildings, bird and bat boxes | 154 |
| Myotis blythii | Vespertilionidae | 1 | attics, mines, and buildings | 154 |
| Myotis bocagii | Vespertilionidae | 0 | NA | 154 |
| Myotis bombinus | Vespertilionidae | 1 | abandoned mines, unused tunnels, buildings, and under bridges | 298 |
| Myotis brandtii | Vespertilionidae | 1 | voids and crevices in buildings, bat and bird boxes | 299 |
| Myotis bucharensis | Vespertilionidae | 0 | abandoned mines | 154 |
| Myotis californicus | Vespertilionidae | 1 | mines and buildings | 96 |
| Myotis capaccinii | Vespertilionidae | 0 | mines | 300 |
| Myotis chiloensis | Vespertilionidae | 1 | abandoned houses, abandoned mines, crevices in walls, attic spaces, or be found under tiles or other roofing materials | 301 |
| Myotis chinensis | Vespertilionidae | 0 | NA | 154 |
| Myotis ciliolabrum | Vespertilionidae | 1 | buildings, tunnels, mines | 96 |
| Myotis cobanensis | Vespertilionidae | 1 | cathedral | 302 |
| Myotis csorbai | Vespertilionidae | 0 | NA | 154 |
| Myotis dasycneme | Vespertilionidae | 1 | large attics and church steeples, | 154 |
| Myotis daubentonii | Vespertilionidae | 1 | buildings, bridges, bat boxes | 303 |
| Myotis davidii | Vespertilionidae | 1 | NA | 154, 297 |
| Myotis dieteri | Vespertilionidae | 0 | NA | 154 |
| Myotis dinellii | Vespertilionidae | 1 | abandoned house | 154 |
| Myotis dominicensis | Vespertilionidae | 0 | NA | 154 |
| Myotis emarginatus | Vespertilionidae | 1 | wine cellar, houses, and barns | 304, 305 |
| Myotis escalerai | Vespertilionidae | 1 | mines, tunnels, bridges | 154 |
| Myotis evotis | Vespertilionidae | 1 | Abandoned buildings, mines | 96 |
| Myotis fimbriatus | Vespertilionidae | 0 | NA | 154 |
| Myotis findleyi | Vespertilionidae | 0 | NA | 306 |
| Myotis flavus | Vespertilionidae | 1 | Abandoned mine | 307, 308 |
| Myotis formosus | Vespertilionidae | 1 | buildlings | 307, 308 |
| Myotis fortidens | Vespertilionidae | 1 | tunnels, abandoned constructions, and thatched roofs | 154 |
| Myotis frater | Vespertilionidae | 1 | bat boxes and tunnels | 309 |
| Myotis gomantongensis | Vespertilionidae | 0 | NA | 154 |
| Myotis goudoti | Vespertilionidae | 0 | NA | 154 |
| Myotis grisescens | Vespertilionidae | 1 | mines and buildings | 310 |
| Myotis hajastanicus | Vespertilionidae | 1 | NA | 311 |
| Myotis hasseltii | Vespertilionidae | 1 | abandoned building, meter box | 154 |

| | | | | |
|---|---|---|---|---|
| Myotis horsfieldii | Vespertilionidae | 1 | tunnels, bridges, and crevices in old buildings, and between wooden beams | 154 |
| Myotis ikonnikovi | Vespertilionidae | 1 | houses, under bridges, farm roofs | 154 |
| Myotis izecksohni | Vespertilionidae | 1 | Abandoned church | 154 |
| Myotis keaysi | Vespertilionidae | 1 | buildings and roofs of buildings | 212 |
| Myotis keenii | Vespertilionidae | 1 | bridges and house attics | 154 |
| Myotis laniger | Vespertilionidae | 0 | NA | 154 |
| Myotis lavali | Vespertilionidae | 0 | NA | 213 |
| Myotis leibii | Vespertilionidae | 1 | expansion joints of bridges | 96 |
| Myotis levis | Vespertilionidae | 1 | attic of chapel | 214 |
| Myotis longipes | Vespertilionidae | 1 | old disused buildings, tunnels, underground canals | 154 |
| Myotis lucifugus | Vespertilionidae | 1 | mines, attics of buildings | 96 |
| Myotis macrodactylus | Vespertilionidae | 1 | abandoned mine, tunnels, bomb shelters | 154 |
| Myotis macropus | Vespertilionidae | 1 | mines, tunnels, road culverts, storm drains | 154 |
| Myotis macrotarsus | Vespertilionidae | 0 | NA | 154 |
| Myotis martiniquensis | Vespertilionidae | 1 | concrete bridge | 154 |
| Myotis melanorhinus | Vespertilionidae | 1 | barns | 96 |
| Myotis midastactus | Vespertilionidae | 1 | thatched roof | 315 |
| Myotis moluccarum | Vespertilionidae | 1 | tunnels, mines, bridges | 154 |
| Myotis muricola | Vespertilionidae | 0 | NA | 154 |
| Myotis myotis | Vespertilionidae | 1 | churches, loft spaces, and castles | 154 |
| Myotis mystacinus | Vespertilionidae | 1 | buildings, bird and bat boxes | 316 |
| Myotis nattereri | Vespertilionidae | 1 | buildings, bird and bat boxes | 154 |
| Myotis nesopolus | Vespertilionidae | 0 | NA | 317 |
| Myotis nigricans | Vespertilionidae | 1 | buildings, mines | 154 |
| Myotis nipalensis | Vespertilionidae | 1 | cracks in buildings and old mines | 318 |
| Myotis occultus | Vespertilionidae | 1 | bridges, attics, and mines | 96 |
| Myotis oxyotus | Vespertilionidae | 0 | NA | 319 |
| Myotis peninsularis | Vespertilionidae | 1 | tunnels, sewers, abandoned buildings, and palm roofs of houses. | 154 |
| Myotis pequinius | Vespertilionidae | 1 | dwellings and temples | 320 |
| Myotis petax | Vespertilionidae | 1 | under bridges, mine tunnels, cellars, wells, attics, and steeples | 321 |
| Myotis pilosus | Vespertilionidae | 0 | NA | 322, 154 |
| Myotis planiceps | Vespertilionidae | 0 | NA | 323 |
| Myotis pruinosus | Vespertilionidae | 0 | NA | 154 |
| Myotis punicus | Vespertilionidae | 1 | WW2 shelter, tunnel | 325, 326 |
| Myotis ridleyi | Vespertilionidae | 1 | under house | 154 |
| Myotis riparius | Vespertilionidae | 1 | under roof | 327 |

| | | | | |
|---|---|---|---|---|
| Myotis rosseti | Vespertilionidae | 1 | house roofs | 154 |
| Myotis ruber | Vespertilionidae | 1 | house roofs | 154 |
| Myotis rufopictus | Vespertilionidae | 1 | recorded in room of building | 328 |
| Myotis schaubi | Vespertilionidae | 1 | buildings | 154 |
| Myotis secundus | Vespertilionidae | 1 | tunnels | 154 |
| Myotis septentrionalis | Vespertilionidae | 1 | buildings, mines | 154 |
| Myotis siligorensis | Vespertilionidae | 1 | crevices in old buildings | 154 |
| Myotis simus | Vespertilionidae | 1 | thatched roofs of houses | 329 |
| Myotis sodalis | Vespertilionidae | 1 | buildings and bridges | 154 |
| Myotis stalkeri | Vespertilionidae | 0 | NA | 154 |
| Myotis thysanodes | Vespertilionidae | 1 | mines and buildings | 96 |
| Myotis tricolor | Vespertilionidae | 1 | abandoned mines | 154 |
| Myotis velifer | Vespertilionidae | 1 | old buildings and under bridges | 154 |
| Myotis vivesi | Vespertilionidae | 1 | NA | 330 |
| Myotis volans | Vespertilionidae | 1 | buildings, mines | 154 |
| Myotis welwitschii | Vespertilionidae | 1 | factories and houses | 331 |
| Myotis yanbarensis | Vespertilionidae | 0 | NA | 154 |
| Myotis yumanensis | Vespertilionidae | 1 | buildings, mines, underbridges | 96 |
| Mystacina robusta | Mystacinidae | 0 | NA | 331 |
| Mystacina tuberculata | Mystacinidae | 1 | bridges | 333 |
| Myzopoda aurita | Myzopodidae | 0 | NA | 334 |
| Myzopoda schliemanni | Myzopodidae | 0 | NA | 335, 336 |
| Nanonycteris veldkampii | Pteropodidae | 0 | NA | 83 |
| Natalus espiritosantensis | Natalidae | 0 | NA | 78 |
| Natalus jamaicensis | Natalidae | 0 | NA | 78 |
| Natalus lanatus | Natalidae | 1 | mine | 337 |
| Natalus major | Natalidae | 0 | NA | 78 |
| Natalus mexicanus | Natalidae | 1 | mine | 338 |
| Natalus primus | Natalidae | 0 | NA | 78 |
| Natalus stramineus | Natalidae | 1 | brick tunnel | 78 |
| Natalus tumidirostris | Natalidae | 1 | mines | 78 |
| Neoromicia brunnea | Vespertilionidae | 1 | roofs of unused houses | 339 |
| Neoromicia capensis | Vespertilionidae | 1 | between cracks in walls, under the roofs of houses | 86, 154 |
| Neoromicia matroka | Vespertilionidae | 1 | buildings | 154 |
| Neoromicia melckorum | Vespertilionidae | 1 | under roofs of buildings | 340 |
| Neoromicia nana | Vespertilionidae | 1 | roofs and in thatch of rural huts | 341 |
| Neoromicia rendalli | Vespertilionidae | 1 | thatched huts, brick walls and rafters | 154 |

| | | | | |
|---|---|---|---|---|
| Neoromicia somalica | Vespertilionidae | 1 | houses | 342 |
| Neoromicia tenuipinnis | Vespertilionidae | 1 | roofs, eaves of houses and small crevices in houses | 154 |
| Noctilio albiventris | Noctilionidae | 1 | man-made structures, buildings, installations associated with a Naval Base, bridge over marsh | 343 |
| Noctilio leporinus | Noctilionidae | 1 | bridges | 344 |
| Notopteris macdonaldi | Pteropodidae | 0 | NA | 83 |
| Notopteris neocaledonica | Pteropodidae | 0 | NA | 83 |
| Nyctalus aviator | Vespertilionidae | 1 | buildings and bird boxes, slits above railway | 345, 154 |
| Nyctalus azoreum | Vespertilionidae | 1 | buildings, houses | 154, 346 |
| Nyctalus furvus | Vespertilionidae | 1 | building | 154 |
| Nyctalus lasiopterus | Vespertilionidae | 1 | bat-boxes, buildings | 154 |
| Nyctalus leisleri | Vespertilionidae | 1 | bat-boxes, buildings | 347, 154 |
| Nyctalus montanus | Vespertilionidae | 1 | roof of bungalow | 154 |
| Nyctalus noctula | Vespertilionidae | 1 | buildings, bat-boxes, church steeple, attics | 348, 349, 154 |
| Nyctalus plancyi | Vespertilionidae | 1 | old temples, under roof tiles, various parts of buildings | 154 |
| Nycteris arge | Nycteridae | 1 | culverts | 154, 1 |
| Nycteris gambiensis | Nycteridae | 1 | roofs and cellars of buildings | 350, 1 |
| Nycteris grandis | Nycteridae | 1 | houses, disused water tower, culverts | 351, 1 |
| Nycteris hispida | Nycteridae | 1 | roofs and empty rooms of buildings | 352, 1 |
| Nycteris intermedia | Nycteridae | 1 | abandoned house | 352, 1 |
| Nycteris javanica | Nycteridae | 1 | culverts | 353, 1 |
| Nycteris macrotis | Nycteridae | 1 | thatched house roofs | 352, 1 |
| Nycteris major | Nycteridae | 1 | building | 352, 1 |
| Nycteris nana | Nycteridae | 1 | mines, culverts | 352, 1 |
| Nycteris thebaica | Nycteridae | 1 | mine tunnels, military bunkers, fireplaces, buildings, culverts, abandoned wells, pit latrines | 275, 1 |
| Nycteris tragata | Nycteridae | 1 | culverts | 354, 1 |
| Nycteris vinsoni | Nycteridae | 0 | NA | 355, 1 |
| Nycteris woodi | Nycteridae | 1 | buildings | 352, 1 |
| Nycticeinops schlieffeni | Vespertilionidae | 1 | buildings and cellars, roof of huts | 356, 1 |
| Nycticeius cubanus | Vespertilionidae | 1 | buildings and light posts | 154, 1 |
| Nycticeius humeralis | Vespertilionidae | 1 | Buildings | 96, 1 |
| Nyctiellus lepidus | Natalidae | 1 | abandoned hotel | 154, 1 |
| Nyctimene aello | Pteropodidae | 0 | NA | 357, 1 |
| Nyctimene albiventer | Pteropodidae | 0 | NA | 83, 1 |
| Nyctimene cephalotes | Pteropodidae | 0 | NA | 83, 1 |
| Nyctimene certans | Pteropodidae | 0 | NA | 83, 1 |
| Nyctimene cyclotis | Pteropodidae | 0 | NA | 1 |

| | | | | |
|---|---|---|---|---|
| Nyctimene draconilla | Pteropodidae | 0 | NA | 1 |
| Nyctimene keasti | Pteropodidae | 0 | NA | 1 |
| Nyctimene major | Pteropodidae | 0 | NA | 83, 1 |
| Nyctimene malaitensis | Pteropodidae | 0 | NA | 1 |
| Nyctimene masalai | Pteropodidae | 0 | NA | 1 |
| Nyctimene minutus | Pteropodidae | 0 | NA | 1 |
| Nyctimene rabori | Pteropodidae | 0 | NA | 1 |
| Nyctimene robinsoni | Pteropodidae | 0 | NA | 358, 1 |
| Nyctimene sanctacrucis | Pteropodidae | 0 | NA | 1 |
| Nyctimene vizcaccia | Pteropodidae | 0 | NA | 83, 1 |
| Nyctinomops aurispinosus | Molossidae | 1 | attic of a house | 359, 1 |
| Nyctinomops femorosaccus | Molossidae | 1 | Buildlings, under roof tiles | 360, 1 |
| Nyctinomops kalinowskii | Molossidae | 0 | NA | 361 |
| Nyctinomops laticaudatus | Molossidae | 1 | human-made structures and Mayan ruins | 362 |
| Nyctinomops macrotis | Molossidae | 1 | buildings, school building | 96, 363 |
| Nyctophilus arnhemensis | Vespertilionidae | 1 | under roof of house | 154 |
| Nyctophilus bifax | Vespertilionidae | 1 | houses | 364, 154 |
| Nyctophilus corbeni | Vespertilionidae | 0 | NA | 365 |
| Nyctophilus geoffroyi | Vespertilionidae | 1 | roofs of buildings | 366, 1 |
| Nyctophilus gouldi | Vespertilionidae | 1 | buildings | 367 |
| Nyctophilus microdon | Vespertilionidae | 0 | NA | 154 |
| Nyctophilus microtis | Vespertilionidae | 0 | NA | 154 |
| Nyctophilus sherrini | Vespertilionidae | 1 | timber hut | 368 |
| Nyctophilus timoriensis | Vespertilionidae | 0 | NA | 369, 365 |
| Otomops johnstonei | Molossidae | 0 | NA | 99 |
| Otomops madagascariensis | Molossidae | 0 | NA | 258 |
| Otomops martiensseni | Molossidae | 1 | attics of buildings | 99, 370 |
| Otomops papuensis | Molossidae | 0 | NA | 99 |
| Otomops secundus | Molossidae | 0 | NA | 371 |
| Otomops wroughtoni | Molossidae | 0 | NA | 99 |
| Otonycteris hemprichii | Vespertilionidae | 1 | human constructions | 372 |
| Paracoelops megalotis | Hipposideridae | 1 | disused mine | 205, 373 |
| Paranyctimene raptor | Pteropodidae | 0 | NA | 83, 1 |
| Paranyctimene tenax | Pteropodidae | 0 | NA | 83 |
| Parastrellus hesperus | Vespertilionidae | 1 | Mines, bridges | 154 |
| Paratriaenops auritus | Hipposideridae | 0 | NA | 374 |
| Paratriaenops furculus | Hipposideridae | 0 | NA | 375 |

| | | | | |
|---|---|---|---|---|
| Paratriaenops pauliani | Hipposideridae | 1 | houses | 376 |
| Penthetor lucasi | Pteropodidae | 0 | NA | 83, 1 |
| Peropteryx kappleri | Emballonuridae | 1 | abandoned coal mines, buildings | 377, 144 |
| Peropteryx leucoptera | Emballonuridae | 0 | NA | 210 |
| Peropteryx macrotis | Emballonuridae | 1 | bell towers and pre-columbian ruins | 378, 1 |
| Peropteryx pallidoptera | Emballonuridae | 0 | NA | 379 |
| Peropteryx trinitatis | Emballonuridae | 1 | houses in Venezuela | 380 |
| Pharotis imogene | Vespertilionidae | 0 | NA | 154 |
| Philetor brachypterus | Vespertilionidae | 0 | NA | 154 |
| Phoniscus atrox | Vespertilionidae | 0 | NA | 381 |
| Phoniscus jagorii | Vespertilionidae | 0 | NA | 382 |
| Phoniscus papuensis | Vespertilionidae | 0 | NA | 383 |
| Phyllonycteris aphylla | Phyllostomidae | 0 | NA | 41, 1 |
| Phyllonycteris poeyi | Phyllostomidae | 0 | NA | 384, 1 |
| Phyllops falcatus | Phyllostomidae | 1 | NA | 385, 1 |
| Phyllostomus discolor | Phyllostomidae | 0 | NA | 386 |
| Phyllostomus elongatus | Phyllostomidae | 1 | culverts, buildings | 232, 1 |
| Phyllostomus hastatus | Phyllostomidae | 1 | Thatched roofs | 387, 1 |
| Phyllostomus latifolius | Phyllostomidae | 0 | NA | 41 |
| Pipistrellus abramus | Vespertilionidae | 1 | including narrow spaces in buildings, under roof tiles, parapet caps, small roof covers over windows, inside sliding wings, under bridges | 154 |
| Pipistrellus adamsi | Vespertilionidae | 0 | NA | 154 |
| Pipistrellus alaschanicus | Vespertilionidae | 1 | eaves of houses, bridges | 154 |
| Pipistrellus angulatus | Vespertilionidae | 1 | buildings | 154 |
| Pipistrellus ariel | Vespertilionidae | 1 | ruins of a castle | 154, 388 |
| Pipistrellus cadornae | Vespertilionidae | 1 | old building and concrete bridge | 389 |
| Pipistrellus ceylonicus | Vespertilionidae | 1 | old dilapidated buildings, wells, and old temples | 154 |
| Pipistrellus collinus | Vespertilionidae | 0 | NA | 154 |
| Pipistrellus coromandra | Vespertilionidae | 1 | cracks in walls, ceilings of houses, tiles of huts | 154 |
| Pipistrellus deserti | Vespertilionidae | 1 | crevices in buildings | 390, 391 |
| Pipistrellus endoi | Vespertilionidae | 1 | buildings | 154, 1 |
| Pipistrellus hanaki | Vespertilionidae | 1 | buildings | 392 |
| Pipistrellus hesperidus | Vespertilionidae | 1 | iron roof | 86 |
| Pipistrellus imbricatus | Vespertilionidae | 0 | NA | 154 |
| Pipistrellus javanicus | Vespertilionidae | 1 | houses, window of office building | 154, 293, 1 |
| Pipistrellus kuhlii | Vespertilionidae | 1 | crevices in buildings | 391 |
| Pipistrellus macrotis | Vespertilionidae | 1 | crevice between pillar and pipeline in building | 154 |

| | | | | |
|---|---|---|---|---|
| Pipistrellus maderensis | Vespertilionidae | 1 | bird boxes, disused buildings | 394 |
| Pipistrellus nanulus | Vespertilionidae | 1 | house | 350, 395 |
| Pipistrellus nathusii | Vespertilionidae | 1 | buildings, bat boxes, houses, timber claddings,wooden churches | 154 |
| Pipistrellus papuanus | Vespertilionidae | 1 | buildings | 154 |
| Pipistrellus paterculus | Vespertilionidae | 1 | roofs of thatched huts | 154 |
| Pipistrellus pipistrellus | Vespertilionidae | 1 | cracks in building, houses, churches | 154 |
| Pipistrellus pulveratus | Vespertilionidae | 1 | houses | 154 |
| Pipistrellus pygmaeus | Vespertilionidae | 1 | roof of houses, attics, crevices in buildings, bat boxes | 396, 154 |
| Pipistrellus raceyi | Vespertilionidae | 1 | concrete wall of occupied building | 397 |
| Pipistrellus rueppellii | Vespertilionidae | 1 | buildings, board against wall, roost in tent flap | 398, 399, 154 |
| Pipistrellus rusticus | Vespertilionidae | 1 | old buildings | 154 |
| Pipistrellus savii | Vespertilionidae | 1 | fissures in buildings, attics, roof gutter, roller type shutters, abandoned shepherd cabins, tunnels | 400 |
| Pipistrellus stenopterus | Vespertilionidae | 1 | roofs of houses | 154 |
| Pipistrellus subflavus | Vespertilionidae | 1 | buildings, mines, abd bridges | 96, 401 |
| Pipistrellus tenuis | Vespertilionidae | 1 | walls and ceilings of buildings | 154 |
| Pipistrellus wattsi | Vespertilionidae | 1 | ceilings of houses | 154 |
| Pipistrellus westralis | Vespertilionidae | 0 | NA | 402 |
| Platalina genovensium | Phyllostomidae | 1 | mines, caves, and bridges | 41 |
| Platymops setiger | Molossidae | 0 | NA | 1 |
| Platyrrhinus albericoi | Phyllostomidae | 0 | NA | 403 |
| Platyrrhinus angustirostris | Phyllostomidae | 1 | buildings, bridges, and culverts | 404, 405 |
| Platyrrhinus aurarius | Phyllostomidae | 0 | NA | 41 |
| Platyrrhinus brachycephalus | Phyllostomidae | 0 | NA | 41 |
| Platyrrhinus chocoensis | Phyllostomidae | 0 | NA | 13 |
| Platyrrhinus dorsalis | Phyllostomidae | 0 | NA | 41 |
| Platyrrhinus fusciventris | Phyllostomidae | 1 | buildings, bridges, and culverts | 404, 405 |
| Platyrrhinus helleri | Phyllostomidae | 1 | buildings, bridges, and culverts | 404 |
| Platyrrhinus infuscus | Phyllostomidae | 0 | NA | 41 |
| Platyrrhinus ismaeli | Phyllostomidae | 0 | NA | 406 |
| Platyrrhinus lineatus | Phyllostomidae | 0 | NA | 41 |
| Platyrrhinus masu | Phyllostomidae | 0 | NA | 407 |
| Platyrrhinus nigellus | Phyllostomidae | 0 | NA | 408 |
| Platyrrhinus recifinus | Phyllostomidae | 0 | NA | 409 |
| Platyrrhinus umbratus | Phyllostomidae | 0 | NA | 410, 13 |
| Platyrrhinus vittatus | Phyllostomidae | 0 | NA | 403 |

| | | | | |
|---|---|---|---|---|
| Plecotus auritus | Vespertilionidae | 1 | attics of churches and barns, between roof tiles, behind wood cladding, bunkers, mines, and bird or bat boxes | 154, 1 |
| Plecotus austriacus | Vespertilionidae | 1 | roofs of cathedrals, cellars, churches, old buildings | 154, 1 |
| Plecotus christii | Vespertilionidae | 1 | remote buildings outside of urban areas | 411 |
| Plecotus kolombatovici | Vespertilionidae | 1 | bunkers | 154 |
| Plecotus macrobullaris | Vespertilionidae | 1 | attics of churches | 154 |
| Plecotus ognevi | Vespertilionidae | 1 | grotto, buildings, tunnel | 412 |
| Plecotus sacrimontis | Vespertilionidae | 1 | buildings in Japan | 154 |
| Plecotus sardus | Vespertilionidae | 1 | buildings | 413 |
| Plecotus taivanus | Vespertilionidae | 0 | mines, tunnels, buildings | 154 |
| Plecotus teneriffae | Vespertilionidae | 1 | crevices in abandoned buildings | 154 |
| Plerotes anchietae | Pteropodidae | 0 | NA | 1 |
| Promops centralis | Molossidae | 0 | roof tiles | 414 |
| Promops nasutus | Molossidae | 1 | roofs of houses | 1 |
| Ptenochirus jagori | Pteropodidae | 0 | NA | 1, 83 |
| Ptenochirus minor | Pteropodidae | 0 | NA | 1, 83 |
| Pteralopex anceps | Pteropodidae | 0 | NA | 83 |
| Pteralopex atrata | Pteropodidae | 0 | NA | 83 |
| Pteralopex flanneryi | Pteropodidae | 0 | NA | 83 |
| Pteralopex pulchra | Pteropodidae | 0 | NA | 1 |
| Pteralopex taki | Pteropodidae | 0 | NA | 83 |
| Pteronotus davyi | Mormoopidae | 1 | houses, mines, chicken coops, barns | 415 |
| Pteronotus gymnonotus | Mormoopidae | 0 | NA | 416 |
| Pteronotus macleayii | Mormoopidae | 0 | NA | 417 |
| Pteronotus paraguanensis | Mormoopidae | 0 | NA | 286 |
| Pteronotus parnellii | Mormoopidae | 1 | mines | 418, 419 |
| Pteronotus personatus | Mormoopidae | 1 | mines | 420 |
| Pteronotus quadridens | Mormoopidae | 0 | NA | 421 |
| Pteropus admiralitatum | Pteropodidae | 0 | NA | 83 |
| Pteropus aldabrensis | Pteropodidae | 0 | NA | 83 |
| Pteropus alecto | Pteropodidae | 0 | NA | 422 |
| Pteropus anetianus | Pteropodidae | 0 | NA | 423 |
| Pteropus argentatus | Pteropodidae | 0 | NA | 424 |
| Pteropus caniceps | Pteropodidae | 0 | NA | 83 |
| Pteropus capistratus | Pteropodidae | 0 | NA | 425 |
| Pteropus chrysoproctus | Pteropodidae | 0 | NA | 424 |
| Pteropus cognatus | Pteropodidae | 0 | NA | 426 |
| Pteropus conspicillatus | Pteropodidae | 0 | NA | 427 |

| | | | | |
|---|---|---|---|---|
| Pteropus dasymallus | Pteropodidae | 0 | NA | 83 |
| Pteropus faunulus | Pteropodidae | 0 | NA | 83 |
| Pteropus fundatus | Pteropodidae | 0 | NA | 83 |
| Pteropus giganteus | Pteropodidae | 0 | NA | 428 |
| Pteropus griseus | Pteropodidae | 0 | NA | 83 |
| Pteropus howensis | Pteropodidae | 0 | NA | 429 |
| Pteropus hypomelanus | Pteropodidae | 0 | NA | 430, 83 |
| Pteropus intermedius | Pteropodidae | 0 | NA | 44 |
| Pteropus livingstonii | Pteropodidae | 0 | NA | 432 |
| Pteropus lylei | Pteropodidae | 0 | NA | 433, 83 |
| Pteropus macrotis | Pteropodidae | 0 | NA | 434 |
| Pteropus mahaganus | Pteropodidae | 0 | NA | 435 |
| Pteropus mariannus | Pteropodidae | 0 | NA | 83, 436 |
| Pteropus melanopogon | Pteropodidae | 0 | NA | 424 |
| Pteropus melanotus | Pteropodidae | 0 | NA | 83 |
| Pteropus molossinus | Pteropodidae | 0 | NA | 83 |
| Pteropus neohibernicus | Pteropodidae | 0 | NA | 83 |
| Pteropus niger | Pteropodidae | 0 | NA | 437, 83 |
| Pteropus nitendiensis | Pteropodidae | 0 | NA | 83 |
| Pteropus ornatus | Pteropodidae | 0 | NA | 83 |
| Pteropus pelagicus | Pteropodidae | 0 | NA | 438 |
| Pteropus pelewensis | Pteropodidae | 0 | NA | 439 |
| Pteropus personatus | Pteropodidae | 0 | NA | 83 |
| Pteropus pohlei | Pteropodidae | 0 | NA | 340 |
| Pteropus poliocephalus | Pteropodidae | 0 | NA | 441, 442 |
| Pteropus pselaphon | Pteropodidae | 0 | NA | 343 |
| Pteropus pumilus | Pteropodidae | 0 | NA | 444, 445 |
| Pteropus rayneri | Pteropodidae | 0 | NA | 83 |
| Pteropus rennelli | Pteropodidae | 0 | NA | 83 |
| Pteropus rodricensis | Pteropodidae | 0 | NA | 446 |
| Pteropus rufus | Pteropodidae | 0 | NA | 447 |
| Pteropus samoensis | Pteropodidae | 0 | NA | 448 |
| Pteropus scapulatus | Pteropodidae | 0 | NA | 449 |
| Pteropus seychellensis | Pteropodidae | 0 | NA | 83 |
| Pteropus speciosus | Pteropodidae | 0 | NA | 83 |
| Pteropus temminckii | Pteropodidae | 0 | NA | 450 |
| Pteropus tonganus | Pteropodidae | 0 | NA | 451 |

| | | | | |
|---|---|---|---|---|
| Pteropus tuberculatus | Pteropodidae | 0 | NA | 83 |
| Pteropus ualanus | Pteropodidae | 0 | NA | 452 |
| Pteropus vampyrus | Pteropodidae | 0 | NA | 431 |
| Pteropus vetulus | Pteropodidae | 0 | NA | 83 |
| Pteropus voeltzkowi | Pteropodidae | 0 | NA | 453, 83 |
| Pteropus woodfordi | Pteropodidae | 0 | NA | 454 |
| Pteropus yapensis | Pteropodidae | 0 | NA | 439 |
| Pygoderma bilabiatum | Phyllostomidae | 1 | houses | 41 |
| Rhinolophus acuminatus | Rhinolophidae | 1 | field house, old buildings | 455, 456 |
| Rhinolophus adami | Rhinolophidae | 0 | NA | 456 |
| Rhinolophus affinis | Rhinolophidae | 0 | NA | 457, 456 |
| Rhinolophus alcyone | Rhinolophidae | 1 | roofs of thatched houses, old mines | 456 |
| Rhinolophus arcuatus | Rhinolophidae | 0 | NA | 458, 456 |
| Rhinolophus beddomei | Rhinolophidae | 1 | old temple, old forts, dilapidated buildings, unused bungalows, old tunnels | 456 |
| Rhinolophus blasii | Rhinolophidae | 1 | buildings, mine shafts, underground irrigation ditches | 456 |
| Rhinolophus bocharicus | Rhinolophidae | 1 | mine galleries | 456 |
| Rhinolophus borneensis | Rhinolophidae | 0 | NA | 456 |
| Rhinolophus canuti | Rhinolophidae | 0 | NA | 456 |
| Rhinolophus capensis | Rhinolophidae | 1 | lofts and disused mines | 456 |
| Rhinolophus celebensis | Rhinolophidae | 0 | NA | 456 |
| Rhinolophus chaseni | Rhinolophidae | 0 | NA | 459 |
| Rhinolophus clivosus | Rhinolophidae | 1 | disused mines, urban and rural buildings | 456 |
| Rhinolophus coelophyllus | Rhinolophidae | 0 | NA | 460 |
| Rhinolophus cognatus | Rhinolophidae | 0 | NA | 461, 456 |
| Rhinolophus cohenae | Rhinolophidae | 1 | abandoned mine shafts/tunnels | 456 |
| Rhinolophus creaghi | Rhinolophidae | 0 | NA | 456 |
| Rhinolophus darlingi | Rhinolophidae | 1 | mines, unused buildings | 132, 456 |
| Rhinolophus deckenii | Rhinolophidae | 0 | mud house | 456 |
| Rhinolophus denti | Rhinolophidae | 1 | abandoned mine, culverts, thatched roof | 456, 132 |
| Rhinolophus eloquens | Rhinolophidae | 1 | mines | 462 |
| Rhinolophus euryale | Rhinolophidae | 1 | attics, abandoned mines, railway tunnels, and military bunkers | 463, 456 |
| Rhinolophus euryotis | Rhinolophidae | 1 | mine adits and military tunnels | 464 |
| Rhinolophus ferrumequinum | Rhinolophidae | 1 | roofs of buildings | 456 |
| Rhinolophus formosae | Rhinolophidae | 1 | abandoned tunnels, buildings, underground irrigation channels | 456 |
| Rhinolophus fumigatus | Rhinolophidae | 1 | abandoned mine shafts | 456 |
| Rhinolophus guineensis | Rhinolophidae | 1 | mine adit | 465 |

| | | | | |
|---|---|---|---|---|
| Rhinolophus hildebrandtii | Rhinolophidae | 1 | mines, unused buildings, roofs of houses | 456, 133 |
| Rhinolophus hilli | Rhinolophidae | 0 | old mining tunnels | 466 |
| Rhinolophus hillorum | Rhinolophidae | 1 | bridges, culverts | 467, 456 |
| Rhinolophus hipposideros | Rhinolophidae | 1 | roofs, tunnels, attics, and cellars | 456 |
| Rhinolophus huananus | Rhinolophidae | 0 | NA | 468 |
| Rhinolophus inops | Rhinolophidae | 0 | NA | 469, 456 |
| Rhinolophus kahuzi | Rhinolophidae | 0 | NA | 470 |
| Rhinolophus keyensis | Rhinolophidae | 0 | NA | 471 |
| Rhinolophus landeri | Rhinolophidae | 1 | mine adit, thatched buildings, wells | 472, 456 |
| Rhinolophus lepidus | Rhinolophidae | 1 | unused tunnels, old and ruined buildings, old temples | 456 |
| Rhinolophus luctus | Rhinolophidae | 1 | old mine shafts | 473 |
| Rhinolophus maclaudi | Rhinolophidae | 1 | house | 474, 456 |
| Rhinolophus macrotis | Rhinolophidae | 1 | abandoned mine | 456 |
| Rhinolophus madurensis | Rhinolophidae | 0 | NA | 456 |
| Rhinolophus maendeleo | Rhinolophidae | 0 | NA | 456 |
| Rhinolophus malayanus | Rhinolophidae | 0 | NA | 475, 456 |
| Rhinolophus marshalli | Rhinolophidae | 0 | NA | 456 |
| Rhinolophus megaphyllus | Rhinolophidae | 1 | mine adits, unused buildings | 476, 456 |
| Rhinolophus mehelyi | Rhinolophidae | 1 | abandoned house | 477, 456 |
| Rhinolophus montanus | Rhinolophidae | 0 | NA | 456 |
| Rhinolophus mossambicus | Rhinolophidae | 0 | NA | 478 |
| Rhinolophus osgoodi | Rhinolophidae | 0 | NA | 479 |
| Rhinolophus paradoxolophus | Rhinolophidae | 0 | NA | 480 |
| Rhinolophus pearsonii | Rhinolophidae | 1 | bomb shelter | 481 |
| Rhinolophus philippinensis | Rhinolophidae | 1 | abandoned mines | 456 |
| Rhinolophus pusillus | Rhinolophidae | 1 | roofs of bungalows | 456 |
| Rhinolophus rex | Rhinolophidae | 1 | buildlings | 482 |
| Rhinolophus robinsoni | Rhinolophidae | 0 | NA | 456 |
| Rhinolophus rouxii | Rhinolophidae | 1 | disused wells, dilapidated buildings, and temples | 456 |
| Rhinolophus rufus | Rhinolophidae | 0 | NA | 456 |
| Rhinolophus ruwenzorii | Rhinolophidae | 1 | abandoned mines | 483, 456 |
| Rhinolophus sakejiensis | Rhinolophidae | 0 | NA | 484 |
| Rhinolophus schnitzleri | Rhinolophidae | 0 | NA | 485 |
| Rhinolophus sedulus | Rhinolophidae | 1 | drainage pipes and culverts | 486, 456 |
| Rhinolophus shameli | Rhinolophidae | 0 | NA | 456 |
| Rhinolophus siamensis | Rhinolophidae | 0 | NA | 456 |

| | | | | |
|---|---|---|---|---|
| Rhinolophus simulator | Rhinolophidae | 1 | mine adits | 132, 456 |
| Rhinolophus sinicus | Rhinolophidae | 1 | disused temples, houses, tunnels, wells | 487, 456 |
| Rhinolophus smithersi | Rhinolophidae | 1 | disuse mine adits | 488 |
| Rhinolophus stheno | Rhinolophidae | 0 | NA | 489, 456 |
| Rhinolophus subbadius | Rhinolophidae | 0 | NA | 487, 490 |
| Rhinolophus subrufus | Rhinolophidae | 0 | NA | 491, 456 |
| Rhinolophus swinnyi | Rhinolophidae | 1 | disused mines | 492, 132 |
| Rhinolophus thailandensis | Rhinolophidae | 0 | NA | 456 |
| Rhinolophus thomasi | Rhinolophidae | 0 | NA | 493, 494 |
| Rhinolophus trifoliatus | Rhinolophidae | 0 | NA | 354, 456 |
| Rhinolophus virgo | Rhinolophidae | 1 | culvert | 456 |
| Rhinolophus willardi | Rhinolophidae | 1 | mines | 495 |
| Rhinolophus xinanzhongguoensis | Rhinolophidae | 0 | NA | 496 |
| Rhinolophus yunanensis | Rhinolophidae | 1 | thatched roofs | 456 |
| Rhinolophus ziama | Rhinolophidae | 1 | mine-shafts, houses | 497 |
| Rhinonicteris aurantia | Hipposideridae | 1 | mines | 1, 87 |
| Rhinophylla pumilio | Phyllostomidae | 1 | culverts, thatched roofs | 498 |
| Rhinopoma hadramauticum | Rhinopomatidae | 1 | unused house | 499 |
| Rhinopoma hardwickii | Rhinopomatidae | 1 | ruins, catacombs, old buildings, mosques | 500, 1 |
| Rhinopoma macinnesi | Rhinopomatidae | 1 | pyramids, palaces, houses | 1 |
| Rhinopoma microphyllum | Rhinopomatidae | 1 | ruins, mosques, temples, tunnels, old buildings | 501, 1 |
| Rhinopoma muscatellum | Rhinopomatidae | 1 | disused buildings | 502, 1 |
| Rhogeessa minutilla | Vespertilionidae | 0 | NA | 503 |
| Rhogeessa parvula | Vespertilionidae | 0 | NA | 504 |
| Rhogeessa tumida | Vespertilionidae | 1 | church | 283, 154 |
| Rhynchonycteris naso | Emballonuridae | 1 | culverts, under bridges | 210 |
| Rousettus aegyptiacus | Pteropodidae | 1 | underground irrigation tunnels (ghanats), ruins, tombs, mines, military bunkers, underground parkings and open wells | 505, 83 |
| Rousettus amplexicaudatus | Pteropodidae | 1 | old tombs | 83 |
| Rousettus bidens | Pteropodidae | 0 | NA | 83 |
| Rousettus celebensis | Pteropodidae | 0 | NA | 506, 83 |
| Rousettus lanosus | Pteropodidae | 1 | mine adit | 507 |
| Rousettus leschenaultii | Pteropodidae | 1 | forts, houses, water tunnels, abandoned buildings | 508, 509 |
| Rousettus madagascariensis | Pteropodidae | 0 | NA | 83, 510 |
| Rousettus obliviosus | Pteropodidae | 0 | NA | 83 |
| Rousettus spinalatus | Pteropodidae | 0 | NA | 83 |
| Saccolaimus flaviventris | Emballonuridae | 1 | buildings | 144 |

| | | | | |
|---|---|---|---|---|
| Saccolaimus mixtus | Emballonuridae | 0 | NA | 144 |
| Saccolaimus peli | Emballonuridae | 0 | NA | 144 |
| Saccolaimus saccolaimus | Emballonuridae | 1 | old tombs and buildings | 1, 144 |
| Saccopteryx antioquensis | Emballonuridae | 1 | church wall | 144 |
| Saccopteryx bilineata | Emballonuridae | 1 | buildings, under bridges, ruins | 511, 1 |
| Saccopteryx canescens | Emballonuridae | 0 | NA | 512 |
| Saccopteryx leptura | Emballonuridae | 0 | NA | 513 |
| Sauromys petrophilus | Molossidae | 0 | NA | 1, 514 |
| Scoteanax rueppellii | Vespertilionidae | 1 | roofs of old buildings | 1 |
| Scotoecus albofuscus | Vespertilionidae | 0 | NA | 515, 154 |
| Scotoecus hirundo | Vespertilionidae | 1 | under roofs of huts and houses | 154 |
| Scotoecus pallidus | Vespertilionidae | 1 | old buildings | 154 |
| Scotomanes ornatus | Vespertilionidae | 0 | NA | 1 |
| Scotonycteris bergmansi | Pteropodidae | 0 | NA | 516 |
| Scotonycteris occidentalis | Pteropodidae | 0 | NA | 517 |
| Scotonycteris zenkeri | Pteropodidae | 0 | NA | 518, 1 |
| Scotophilus collinus | Vespertilionidae | 1 | roofs of houses | 154 |
| Scotophilus dinganii | Vespertilionidae | 1 | roof of building, metal weather-covering of an electrical conduit pipe | 519, 154 |
| Scotophilus heathii | Vespertilionidae | 1 | old forts, buildings | 154 |
| Scotophilus kuhlii | Vespertilionidae | 1 | roofs of old houses, temples, and huts | 520, 154 |
| Scotophilus leucogaster | Vespertilionidae | 1 | under iron roofs of old houses | 154 |
| Scotophilus marovaza | Vespertilionidae | 1 | roof of house made of palm leaves | 521 |
| Scotophilus nigrita | Vespertilionidae | 1 | house | 154 |
| Scotophilus robustus | Vespertilionidae | 1 | house, old school building, government office | 522 |
| Scotophilus viridis | Vespertilionidae | 0 | NA | 523 |
| Scotorepens balstoni | Vespertilionidae | 1 | building roofs, under metal caps of power poles | 154, 1 |
| Scotorepens greyii | Vespertilionidae | 1 | disused buildings | 154, 1 |
| Scotorepens orion | Vespertilionidae | 1 | buildings and nest boxes | 524, 1 |
| Scotorepens sanborni | Vespertilionidae | 1 | buildings | 154, 1 |
| Scotozous dormeri | Vespertilionidae | 1 | under roof tiles, old temples, disused buildings, and tombs | 1, 154 |
| Sphaeronycteris toxophyllum | Phyllostomidae | 0 | NA | 525 |
| Stenoderma rufum | Phyllostomidae | 0 | NA | 41 |
| Sturnira bidens | Phyllostomidae | 0 | NA | 526 |
| Sturnira erythromos | Phyllostomidae | 0 | NA | 527 |
| Sturnira hondurensis | Phyllostomidae | 0 | NA | 528 |

| | | | | |
|---|---|---|---|---|
| Sturnira lilium | Phyllostomidae | 1 | buildings | 529 |
| Sturnira ludovici | Phyllostomidae | 1 | abandoned mine shaft | 530 |
| Sturnira parvidens | Phyllostomidae | 0 | NA | 531 |
| Sturnira tildae | Phyllostomidae | 0 | NA | 532 |
| Submyotodon latirostris | Vespertilionidae | 1 | buildings | 154 |
| Syconycteris australis | Pteropodidae | 0 | NA | 533, 83 |
| Tadarida aegyptiaca | Molossidae | 1 | walls, creviced in old buildings, temples, banner boards, and forts | 534 |
| Tadarida australis | Molossidae | 1 | ceilings of buildings | 535 |
| Tadarida brasiliensis | Molossidae | 1 | bridges, attics | 99, 1 |
| Tadarida fulminans | Molossidae | 0 | NA | 99 |
| Tadarida insignis | Molossidae | 1 | outer walls of building and abandoned tunnel | 536 |
| Tadarida jobimena | Molossidae | 0 | NA | 99 |
| Tadarida kuboriensis | Molossidae | 0 | NA | 537 |
| Tadarida latouchei | Molossidae | 0 | NA | 99 |
| Tadarida lobata | Molossidae | 0 | NA | 538 |
| Tadarida teniotis | Molossidae | 1 | bridges, buildings | 99, 539 |
| Tadarida ventralis | Molossidae | 1 | high rise building, houses | 540, 99 |
| Taphozous achates | Emballonuridae | 0 | NA | 144, 541 |
| Taphozous australis | Emballonuridae | 1 | Old building, mines, and WWII bunkers | 144, 1 |
| Taphozous georgianus | Emballonuridae | 1 | vertical mine adits and horizontal shafts | 144 |
| Taphozous hamiltoni | Emballonuridae | 0 | NA | 144 |
| Taphozous hildegardeae | Emballonuridae | 0 | NA | 144, 542 |
| Taphozous hilli | Emballonuridae | 1 | disused mines | 144 |
| Taphozous kapalgensis | Emballonuridae | 0 | NA | 144 |
| Taphozous longimanus | Emballonuridae | 1 | old tunnels, caves created due to mud excavation, old forts, dungeons, wells, and eaves of houses | 144 |
| Taphozous mauritianus | Emballonuridae | 1 | outer walls of buildings beneath overhanging eaves, | 542, 543 |
| Taphozous melanopogon | Emballonuridae | 1 | old dilapidated buildings, dungeons of old forts, temples, abandoned mines, tunnels and churches | 544 |
| Taphozous nudiventris | Emballonuridae | 1 | tombs, temples, barns, houses, and underground tunnels | 144 |
| Taphozous perforatus | Emballonuridae | 1 | old forts, mosques, large old wells, artificial tunnels, and old disused buildings | 546 |
| Taphozous theobaldi | Emballonuridae | 0 | NA | 144 |
| Taphozous troughtoni | Emballonuridae | 1 | abandoned mines and tunnels | 547 |
| Thyroptera devivoi | Thyropteridae | 0 | NA | 548, 549 |
| Thyroptera discifera | Thyropteridae | 0 | NA | 549, 550, 1 |
| Thyroptera lavali | Thyropteridae | 0 | NA | 1, 551, 549 |
| Thyroptera tricolor | Thyropteridae | 0 | NA | 552, 549, 1 |

| | | | | |
|---|---|---|---|---|
| Thyroptera wynneae | Thyropteridae | 0 | NA | 551, 549 |
| Tomopeas ravus | Molossidae | 0 | NA | 1, 99 |
| Tonatia bidens | Phyllostomidae | 1 | water mines | 553, 41 |
| Tonatia saurophila | Phyllostomidae | 0 | NA | 554, 443 |
| Trachops cirrhosus | Phyllostomidae | 1 | mayan building, culverts, abandoned railroad tunnel | 1, 41 |
| Triaenops afer | Hipposideridae | 1 | abandoned mine | 132, 87 |
| Triaenops menamena | Hipposideridae | 1 | mine shaft | 555 |
| Triaenops parvus | Hipposideridae | 0 | NA | 87 |
| Triaenops persicus | Hipposideridae | 1 | mines and underground water tunnels | 1, 87 |
| Triaenops rufus | Hipposideridae | 1 | NA | 556 |
| Trinycteris nicefori | Phyllostomidae | 1 | concrete building, tunnel | 41 |
| Tylonycteris pachypus | Vespertilionidae | 0 | NA | 557, 1 |
| Tylonycteris pygmaeus | Vespertilionidae | 0 | NA | 558 |
| Tylonycteris robustula | Vespertilionidae | 0 | NA | 1, 154 |
| Uroderma bakeri | Phyllostomidae | 0 | NA | 1 |
| Uroderma bilobatum | Phyllostomidae | 0 | NA | 559, 1 |
| Uroderma magnirostrum | Phyllostomidae | 0 | NA | 41, 1 |
| Vampyressa bidens | Phyllostomidae | 0 | NA | 560, 561 |
| Vampyressa nymphaea | Phyllostomidae | 0 | NA | 562, 41 |
| Vampyressa pusilla | Phyllostomidae | 0 | NA | 563 |
| Vampyressa thyone | Phyllostomidae | 0 | NA | 41 |
| Vampyrodes caraccioli | Phyllostomidae | 0 | NA | 41 |
| Vampyrum spectrum | Phyllostomidae | 1 | church | 1 |
| Vespadelus baverstocki | Vespertilionidae | 1 | abandoned buildings | 154 |
| Vespadelus caurinus | Vespertilionidae | 1 | disused mines and road culverts | 154 |
| Vespadelus darlingtoni | Vespertilionidae | 1 | bat boxes | 564 |
| Vespadelus douglasorum | Vespertilionidae | 1 | abandoned buildings | 154 |
| Vespadelus finlaysoni | Vespertilionidae | 1 | abandoned mines | 154 |
| Vespadelus pumilus | Vespertilionidae | 0 | NA | 154 |
| Vespadelus regulus | Vespertilionidae | 1 | houses | 154 |
| Vespadelus troughtoni | Vespertilionidae | 1 | shed with tin roof | 565 |
| Vespadelus vulturnus | Vespertilionidae | 0 | NA | 154 |
| Vespertilio murinus | Vespertilionidae | 1 | houses, crevices in tall buildings, nest boxes | 566 |
| Vespertilio sinensis | Vespertilionidae | 1 | houses, buildings, under bridges, and tunnels | 567 |
| Xeronycteris vieirai | Phyllostomidae | 0 | NA | 568 |

**Table S4. Feature coverage across 1,279 bat species included in the BRT models.** Variable names as provided from their original sources (Jones et al. 2009, Soria et al. 2021, IUCN 2022)

| Variable | Coverage |
|---|---|
| cites | 1 |
| fam_EMBALLONURIDAE | 1 |
| fam_HIPPOSIDERIDAE | 1 |
| fam_MINIOPTERIDAE | 1 |
| fam_MOLOSSIDAE | 1 |
| fam_MORMOOPIDAE | 1 |
| fam_NATALIDAE | 1 |
| fam_NYCTERIDAE | 1 |
| fam_PHYLLOSTOMIDAE | 1 |
| fam_PTEROPODIDAE | 1 |
| fam_RHINOLOPHIDAE | 1 |
| fam_VESPERTILIONIDAE | 1 |
| category | 0.97 |
| population_trend | 0.97 |
| adult_mass_g | 0.94 |
| dphy_invertebrate | 0.94 |
| dphy_vertebrate | 0.94 |
| dphy_plant | 0.94 |
| trophic_level | 0.94 |
| foraging_stratum | 0.92 |
| activity_cycle | 0.92 |
| Indomalayan | 0.92 |
| Neotropical | 0.92 |
| Oceanian | 0.92 |

| | |
|---|---|
| Afrotropical | 0.92 |
| Australasian | 0.92 |
| Palearctic | 0.92 |
| Nearctic | 0.92 |
| habitat_breadth_n | 0.9 |
| X26.1_GR_Area_km2 | 0.83 |
| X26.2_GR_MaxLat_dd | 0.83 |
| X26.3_GR_MinLat_dd | 0.83 |
| X26.4_GR_MidRangeLat_dd | 0.83 |
| X26.5_GR_MaxLong_dd | 0.83 |
| X26.6_GR_MinLong_dd | 0.83 |
| X26.7_GR_MidRangeLong_dd | 0.83 |
| X27.1_HuPopDen_Min_n.km2 | 0.83 |
| X27.2_HuPopDen_Mean_n.km2 | 0.83 |
| X27.3_HuPopDen_5p_n.km2 | 0.83 |
| X27.4_HuPopDen_Change | 0.83 |
| Synurbic | 0.81 |
| X28.1_Precip_Mean_mm | 0.79 |
| X28.2_Temp_Mean_01degC | 0.79 |
| adult_forearm_length_mm | 0.76 |
| det_inv | 0.75 |
| det_vend | 0.75 |
| det_vect | 0.75 |
| det_vfish | 0.75 |
| det_fruit | 0.75 |
| det_nect | 0.75 |
| det_seed | 0.75 |
| det_diet_breadth_n | 0.75 |

| | |
|---|---|
| X30.1_AET_Mean_mm | 0.72 |
| X30.2_PET_Mean_mm | 0.72 |
| island_dwelling | 0.6 |
| disected_by_mountains | 0.6 |
| glaciation | 0.6 |
| litter_size_n | 0.55 |
| upper_elevation_m | 0.53 |
| lower_elevation_m | 0.53 |
| altitude_breadth_m | 0.45 |
| adult_body_length_mm | 0.42 |
| litters_per_year_n | 0.32 |

**Roosting Assignment References**


1. Nowak, R. M., & Walker, E. P. (1994). *Walker's Bats of the World*. JHU Press.
2. Freier, S. (2013). *Aethalops aequalis*. Animal Diversity Web. https://animaldiversity.org/accounts/Aethalops_aequalis/
3. Tamsitt, J. R., & Nagorsen, D. (1982). Anoura cultrata. *Mammalian Species*, *179*, 1–5.
4. Muchhala, N., Patricio, M. V., & Luis, A. V. (2005). A New Species of Anoura (Chiroptera: Phyllostomidae) from the Ecuadorian Andes. *Journal of Mammalogy*, *86*(3), 457–461.
5. Mantilla, H, Molinari, & J. (2015). Anoura latidens. *IUCN Red List of Threatened Species*, e.T40776A22134204.
6. Molinari, J. (1994). A new species of Anoura (Mammalia Chiroptera Phyllostomidae) from the Andes of northern South America. *Tropical Zoology*, *7*(1), 73–86.
7. Vaughan, T. A. (1976). Roosting Ecology of the Pallid Bat, Antrozous pallidus. *Journal of Mammalogy*, *57*(1), 19–42.
8. Lewis, S. E. (1994). Night Roosting Ecology of Pallid Bats (Antrozous pallidus) in Oregon. *The American Midland Naturalist*, *132*(2), 219–226.
9. Sagot, M., & Chaverri, G. (2015). Effects of roost specialization on extinction risk in bats. *Conservation Biology: The Journal of the Society for Conservation Biology*, *29*(6), 1666–1673.
10. Flannery, T. F., & Seri, L. (1993). Rediscovery of Aproteles bulmerae (Chiroptera: Pteropodidae). Morphology, ecology and conservation. *Mammalia*, *57*(1), 19–26.
11. Francis, C. M., & Csorba, G. (2020). Arielulus societatis. *IUCN Red List of Threatened Species*, e.T40776A22134204.

12. Ruiz-Ramoni, D., Muñoz-Romo, M., Ramoni-Perazzi, P., Aranguren, Y., & Fermin, G. (2011). Folivory in the Giant Fruit-Eating Bat Artibeus amplus (Phyllostomidae): A Non-Seasonal Phenomenon. *Acta Chiropterologica*, *13*(1), 195–199.
13. Garbino, G. S. T., & Tavares, V. da C. (2018). Roosting ecology of Stenodermatinae bats (Phyllostomidae): evolution of foliage roosting and correlated phenotypes. *Mammal Review*, *48*(2), 75–89.
14. Salas, J. A., Loaiza S, C. R., & Pacheco, V. (2018). Artibeus fraterculus (Chiroptera: Phyllostomidae). *Mammalian Species*, *50*(962), 67–73.
15. Webster, W. D., & Jones, J. K. (1983). Artibeus hirsutus and Artibeus inopinatus. *Mammalian Species*, *199*, 1–3.
16. Perini, F. A., Tavaores, V. C., & Nascimento, C. (2014). Bats from the city of Belo Horizonte, Minas Gerais, southeastern Brazil. *Chiroptera Neotropical*, *9*(1-2), 169–173.
17. Sampaio, E., Lim, B., & Peters, S. (2016). Artibeus obscurus. *IUCN Red List of Threatened Species*, e.T2137A21998064.
18. Haynes, M. A., & Lee, T. E. (2004). Artibeus obscurus. *Mammalian Species*, *752*, 1–5.
19. Hollis, L. (2005). Artibeus planirostris. *Mammalian Species*, *775*, 1–6.
20. Benda, P. (2016). Asellia arabica. *IUCN Red List of Threatened Species*, e.T80222726A95642180.
21. Hill, J. E., & Morris, P. (1971). Bats from Ethiopia collected by the Great Abbai Expedition, 1968. *Bulletin of the British Museum. Zoology*, *21*, 27–49.
22. Monadjem, A., Bergmans, W., Mickleburgh, S., & Hutson, A. M. (2016). Asellia patrizii. *IUCN Red List of Threatened Species*, e.T2153A21975955.

23. Monadjem, A., Bergmans, W., Mickleburgh, S., Kock, D., Amr, Z. S. S., & Hutson, A. M. (2016). Asellia tridens. *IUCN Red List of Threatened Species*, e.T80221529A21975715.
24. Armstrong, K. N. (2020). Aselliscus tricuspidatus. *IUCN Red List of Threatened Species*, e.T2156A21976047.
25. Tirira, D. (2014). Balantiopteryx infusca. *IUCN Red List of Threatened Species*, e.T2531A97206692.
26. Burton, L. (2014). Balantiopteryx io. *IUCN Red List of Threatened Species*, e.T2532A22030080.
27. Lim, B., Miller, B., Reid, F., Arroyo-Cabrales, J., Cuarón, A. D., & de Grammont, P. C. (2016). Balantiopteryx plicata. *IUCN Red List of Threatened Species*, e.T2533A22029659.
28. Zhang, J.-S., Han, N.-J., Jones, G., Lin, L.-K., Zhang, J.-P., Zhu, G.-J., Huang, D.-W., & Zhang, S.-Y. (2007). A New Species of Barbastella (Chiroptera: Vespertilionidae) from North China. *Journal of Mammalogy*, *88*(6), 1393–1403.
29. Solari, S. (2018). Bauerus dubiaquercus. *IUCN Red List of Threatened Species*, e.T1789A22129523.
30. Swanepoel, P., & Genoways, H. H. (1983). Brachyphylla cavernarum. *Mammalian Species*, *205*, 1–6.
31. Arnold, B., De La Cruz Mora, J. M., & Roesch, J. (2022). Assessing the Structure and Function of Distress Calls in Cuban Fruit-Eating Bats (Brachyphylla nana). *Frontiers in Ecology and Evolution*, *10*. https://doi.org/10.3389/fevo.2022.907751
32. Swanepoel, P., & Genoways, H. H. (1983). Brachyphylla nana. *Mammalian Species*, *206*, 1–3.
33. Csada, R. (1996). Cardioderma cor. *Mammalian Species*, *519*, 1–4.
34. Pearch, M. J., Bates, P. J. J., & Magin, C. (2001). A review of the small mammal fauna of Djibouti and the results of a recent survey. *Mammalia*, *65*(3), 387–410.

35. Gutierrez-Sanabria, D. R., & Lizcano, D. J. (2022). Dinámica poblacional y fidelidad de refugio de Carollia brevicauda (Chiroptera: Phyllostomidae) en un refugio artificial, en los Andes nororientales de Colombia. *Mammalogy Notes*, *8*(1), 203–203.
36. Kelm, D. H., Toelch, U., & Jones, M. M. (2021). Mixed-species groups in bats: non-random roost associations and roost selection in neotropical understory bats. *Frontiers in Zoology*, *18*(1), 53.
37. Thies, W., Kalko, E. K. V., & Schnitzler, H.-U. (2006). Influence of Environment and Resource Availability on Activity Patterns of Carollia castanea (Phyllostomidae) in Panama. *Journal of Mammalogy*, *87*(2), 331–338.
38. Velazco, P. M., & Aguirre, L. (2015). Carollia manu. *IUCN Red List of Threatened Species*, e.T136782A22033116.
39. Cloutier, D., & Thomas, D. W. (1992). Carollia perspicillata. *Mammalian Species*, *417*, 1–9.
40. Miller, B., Reid, F., Arroyo-Cabrales, J., Cuarón, A. D., & de Grammont, P. C. (2015). Carollia sowelli. *IUCN Red List of Threatened Species*, e.T136268A22003903.
41. Wilson, D. E., & Mittermeier, R. A. (Eds.). (2019). Phyllostomidae. In *Handbook of the Mammals of the World – Volume 9 Bats* (pp. 444–583). Lynx Edicions, Barcelona.
42. Reid, F. (1997). *A Field Guide to the Mammals of Central America and Southeast Mexico*. Oxford University Press.
43. Fahr, J. (2013). Scotonycteris ophiodon Pohle's fruit bat. In *Mammals of Africa, Vol. IV: Hedgehogs, shrews and bats* (pp. 295–297). Bloomsbury.
44. Soisook, P., & Tsang, S. M. (2020). Pteropus intermedius. *IUCN Red List of Threatened Species*, e.T136841A22042098.

45. Arroyo-Cabrales, J., Miller, B., Reid, F., Cuarón, A. D., & de Grammont, P. C. (2015). Centronycteris centralis. *IUCN Red List of Threatened Species*, e.T136350A22023809.

46. Sampaio, E., Lim, B., & Peters, S. (2016). Centronycteris maximiliani. *IUCN Red List of Threatened Species*, e.T4112A22002444.

47. Monadjem, A., Fahr, J., Hutson, A. M., Mickleburgh, S., & Bergmans, W. (2016). Chaerephon aloysiisabaudiae. *IUCN Red List of Threatened Species*, e.T4305A22020676.

48. Bouchard, S. (2001). Chaerephon ansorgei. *Mammalian Species*, *660*, 1–3.

49. López-Baucells, A., Rocha, R., Andriatafika, Z., Tojosoa, T., Kemp, J., Forbes, K., & Cabeza, M. (2017). Roost selection by synanthropic bats in rural Madagascar: what makes non-traditional structures so tempting? *Hystrix* , *28*(1), 28–35.

50. Goodman, S. (2016). Chaerephon atsinanana. *IUCN Red List of Threatened Species*, e.T67360705A67360707.

51. Monadjem, A., Fahr, J., Bergmans, W., Mickleburgh, S., Hutson, A. M., & Cotterill, F. (2016). Chaerephon bemmeleni. *IUCN Red List of Threatened Species*, e.T4307A22020379.

52. Smithers, R. H. N., & Tello, J. L. P. L. (1976). *Check list and atlas of the mammals of Mozambique*. Salisbury (Zimbabwe) Trustees of the National Museums of Rhodesia.

53. Palmeirim, J. M., Champion, A., Naikatini, A., Niukula, J., Tuiwawa, M., Fisher, M., Yabaki-Gounder, M., Thorsteinsdóttir, S., Qalovaki, S., & Dunn, T. (2007). Distribution, status and conservation of the bats of the Fiji Islands. *Oryx: The Journal of the Fauna Preservation Society*, *41*(4), 509–519.

54. Scanlon, A., Petit, S., & Bottroff, G. (2014). The conservation status of bats in Fiji. *Oryx: The Journal of the Fauna Preservation Society*, *48*(3), 451–459.

55. Fenton, M. B., & Eger, J. L. (2002). Chaerephon chapini. *Mammalian Species*, *692*, 1–2.

56. Mickleburgh, S., Hutson, A. M., Bergmans, W., & Cotterill, F. P. D. (2018). Chaerephon gallagheri. *IUCN Red List of Threatened Species*, e.T4311A22019365.
57. Lumsden, L. F., Johnstone, P. D., & Temby, I. D. (1993). A roosting colony of the Northern Mastiff-bat Chaerephon jobensis at Derby, Western Australia. *Australasian Bat Society Newsletter*, 13–15.
58. Kutt, A., Milne, D., & Richards, G. (2008). Northern Freetail-bat Chaerephon jobensis. In S. Van Dyck & R. Strahan (Eds.), *The Mammals of Australia* (pp. 485–486). Reed New Holland.
59. McLellan, L. J. (1986). *Notes on bats of Sudan. American Museum novitates; no. 2839*.
60. Willis, C. K. R., Psyllakis, J. M., & Sleep, D. J. H. (2002). Chaerephon nigeriae. *Mammalian Species*, *710*, 1–3.
61. Csorba, G., Bumrungsri, S., Francis, C., Bates, P., Ong, P., Gumal, M., Kingston, T., Heaney, L., Balete, D. S., Molur, S., & Srinivasulu, C. (2018). Chaerephon plicatus. *IUCN Red List of Threatened Species*, e.T4316A22018444.
62. Bouchard, S. (1998). Chaerephon pumilus. *Mammalian Species*, *574*, 1–6.
63. Mickleburgh, S., Hutson, A. M., Bergmans, W., Fahr, J., & Cotterill, F. P. D. (2019). Chaerephon russatus. *IUCN Red List of Threatened Species*, e.T4319A22017886.
64. Pennay, M., & Leary, T. (2020). Chaerephon solomonis. *IUCN Red List of Threatened Species*, e.T4320A22017829.
65. Dwyer, P. D. (1966). Observations on Chalinolobus Dwyeri (Chiroptera: Vespertilionidae) in Australia. *Journal of Mammalogy*, *47*(4), 716–718.
66. Chruszcz, B., & Barclay, R. M. R. (2002). Chalinolobus gouldii. *Mammalian Species*, *690*, 1–4.
67. Griffiths, S. R., Bender, R., Godinho, L. N., Lentini, P. E., Lumsden, L. F., & Robert, K. A. (2017). Bat boxes are not a silver bullet conservation tool. *Mammal Review*, *47*(4), 261–265.

68. Young, R. A. (1979). Observations on parturition, litter size, and foetal development at birth in the Chocolate Wattled Bat, Chalinolobus morio (Vespertilionidae). *The Victorian Naturalist*, *96*, 90–91.

69. Sanderson, K. J., Napier, G., & Johnston, G. R. (2010). Observations of a large colony of bats roosting in a church. *Australian Mammalogy*, *32*(2), 161–163.

70. Thomson, B. G. (2020). Social interactions, roost usage and notes on the breeding system of the chocolate wattled bat (Chalinolobus morio) in south-east Queensland, Australia. *Australian Journal of Zoology*, *67*(6), 290–300.

71. Brescia, F. (2020). Chalinolobus neocaledonicus. *IUCN Red List of Threatened Species*, e.T4420A21984825.

72. Hutson, A. M., Schlitter, D., Csorba, G., Thomson, B., & McKenzie, N. (2020). Chalinolobus nigrogriseus. *IUCN Red List of Threatened Species*, e.T4421A21984276.

73. Pennay, M. (2020). Chalinolobus picatus. *IUCN Red List of Threatened Species*, e.T4422A21984147.

74. Sedgeley, J. A. (2003). Roost site selection and roosting behaviour in lesser short-tailed bats (Mystacina tuberculata) in comparison with long-tailed bats (Chalinolobus tuberculatus) in Nothofagus forest, Fiordland. *New Zealand Journal of Zoology*.

75. O'Donnell, C., & Sedgeley, J. A. (2006). Causes and consequences of tree-cavity roosting in a temperate bat, Chalinolobus tuberculatus, from New Zealand. In A. Zubaid, G. F. McCracken, & T. H. Kunz (Eds.), *Functional and Evolutionary Biology of Bats* (pp. 308–328). Oxford University Press.

76. Alviola, P. A., Duya, M. R., Alvarez, J., Fidelino, J., Gatan-Balbas, M., Pedregosa, M., Veluz, M. J., & Jakosalem P G Tanalgo. (2019). Cheiromeles parvidens. *IUCN Red List of Threatened Species*, e.T4600A22034921.

77. Leong, T. M., Teo, S. C., & Lim, K. K. P. (2009). The naked bulldog bat, Cheiromeles torquatus in Singapore-past and present records, with highlights on its unique morphology (Microchiroptera: Molossidae). *Nature in Singapore*.

78. Tejedor, A. (2011). Systematics of Funnel-Eared Bats (Chiroptera: Natalidae). *Bulletin of the American Museum of Natural History*, *2011*(353), 1–140.

79. Handley. (1976). Mammals of the Smithsonian Venezuelan Project. *Brigham Young University Science Bulletin*, *20*, 1–89.

80. Goodwin, G. G., & Greenhall, A. M. (1961). A review of the bats of Trinidad and Tobago: descriptions, rabies infection, and ecology. *Bulletin of the AMNH*, *122*(3).

81. Garbino, G. S. T., Lim, B. K., & Tavares, V. D. A. C. (2020). Systematics of big-eyed bats, genus Chiroderma Peters, 1860 (Chiroptera: Phyllostomidae). *Zootaxa*, *4846*(1), zootaxa.4846.1.1.

82. Goldman, E. A. (1920). *Mammals of Panama: (with Thirty-nine Plates)*. Smithsonian institution.

88. Huang, J. C.-C., Thong, V. D., & Ho, Y. (2019). Coelops frithii. *IUCN Red List of Threatened Species*, e.T5074A22030377.

89. Heaney, L. (2008). Coelops robinsoni. *IUCN Red List of Threatened Species*, e.T5076A11112095.

90. Dunlop, J. (1997). Coleura afra. *Mammalian Species*, *566*, 1–4.

91. Goodman, S. (2016). Coleura kibomalandy. *IUCN Red List of Threatened Species*, e.T80221085A95642170.

92. Gerlach, J., & Taylor, M. (2006). Habitat use, roost characteristics and diet of the Seychelles sheath-tailed bat Coleura seychellensis. *Acta Chiropterologica / Museum and Institute of Zoology, Polish Academy of Sciences*, *8*(1), 129–139.

93. Bernard, E. (2003). Cormura brevirostris. *Mammalian Species*, *737*, 1–3.

94. Solari, S. (2019). Corynorhinus mexicanus. *IUCN Red List of Threatened Species*, e.T17599A21976792.

95. Tumlison, R. (1992). Plecotus mexicanus. *Mammalian Species*, *401*, 1–3.

96. Harvey, M. J., Scott Altenbach, J., & Best, T. L. (2011). *Bats of the United States and Canada*. JHU Press.

97. Kunz, T. H., & Martin, R. A. (1982). Plecotus townsendii. *Mammalian Species*, *175*, 1–6.

98. Hill, J. E., & Smith, S. E. (1981). Craseonycteris thonglongyai. *Mammalian Species*, *160*, 1–4.

99. Wilson, D. E., & Mittermeier, R. A. (Eds.). (2019). Molossidae. In *Handbook of the Mammals of the World – Volume 9 Bats* (pp. 598–672). Barcelona: Lynx Edicions.

100. Solari, S. (2015). Cynomops greenhalli. *IUCN Red List of Threatened Species*, e.T13639A22109178.

101. Rodriguez, B, Miller, & B. (2015). Cynomops mexicanus. *IUCN Red List of Threatened Species*, e.T136611A21987867.

102. López Berrizbeitia, M. F., & Díaz, M. M. (2021). Cynomops planirostris (Chiroptera: Molossidae). *Mammalian Species*, *53*(1013), 174–185.

103. Csorba, G., Bumrungsri, S., Bates P., Gumal, M., Kingston, T., Molur, S., & Srinivasulu, C. (2019). Cynopterus brachyotis. *IUCN Red List of Threatened Species*, e.T6103A22113381.

104. Campbell, P., & Kunz, T. H. (2006). Cynopterus horsfieldii. *Mammalian Species*, *802*, 1–5.

105. Storz, J. F., & Kunz, T. H. (1999). Cynopterus sphinx. *Mammalian Species*, *613*, 1–8.

106. Tsang, S. (2016). Cynopterus titthaecheilus. *IUCN Red List of Threatened Species*, e.T6107A22114054.

113. Copete Mosquera, Y. del C., Rentería Machado, Y., Palacios Mosquera, L., Mantilla Meluk, H., & Jiménez Ortega, A. M. (2018). Plantas utilizadas como tiendas por murciélagos tenderos en la selva pluvial central del Chocó, Colombia. *Academia Colombiana de Ciencias Exactas, Físicas Y Naturales*, *42*(162).

114. Webster, W. D., & Jones, J. K. (1982). Artibeus toltecus. *Mammalian Species*, *178*, 1–3.

115. Solari, S. (2016). Dermanura watsoni. *IUCN Red List of Threatened Species*, e.T99586593A21997358.

116. Waldien, D. L., & Duya, M. R. (2020). Desmalopex leucopterus. *IUCN Red List of Threatened Species*, e.T18731A22081331.

117. Greenhall, A. M., Joermann, G., & Schmidt, U. (1983). Desmodus rotundus. *Mammalian Species*, *202*, 1–6.

118. Greenhall, A. M., & Schutt, W. A. (1996). Diaemus youngi. *Mammalian Species*, *533*, 1–7.

119. Ceballos, G., & Medellín, R. A. (1988). Diclidurus albus. *Mammalian Species*, *316*, 1–4.

120. Mantilla-Meluk, H., Jiménez-Ortega, A. M., Palacios, L., & Baker, R. J. (2009). Unexpected finding of diclidurus ingens, Hernandez-Camacho, 1955 (Chiroptera, emballonuridae), in the Colombian biogeographic Chocó. *Mastozoologia Neotropical*, *16*(1), 229–232.

121. Sampaio, E., Lim, B., & Peters, S. (2016). Diclidurus scutatus. *IUCN Red List of Threatened Species*, e.T6564A21986499.

122. Greenhall, A. M., Schmidt, U., & Joermann, G. (1984). Diphylla ecaudata. *Mammalian Species*, *227*, 1–3.

123. Leary, T., Helgen, K., & Bonaccorso, F. J. (2020). Dobsonia anderseni. *IUCN Red List of Threatened Species*, e.T136374A22012133.

124. Tsang, S. (2016). Dobsonia moluccensis. *IUCN Red List of Threatened Species*, e.T84882605A22033630.

125. Leary, T., Helgen, K., & Bonaccorso, F. J. (2020). Dobsonia pannietensis. *IUCN Red List of Threatened Species*, e.T6776A22034157.

126. Hutson, A. M., Suyanto, A., & Helgen, K. (2019). Dobsonia peronii. *IUCN Red List of Threatened Species*, e.T6771A22034782.

127. Leary, T., Helgen, K., & Bonaccorso, F. J. (2019). Dobsonia praedatrix. *IUCN Red List of Threatened Species*, e.T6777A22033332.

128. Tsang, S. (2016). Dobsonia viridis. *IUCN Red List of Threatened Species*, e.T6780A22033412.

129. Helgen, K. M., Kock, D., Gomez, R. K. S. C., Ingle, N. R., & Sinaga, M. H. (2007). Taxonomy and Natural History of the Southeast Asian Fruit-Bat Genus Dyacopterus. *Journal of Mammalogy*, *88*(2), 302–318.

130. Timm, R. M. (1982). Ectophylla alba. *Mammalian Species*, *166*, 1–4.

131. Andriafidison, D., Andrianaivoarivelo, R., Cardiff, S. G., Goodman, S. M., Hutson, A. M., Jenkins, R. K. B., Kofoky, A., Picot, M., Racey, P. A., Ranivo, J., Ratrimomanarivo, F. H., & Razafimanahaka, J. (2020). Eidolon dupreanum. *IUCN Red List of Threatened Species*, e.T7083A22027891.

132. Skinner, J. D., & Chimimba, C. T. (2005). *The Mammals of the Southern African Sub-region*. Cambridge University Press.

133. DeFrees, S. L., & Wilson, D. E. (1988). Eidolon helvum. *Mammalian Species*, *312*, 1–5.

134. Armstrong, K. N., & Wiantoro, S. (2021). Emballonura alecto. *IUCN Red List of Threatened Species*, e.T7670A209548087.

135. Heaney, L., Balete, D., Dolar, L., & Ong, P. (1998). A Synopsis of the Mammalian Fauna of the Philippine Islands. *Fieldiana. Zoology*.

136. Monadjem, A., Cardiff, S. G., & Rakotoarivelo, A.R., Jenkins, R.K.B., Ratrimomanarivo, F.H. (2017). Paremballonura atrata. *IUCN Red List of Threatened Species*, e.T7671A22135427.

137. Armstrong, K. (2021). Emballonura beccarii (amended version of 2019 assessment). *IUCN Red List of Threatened Species*, e.T7672A209521847.

138. Armstrong, K. (2021). Emballonura dianae (amended version of 2019 assessment). *IUCN Red List of Threatened Species*, e.T7673A209522232.

139. Armstrong, K. N., & Aplin, K. (2021). Emballonura furax (amended version of 2017 assessment). *IUCN Red List of Threatened Species*, e.T7667A209536771.

140. Bates, P. J. J., Francis, C. M., & Kingston, T. (2021). Emballonura monticola. *IUCN Red List of Threatened Species*, e.T7674A22134864.

141. Armstrong, K. (2021). Emballonura raffrayana (amended version of 2019 assessment). *IUCN Red List of Threatened Species*, e.T7668A209522673.

142. Waldien, D. L., & Scanlon, A. (2021). Emballonura semicaudata. *IUCN Red List of Threatened Species*, e.T7669A22135085.

143. Armstrong, K. (2021). Emballonura serii (amended version of 2019 assessment). *IUCN Red List of Threatened Species*, e.T41528A209523175.

149. Barquez, R., Perez, S., Miller, B., & Diaz, M. (2016). Eptesicus brasiliensis. *IUCN Red List of Threatened Species*, e.T7916A22114459.

150. Mies, R., Kurta, A., & King, D. G. (1996). Eptesicus furinalis. *Mammalian Species*, *526*, 1–7.

151. Kurta, A., & Baker, R. H. (1990). Eptesicus fuscus. *Mammalian Species*, *356*, 1–10.

152. Srinivasulu, C., & Srinivasulu, B. (2018). Eptesicus gobiensis. *IUCN Red List of Threatened Species*, e.T41531A22004381.

153. Benda, P., & Mashkour, M. (2021). A finding of Eptesicus gobiensis in an ancient salt mine in Iran and notes on the status of this bat in the Middle East (Mammalia: Chiroptera). *Journal of the National Museum (Prague), Natural History Series*, *190*, 61–72.

154. Wilson, D. E., & Mittermeier, R. A. (2019). Vespertilionidae. In *Handbook of the Mammals of the World – Volume 9 Bats* (pp. 716–981). Barcelona: Lynx Edicions.

155. Linares, O. J., & Zabala, E. (2018). Refugios Diurnos de Eptesicus innoxius (Chiroptera, Vespertilionidae), en la Provincia de Guayas, Ecuador. *INVESTIGATIO*, *11*, 29–40.

156. Srinivasulu, C., & Csorba, G. (2019). Eptesicus lobatus. *IUCN Red List of Threatened Species*, e.T85200388A85200399.

157. Benda, P., Srinivasulu, C., & Srinivasulu, B. (2019). Rhyneptesicus nasutus. *IUCN Red List of Threatened Species*, e.T7935A22117147.

158. Rydell, J. (1993). Eptesicus nilssonii. *Mammalian Species*, *430*, 1–7.

159. Pachyomus, E. (2018). *A New Locality Record for the Asian Serotine Bat*. Kadoorie Farm & Botanic Garden.

160. Srinivasulu, C., Csorba, G., & Srinivasulu, B. (2019). Eptesicus pachyomus. *IUCN Red List of Threatened Species*, e.T85200202A85200236.

161. Khan, M. A. R. (2001). Status and distribution of bats in Bangladesh with notes on their ecology. *Zoos' Print Journal*, *16*(5), 479–483.

162. Godlevska, L., Kruskop, S. V., & Gazaryan, S. (2021). Eptesicus serotinus (amended version of 2020 assessment). *IUCN Red List of Threatened Species*, e.T85199559A195834153.

163. Soisook, P. (2017). Eudiscoderma thongareeae. *IUCN Red List of Threatened Species*, e.T80263386A95642210.

164. Soisook, P., Csorba, G., Bumrungsri, S., Francis, C. M., Bates, P., & Kingston, T. (2016). Eudiscopus denticulus. *IUCN Red List of Threatened Species*, e.T8168A22028419.

165. Best, T. L., Hunt, J. L., McWilliams, L. A., & Smith, K. G. (2002). Eumops auripendulus. *Mammalian Species*, *708*, 1–5.

166. McWilliams, L. A., Best, T. L., Hunt, J. L., & Smith, K. G. (2002). Eumops dabbenei. *Mammalian Species*, *707*, 1–3.

167. Solari, S. (2019). Eumops ferox. *IUCN Red List of Threatened Species*, e.T87994072A87994075.

168. Best, T. L., Kiser, W. M., & Rainey, J. C. (1997). Eumops glaucinus. *Mammalian Species*, *551*, 1–6.

169. Best, T. L., Hunt, J. L., McWilliams, L. A., & Smith, K. G. (2001). Eumops hansae. *Mammalian Species*, *687*, 1–3.

170. Sodré, M. M., Rosa, A. R. da, Gregorin, R., & Guimarães, M. M. (2008). Range extension for Thomas' Mastiff bat Eumops maurus (Chiroptera: Molossidae) in northern, central and southeastern Brazil. *Revista Brasileira de Zoologia*, *25*(2), 379–382.

171. Best, T. L., Kiser, W. M., & Freeman, P. W. (1996). Eumops perotis. *Mammalian Species*, *534*, 1–8.

172. Srinivasulu, B., & Srinivasulu, C. (2019). Hypsugo affinis. *IUCN Red List of Threatened Species*, e.T17324A22131594.

173. Miller, B., Reid, F., Arroyo-Cabrales, J., Cuarón, A. D., & de Grammont, P. C. (2016). Furipterus horrens. *IUCN Red List of Threatened Species*, e.T8771A21971535.

174. Monadjem, A., Taylor, P. J., Jacobs, D., & Cotterill, F. (2017). Glauconycteris beatrix. *IUCN Red List of Threatened Species*, e.T44791A22068514.

175. Schlitter, D. (2019). Glauconycteris humeralis. *IUCN Red List of Threatened Species*, e.T44795A22070303.

176. Monadjem, A., Taylor, P. J., Jacobs, D., & Cotterill, F. (2017). Glauconycteris poensis. *IUCN Red List of Threatened Species*, e.T44798A22069513.

177. Monadjem, A., Taylor, P. J., Jacobs, D., & Cotterill, F. (2017). Glauconycteris variegata. *IUCN Red List of Threatened Species*, e.T44800A22069727.

178. Webster, W. D., Handley, C. O., & Soriano, P. J. (1998). Glossophaga longirostris. *Mammalian Species*, *576*, 1–5.

179. Alvarez, J., Willig, M. R., Jones, J. K., & Webster, W. D. (1991). Glossophaga soricina. *Mammalian Species*, *379*, 1–7.

180. *Philippine Dwarf Fruit Bat Philippines*. (n.d.). Philippines Field Museum. Retrieved April 30, 2023, from https://philippines.fieldmuseum.org/natural-history/narrative/4127

181. Senawi, J., Hutson, A. M., & Kingston, T. (2020). Hesperoptenus doriae. *IUCN Red List of Threatened Species*, e.T9976A22076446.

182. Srinivasulu, B., & Srinivasulu, C. (2019). Hesperoptenus tickelli. *IUCN Red List of Threatened Species*, e.T9978A22075896.

183. Monadjem, A., Fahr, J., Hutson, A. M., Mickleburgh, S., & Bergmans, W. (2017). Hipposideros abae. *IUCN Red List of Threatened Species*, e.T10109A22097582.

184. Thong, V. D., & Bates, P. J. J. (2019). Hipposideros alongensis. *IUCN Red List of Threatened Species*, e.T80224880A95642200.

185. Bates, P. J. J., Bumrungsri, S., Francis, C., Csorba, G., & Oo, S. S. L. (2020). Hipposideros armiger. *IUCN Red List of Threatened Species*, e.T10110A22097743.

186. Armstrong, K. (2021). Hipposideros ater. *IUCN Red List of Threatened Species*, e.T80457009A22097974.

187. Monadjem, A., Juste, J., Bergmans, W., Mickleburgh, S., & Hutson A M & Fahr. (2017). Hipposideros beatus. *IUCN Red List of Threatened Species*, e.T10112A22098184.

188. Khan, F. A. A., Rajasegaran, P., & Shazali, N. (2020). Hipposideros bicolor. *IUCN Red List of Threatened Species*, e.T80258800A22095301.

189. Huang, J. C.-C., & W., S. (2016). Hipposideros breviceps. *IUCN Red List of Threatened Species*, e.T10114A22094935.

190. Richards, L. R., Cooper-Bohannon, R., Kock, D., Amr, Z. S. S., Mickleburgh, S., Hutson, A. M., Bergmans, W., & Aulagnier, S. (2019). Hipposideros caffer. *IUCN Red List of Threatened Species*, e.T80459007A22094271.

191. Armstrong, K. N., Wiantoro, S., & Lavery, T. H. (2021). Hipposideros calcaratus. *IUCN Red List of Threatened Species*, e.T10116A22094185.

192. Armstrong, K. (2021). Hipposideros cervinus. *IUCN Red List of Threatened Species*, e.T10118A22093732.

193. Douangboubpha, B., Srinivasulu, B., & Srinivasulu, C. (2019). Hipposideros cineraceus. *IUCN Red List of Threatened Species*, e.T10119A22093106.

194. Wilson, D. E., & Mittermeier, R. A. (2019). Hipposideridae. In *Handbook of the Mammals of the World – Volume 9 Bats* (pp. 227–258). Barcelona: Lynx Edicions.

195. Tanshi, I. (2020). Hipposideros curtus. *IUCN Red List of Threatened Species*, e.T10125A22096364.

196. Decher, J., & Fahr, J. (2005). Hipposideros cyclops. *Mammalian Species*, *763*, 1–7.

197. Aguilar, J., & Waldien, D. L. (2021). Hipposideros diadema. *IUCN Red List of Threatened Species*, e.T10128A22095445.

198. Mishra, R., & Dookia, S. (2016). Hipposideros durgadasi. *IUCN Red List of Threatened Species*, e.T10131A22090631.

199. Srinivasulu, B., & Srinivasulu, C. (2019). Hipposideros fulvus. *IUCN Red List of Threatened Species*, e.T10135A22089934.

200. Srinivasulu, B., & Srinivasulu, C. (2019). Hipposideros galeritus. *IUCN Red List of Threatened Species*, e.T10136A22090092.

201. Chakravarty, R., Srinivasulu, B., & Srinivasulu, C. (2016). Hipposideros hypophyllus. *IUCN Red List of Threatened Species*, e.T10138A22092730.

202. Milne, D. (2020). Hipposideros inornatus. *IUCN Red List of Threatened Species*, e.T136739A22035711.

203. Srinivasulu, C., & Srinivasulu, A. (2020). Hipposideros larvatus. *IUCN Red List of Threatened Species*, e.T85646564A22091287.

204. Monadjem, A., Fahr, J., Hutson, A. M., Mickleburgh, S., & Bergmans, W. (2017). Hipposideros megalotis. *IUCN Red List of Threatened Species*, e.T10150A22101286.

205. Srinivasulu, B., & Srinivasulu, C. (2018). In plain sight: Bacular and noseleaf morphology supports distinct specific status of Roundleaf Bats Hipposideros pomona Andersen, 1918 and Hipposideros gentilis Andersen, 1918 (Chiroptera: Hipposideridae). *Journal of Threatened Taxa*, *10*(8), 12018–12026.

206. Armstrong, K. N., Woinarski, J. C. Z., & Milne, D. J. (2021). Hipposideros stenotis. *IUCN Red List of Threatened Species*, e.T10163A22099463.

207. Russo, D., Maglio, G., Rainho, A., Meyer, C. F. J., & Palmeirim, J. M. (2011). Out of the dark: Diurnal activity in the bat Hipposideros ruber on São Tomé island (West Africa). *Mammalian Biology = Zeitschrift Fur Saugetierkunde*, *76*(6), 701–708.

208. Ossa, G., Lilley, T. M., Waag, A. G., Meierhofer, M. B., & Johnson, J. S. (2020). Roosting ecology of the southernmost bats, Myotis chiloensis and Histiotus magellanicus, in southern Tierra del Fuego, Chile. *Austral Ecology*, *45*(8), 1169–1178.

209. Barquez, R., & Diaz, M. (2016). Histiotus montanus. *IUCN Red List of Threatened Species*, e.T10202A22098875.

210. Simmons, N. B., & Voss, R. S. (1998). The mammals of Paracou, French Guiana, a Neotropical lowland rainforest fauna. Part 1, Bats. *Bulletin of the AMNH*, *237*, 1–219.

211. Kearney, T. (01 2013). Pipistrellus anchietae Anchieta's Pipistrelle. In M. Happold & D. C. D. Happlod (Eds.), *Mammals of Africa, Vol. IV: Hedgehogs, Shrews and Bats* (pp. 610–611). Bloomsbury Publishing.

212. Görföl, T., Furey, N. M., Bates, P. J. J., & Csorba, G. (2019). The Identity of “Falsistrellus” affinis from Myanmar and Cambodia and New Records of Hypsugo dolichodon from these Countries. *Acta Chiropterologica*, *20*(2), 301–309.

213. Jiang, T. L., & Feng, J. (2020). Ia io. *IUCN Red List of Threatened Species*, e.T10755A21993508.

214. Cockle, A., Kock, D., Stublefield, L., Howell, K. M., & Burgess, N. D. (1998). Bat assemblages in Tanzanian coastal forests. *Mammalia*, *62*(1), 53–68.

215. Sedlock, J. L., Heaney, L. R., Balete, D. S., & Ruedi, M. (2020). Philippine bats of the genus Kerivoula (Chiroptera: Vespertilionidae): Overview and assessment of variation in K. pellucida and K. whiteheadi. *Zootaxa*, *4755*(3), zootaxa.4755.3.2.

216. Jacobs, D. S., Barclay, R. M. R., & Corrie Schoeman, M. (2005). Foraging and roosting ecology of a rare insectivorous bat species, Laephotis wintoni (Thomas, 1901), Vespertilionidae. *Acta Chiropterologica*, *7*(1), 101–109.

217. Solari, S. (2018). Lampronycteris brachyotis. *IUCN Red List of Threatened Species*, e.T13376A22131330.

218. Solari, S. (2019). Lasiurus atratus. *IUCN Red List of Threatened Species*, e.T29607A22046087.

219. Shump, K. A., & Shump, A. U. (1982). Lasiurus cinereus. *Mammalian Species*, *185*, 1–5.

220. Aguiar, L., & Bernard, E. (2016). Lasiurus degelidus. *IUCN Red List of Threatened Species*, e.T136306A22018027.

221. Mancina, C. A. (2016). Lasiurus insularis. *IUCN Red List of Threatened Species*, e.T136754A22036556.

222. Webster, W. D., Jones, J. K., & Baker, R. J. (1980). Lasiurus intermedius. *Mammalian Species*, *132*, 1–3.

223. Duran, A. R. (2016). Lasiurus minor. *IUCN Red List of Threatened Species*, e.T136627A21987501.

232. Williams, S. L., & Genoways, H. H. (2007). Subfamily Phyllostominae Gray, 1825 from Mammals of South America. *Mammalogy Papers: University of Nebraska State Museum*.

233. Suárez-Castro, A. F., Ramírez-Chaves, H. E., & Velazco, P. M. (2017). Lonchorhina marinkellei (Chiroptera: Phyllostomidae). *Mammalian Species*, *49*(950), 76–80.

234. McCarthy, T. J., Davis, W. B., Hill, J. E., & Cruz, G. A. (1993). Bat (Mammalia: Chiroptera) records, early collectors, and faunal lists for northern Central America. *Annals of the Carnegie Museum*, *62*, 191–228.

235. Velazco, P. M., & Gardner, A. L. (2012). A new species of Lophostoma (Chiroptera: Phyllostomidae) from Panama. *Journal of Mammalogy*, *93*(2), 605–614.

236. Barquez, R., Diaz, M., Pineda, W., & Rodriguez, B. (2016). Lophostoma silvicolum. *IUCN Red List of Threatened Species*, e.T88149202A22041651.

242. Waldien, D. L., & Wiantoro, S. (2021). Megaerops kusnotoi. *IUCN Red List of Threatened Species*, e.T12945A22024115.

243. Tanalgo, K. C., & Tabora, J. A. G. (2015). Cave-dwelling bats (Mammalia: Chiroptera) and conservation concerns in South central Mindanao, Philippines. *Journal of Threatened Taxa*, *7*(15), 8185–8194.

244. Pennay, M. (2021). Melonycteris melanops. *IUCN Red List of Threatened Species*, e.T13139A21977021.

245. Simmons, N. B., Voss, R. S., & Fleck, D. W. (2002). A New Amazonian Species of Micronycteris (Chiroptera: Phyllostomidae) with Notes on the Roosting Behavior of Sympatric Congeners. *American Museum Novitates*, *2002*(3358), 1–16.

246. Solari, S. (2015). Micronycteris minuta. *IUCN Red List of Threatened Species*, e.T13380A22125019.

247. Ceballos, G. (2014). *Mammals of Mexico*. JHU Press.

248. Owen-Ashley, N. T., & Wilson, D. E. (1998). Micropteropus pusillus. *Mammalian Species*, *577*, 1–5.

249. Ortega, J., & Arita, H. T. (1997). Mimon bennettii. *Mammalian Species*, *549*, 1–4.

250. Goodman, & Steve. (2017). Miniopterus aelleni. *IUCN Red List of Threatened Species*, e.T81629770A95642245.

251. Goodman, S. (2017). Miniopterus ambohitrensis. *IUCN Red List of Threatened Species*, e.T81633128A95642255.

252. Schul, M. (2014). The Little Bent-wing Bat Miniopterus australis roosting in a tree hollow. *The Australian Zoologist*, *30*(3), 329–329.

253. Goodman, S. (2017). Miniopterus brachytragos. *IUCN Red List of Threatened Species*, e.T81629758A95642235.

254. Goodman, S. (2017). Miniopterus egeri. *IUCN Red List of Threatened Species*, e.T81633146A95642260.

255. Han, B. Y., Hua, P. Y., Gu, X. M., Miller-Butterworth, C. M., & Zhang, S. Y. (2008). Isolation and characterization of microsatellite loci in the long-fingered bat Miniopterus fuliginosus. *Molecular Ecology Resources*, *8*(4), 799–801.

256. Kim, S.-S., Choi, Y.-S., & Yoo, J.-C. (2014). The thermal preference and the selection of hibernacula in seven cave-dwelling bats. *Han'gug Hwan'gyeong Saengtae Haghoeji = Korean Journal of Environment and Ecology*, *47*(4), 258–272.

257. Fukui, D., & Sano, A. (2021). Miniopterus fuscus. *IUCN Red List of Threatened Species*, e.T13564A209553784.

258. Kofoky, A., Andriafidison, D., Ratrimomanarivo, F., Razafimanahaka, H. J., Rakotondravony, D., Racey, P. A., & Jenkins, R. K. B. (2007). Habitat use, roost selection and conservation of bats in Tsingy de Bemaraha National Park, Madagascar. In D. L. Hawksworth & A. T. Bull (Eds.), *Vertebrate Conservation and Biodiversity* (pp. 213–227). Springer Netherlands.

259. Reher, S., Rabarison, H., & Dausmann, K. (2019). Seasonal movements of insectivorous bat species in southwestern Madagascar. *Malagasy Nat*, *13*, 117–124.

260. Goodman, S. M., Weyeneth, N., Ibrahim, Y., Saïd, I., & Ruedi, M. (2010). A Review of the Bat Fauna of the Comoro Archipelago. *Acta Chiropterologica*, *12*(1), 117–141.

261. Armstrong, K. N., Wiantoro, S., & Aplin, K. (2021). Miniopterus macrocneme. *IUCN Red List of Threatened Species*, e.T136579A209529376.

262. Benda, P., & Piraccini, R. (2017). Miniopterus maghrebensis. *IUCN Red List of Threatened Species*, e.T81633156A95642265.

263. Armstrong, K. N., Wiantoro, S., & Aplin, K. (2021). Miniopterus magnater. *IUCN Red List of Threatened Species*, e.T13566A209529644.

264. Goodman, S. (2017). Miniopterus mahafaliensis. *IUCN Red List of Threatened Species*, e.T81629764A95642240.

265. Monadjem, A., Rakotoarivelo, A. R., & Jenkins, R. K. B. (2017). Miniopterus majori. *IUCN Red List of Threatened Species*, e.T40039A22061249.

266. Armstrong, K. N., Wiantoro, S., & Aplin, K. (2021). Miniopterus medius. *IUCN Red List of Threatened Species*, e.T13567A209529904.

267. McWilliam, A. N. (2010). Mating system of the bat Miniopterus minor (Chiroptera: Vespertilionidae) in Kenya, east Africa: A lek? *Ethology: Formerly Zeitschrift Fur Tierpsychologie*, *85*(4), 302–312.

268. Monadjem, A., Goodman, S. M., Stanley, W. T., & Appleton, B. (2013). A cryptic new species of Miniopterus from south-eastern Africa based on molecular and morphological characters. *Zootaxa*, *3746*, 123–142.

269. Monadjem, A., Griffin, M., Cotterill F Jacobs, & Taylor, P. J. (2017). Miniopterus natalensis. *IUCN Red List of Threatened Species*, e.T44862A22073129.

270. Rainho, A., Meyer, C. F. J., Thorsteinsdóttir, S., Juste, J., & Palmeirim, J. M. (2022). Current knowledge and conservation of the wild mammals of the Gulf of Guinea oceanic islands. In L. M. P. Ceríaco, R. F. de Lima, M. Melo, & R. C. Bell (Eds.), *Biodiversity of the Gulf of Guinea Oceanic Islands* (pp. 593–620). Springer, Cham.

271. Bouillard, N. (2021). Miniopterus paululus. *IUCN Red List of Threatened Species*, e.T136233A22001879.

272. Jenkins, R. K. B., & Rakotoarivelo, A. (2019). Miniopterus petersoni. *IUCN Red List of Threatened Species*, e.T81633135A22035230.

273. Bumrungsri, S., Bates, P. J. J., Molur, S., Srinivasulu, C., & Furey, N. (2021). Miniopterus pusillus. *IUCN Red List of Threatened Species*, e.T13569A22103542.

274. Waldien, D. L., & Brescia, F. (2020). Miniopterus robustior. *IUCN Red List of Threatened Species*, e.T13570A22103451.

275. Gray, P. A., Fenton, M. B., & Van Cakenberghe, V. (1999). Nycteris thebaica. *Mammalian Species*, *612*, 1–8.

276. Goodman, S. M., Ryan, K. E., Maminirina, C. P., Fahr, J., Christidis, L., & Appleton, B. (2007). Specific Status of Populations on Madagascar Referred to Miniopterus fraterculus (Chiroptera: Vespertilionidae), with Description of a New Species. *Journal of Mammalogy*, *88*(5), 1216–1229.

277. Armstrong, K. N., Wiantoro, S., & Aplin, K. (2021). Miniopterus tristis. *IUCN Red List of Threatened Species*, e.T13571A209530159.

285. Miller, B., Reid, F., Arroyo-Cabrales, J., Cuarón, A. D., & de Grammont, P. C. (2016). Molossus sinaloae. *IUCN Red List of Threatened Species*, e.T13650A22106433.

286. Wilson, D. E., & Mittermeier, R. A. (2019). Mormoopidae. In *Handbook of the Mammals of the World – Volume 9 Bats* (pp. 424–443). Barcelona: Lynx Edicions.

287. Goodman, S. M., van Vuuren, B. J., Ratrimomanarivo, F., Probst, J.-M., & Bowie, R. C. K. (2008). Specific Status of Populations in the Mascarene Islands Referred to Mormopterus acetabulosus (Chiroptera: Molossidae), with Description of a New Species. *Journal of Mammalogy*, *89*(5), 1316–1327.

288. Ratrimomanarivo, F. H., Goodman, S. M., Taylor, P. J., Melson, B., & Lamb, J. (2009). Morphological and genetic variation in Mormopterus jugularis (Chiroptera: Molossidae) in

different bioclimatic regions of Madagascar with natural history notes. *Mammalia*, *73*(2), 110–129.

289. Azhar, M. I., & Rossiter, S. J. (2020). Murina aenea. *IUCN Red List of Threatened Species*, e.T13936A22091750.

290. Soisook, P. (2017). Murina balaensis. *IUCN Red List of Threatened Species*, e.T84487939A84487985.

291. Furey, N., & Csorba, G. (2021). Murina eleryi. *IUCN Red List of Threatened Species*, e.T84557696A84557699.

292. Khan, F. A. A., & Rosli, Q. (2020). Murina rozendaali. *IUCN Red List of Threatened Species*, e.T13945A22097407.

293. Watari, Y., & Funakoshi, K. (2013). Use of dead-leaf foliage as day-roosts by the Ryukyu tube-nosed bat, *Murina ryukyuana*. *Mammalian Science*, *53*(2), 331–334.

294. Braun, J. K., Layman, Q. D., & Mares, M. A. (2009). Myotis albescens (Chiroptera: Vespertilionidae). *Mammalian Species*, *846*, 1–9.

295. Jeon, Y. S., Kim, S. C., Han, S. H., & Chung, C. U. (2019). First utilization record of bat box for bat conservation in Korea. *Journal of Environmental Science International*, *28*(1), 163–167.

296. Chung, C. U., Kim, S. C., & Han, S. H. (2013). Diurnal Roosts Selection and Home Range Size in the Myotis Aurascens (Chiroptera: Vespertilionidae) Inhabiting a Rural Area. *Journal of Environmental Science International*, *22*(9), 1227–1234.

297. Zhigalin, A. (2019). New data on David's myotis, Myotis davidii (Peters, 1869) (Mammalia, Chiroptera, Vespertilionidae), in Siberia and the Urals. *Biodiversity Data Journal*, *7*, e34211.

298. Fukui, D., Sano, A., & Kruskop, S. V. (2019). Myotis bombinus. *IUCN Red List of Threatened Species*, e.T14149A22061650.

299. Gazaryan, S., Kruskop, S. V., & Godlevska, L. (2020). Myotis brandtii. *IUCN Red List of Threatened Species*, e.T85566997A195857637.

306. Arroyo-Cabrales, J., & Ospina-Garces, S. (2016). Myotis findleyi. *IUCN Red List of Threatened Species*, e.T14159A22058800.

307. Kim, S. S., Choi, Y. S., Kim, B. H., & Yoo, J. C. (2009). The current distribution and habitat preferences of hibernating Myotis formosus in Korea. *Journal of Ecology and Field Biology*, *32*(3), 191–195.

308. Ho, Y.-Y. (2008). *Causes and Consequences of Roost Switching by the Bat Myotis Formosus (Vespertilionidae)*. Faculty of Graduate Studies, University of Western Ontario.

309. Vincenot, C. E., Preble, J. H., Huang, J. C.-C., Collazo, A. M., & Kamal, A. (2021). Myotis frater. *IUCN Red List of Threatened Species*, e.T85566806A22056940.

310. Decher, J., & Choate, J. R. (1995). Myotis grisescens. *Mammalian Species*, *510*, 1–7.

311. Dietz, C., Gazaryan, A., Papov, G., Dundarova, H., & Mayer, F. (2016). Myotis hajastanicus is a local vicariant of a widespread species rather than a critically endangered endemic of the Sevan lake basin (Armenia). *Mammalian Biology = Zeitschrift Fur Saugetierkunde*, *81*(5), 518–522.

312. Hernández-Meza, B., Domínguez-Castellanos, Y., & Ortega, J. (2005). Myotis keaysi. *Mammalian Species*, *785*, 1–3.

313. Novaes, R. L. M., Hintze, F., & Moratelli, R. (2022). Myotis lavali (Chiroptera: Vespertilionidae). *Mammalian Species*, *54*(1018).

314. Araújo, R. A., Amaro, B. D., Talamoni, S. A., & Godinho, H. P. (2013). Seasonal reproduction of yellowish myotis, Myotis levis (Chiroptera: Vespertilionidae), from a Neotropical highland. *Journal of Morphology*, *274*(11), 1230–1238.

315. Moratelli, R., & Wilson, D. E. (2014). A new species of Myotis (Chiroptera, Vespertilionidae) from Bolivia. *Journal of Mammalogy*, *95*(4), E17–E25.

316. Buckley, D. J., Lundy, M. G., Boston, E. S. M., Scott, D. D., Gager, Y., Prodöhl, P., Marnell, F., Montgomery, W. I., & Teeling, E. C. (2013). The spatial ecology of the whiskered bat (Myotis mystacinus) at the western extreme of its range provides evidence of regional adaptation. *Mammalian Biology = Zeitschrift Fur Saugetierkunde*, *78*(3), 198–204.

317. Simal, F., Smith, L., Doest, O., de Lannoy, C., Franken, F., Zaandam, I., Simal, D., & Nassar, J. M. (2022). Bat Inventories at Caves and Mines on the Islands of Aruba, Bonaire and Curaçao, and Proposed Conservation Actions. *Acta Chiropterologica*, *23*(2), 455–474.

318. Srinivasulu, B., & Srinivasulu, C. (2018). Myotis nipalensis. *IUCN Red List of Threatened Species*, e.T136495A21976309.

319. Solari, S. (2018). Myotis oxyotus. *IUCN Red List of Threatened Species*, e.T14187A22067211.

320. Jones, G., Parsons, S., Zhang, S., Stadelmann, B., Benda, P., & Ruedi, M. (2006). Echolocation calls, wing shape, diet and phylogenetic diagnosis of the endemic Chinese bat Myotis pequinius. *Acta Chiropterologica / Museum and Institute of Zoology, Polish Academy of Sciences*, *8*(2), 451–463.

321. Fukui, D., & Sano, A. (2020). Myotis petax. *IUCN Red List of Threatened Species*, e.T85342726A85342734.

322. Chang, Y., Song, S., Li, A., Zhang, Y., Li, Z., Xiao, Y., Jiang, T., Feng, J., & Lin, A. (2019). The roles of morphological traits, resource variation and resource partitioning associated with the dietary niche expansion in the fish-eating bat Myotis pilosus. *Molecular Ecology*, *28*(11), 2944–2954.

323. Núñez-Rojo, M. P., Arroyo-Cabrales, J., Rivera-Téllez, E., & Medellín, R. A. (2020). Summer roosts of “The revenant” flat-headed myotis, Myotis planiceps. *Journal of Mammalogy*, *101*(6), 1526–1532.

324. Novaes, R. L. M., Wilson, D. E., Ruedi, M., & Moratelli, R. (2018). The taxonomic status of Myotis aelleni Baud, 1979 (Chiroptera, Vespertilionidae). *Zootaxa*, *4446*(2), 257–264.

325. Baron, B. (2007). A look at the Chiropteran Fauna of the Maltese Islands: Towards an effective Action Plan for their conservation. *Xjenza*, *12*(120201), 1.

326. Baron, B., & Vella, A. (2010). A preliminary analysis of the population genetics of Myotis punicus in the Maltese Islands. *Hystrix* .

327. Novaes, R. L. M., Souza, R. de F., & Moratelli, R. (2017). Myotis riparius (Chiroptera: Vespertilionidae). *Mammalian Species*, *49*(946), 51–56.

328. Sedlock, J. L., Stuart, A. M., Horgan, F. G., Hadi, B., Como Jacobson, A., Alviola, P. A., & Alvarez, J. D. V. (2019). Local-Scale Bat Guild Activity Differs with Rice Growth Stage at Ground Level in the Philippines. *Diversity*, *11*(9), 148.

329. Moratelli, R. (2012). Myotis simus (Chiroptera: Vespertilionidae). *Mammalian Species*, *44*(892), 26–32.

330. Blood, B. R., & Clark, M. K. (1998). Myotis vivesi. *Mammalian Species*, *588*, 1–5.

331. Ratcliffe, J. M. (2002). Myotis welwitschii. *Mammalian Species*, *701*, 1–3.

332. O'Donnell, C. (2021). Mystacina robusta. *IUCN Red List of Threatened Species*, e.T14260A22070387.

339. Schneck, J., Hawkins, F., Cox, N., Mair, L., Thieme, A., & Sexton, J. (2023). Species Threat Abatement and Recovery in Cameroon and Kenya: Findings from a STAR assessment to support biodiversity conservation using high-resolution data. *Gland, Switzerland: IUCN*.

340. Jacobs, D. (2019). Neoromicia melckorum. *IUCN Red List of Threatened Species*, e.T44922A22047486.

341. Lausen, C. L., & Barclay, R. M. R. (2005). Pipistrellus nanus. *Mammalian Species*, *784*, 1–7.

342. Monadjem, A., & Jacobs, D. (2017). Neoromicia somalica. *IUCN Red List of Threatened Species*, e.T44925A22046866.

343. Smith, Paul. (2008). FAUNA Paraguay Handbook of the Mammals of Paraguay Number 22 Noctilio albiventris. p1-16.

344. Wilson, D. E., & Mittermeier, R. A. (2019). Noctilionidae. In *Handbook of the Mammals of the World – Volume 9 Bats* (pp. 404–411). Barcelona: Lynx Edicions.

345. Fukui, D., Dewa, H., Katsuta, S., & Sato, A. (2013). Bird predation by the birdlike noctule in Japan. *Journal of Mammalogy*, *94*(3), 657–661.

346. Leonardo, M., & Medeiros, F. M. (2011). Preliminary data about the breeding cycle and diurnal activity of the azorean bat (Nyctalus azoreum). *Açoreana, Suplemento*, *7*, 139–148.

347. Wojtaszyn, G., Lesiński, G., & Rutkowski, T. (2021). Seasonal Dynamics of Occupation of Bat Boxes by Bats in Forests of South-western Poland. *Acta Zoologica Bulgarica*, *73*(3).

348. Printz, L., Tschapka, M., & Vogeler, A. (2021). The common noctule bat (Nyctalus noctula): population trends from artificial roosts and the effect of biotic and abiotic parameters on the probability of occupation. *Journal of Urban Ecology*, *7*(1).

349. Cel'uch, M., & Kanuch, P. (2005). Winter activity and roosts of the noctule (Nyctalus noctula) in an urban area (Central Slovakia). *Lynx*, *36*, 39–45.

350. Decher, J., Hoffmann, A., Schaer, J., Norris, R. W., Kadjo, B., Astrin, J., Monadjem, A., & Hutterer, R. (2015). Bat diversity in the Simandou Mountain Range of Guinea, with the description of a new white-winged vespertilionid. *Acta Chiropterologica / Museum and Institute of Zoology, Polish Academy of Sciences*, *17*(2), 255–282.

351. Hickey, M. B. C., & Dunlop, J. M. (2000). Nycteris grandis. *Mammalian Species*, *632*, 1–4.

352. Wilson, D. E., & Mittermeier, R. A. (2019). Nycteridae. In *Handbook of the Mammals of the World – Volume 9 Bats* (pp. 374–386). Barcelona: Lynx Edicions.

353. Monadjem, A., Bergmans, W., Mickleburgh, S., & Hutson, A. M. (2017). Nycteris hispida. *IUCN Red List of Threatened Species*, e.T14930A22012843.

354. Pottie, S. A., Lane, D. J. W., Kingston, T., & Lee, B. P. Y.-H. (2005). The microchiropteran bat fauna of Singapore. *Acta Chiropterologica / Museum and Institute of Zoology, Polish Academy of Sciences*, *7*(2), 237–247.

355. Mickleburgh, S., Hutson, A. M., Bergmans, W.: Cotterill, F.P.D., & Jacobs, D. (2019). Nycteris vinsoni. *IUCN Red List of Threatened Species*, e.T44696A22074669.

356. Johnston, D. S. (2006). Nycticeinops schlieffeni. *Mammalian Species*, *798*, 1–4.

357. Armstrong, K. (2021). Nyctimene aello. *IUCN Red List of Threatened Species*, e.T14954A22008855.

358. Loveless, A. M., & McBee, K. (2017). Nyctimene robinsoni (Chiroptera: Pteropodidae). *Mammalian Species*, *49*(949), 68–75.

359. Jones, J. K., & Arroyo-Cabrales, J. (1990). Nyctinomops aurispinosus. *Mammalian Species*, *350*, 1–3.

360. Kumirai, A., & Jones, J. K. (1990). Nyctinomops femorosaccus. *Mammalian Species*, *349*, 1–5.

361. Flores-Quispe, M., Calizaya-Mamani, G., Portugal-Zegarra, G., Alvarado, G. A., Pacheco-Castillo, J., & Rengifo, E. M. (2019). Contributions to the natural history of Mormopterus kalinowskii (Chiroptera: Molossidae) in the southwest of Peru. *THERYA*, *10*(3), 343.

367. Armstrong, K. N., Lumsden, L. F., & Reardon, T. B. (2022). Nyctophilus gouldi. *IUCN Red List of Threatened Species*, e.T218360733A218360491.

368. Driessen, M., Brereton, R., & Pauza, M. (2011). Status and conservation of bats in Tasmania. *The Australian Zoologist*, *35*(Special Issue), 324–336.

369. Parnaby, H. E. (2009). A taxonomic review of Australian Greater Long-eared Bats previously known as Nyctophilus timoriensis (Chiroptera:Vespertilionidae) and some associated taxa. *The Australian Zoologist*, *35*(1), 39–81.

370. Fenton, M. B., Taylor, P. J., Jacobs, D. S., Richardson, E. J., Bernard, E., Bouchard, S., Debaeremaeker, K. R., ter Hofstede, H., Hollis, L., Lausen, C. L., Lister, J. S., Rambaldini, D., Ratcliffe, J. M., & Reddy, E. (2002). Researching little-known species: the African bat Otomops martiensseni (Chiroptera: Molossidae). *Biodiversity & Conservation*, *11*(9), 1583–1606.

371. Armstrong, K. (2021). Otomops secundus. *IUCN Red List of Threatened Species*, e.T15650A209524157.

380. Sampaio, E., Lim, B., & Peters, S. (2016). Peropteryx trinitatis. *IUCN Red List of Threatened Species*, e.T136790A22035534.

398. Kruskop, S. V., Solovyeva, E. N., & Kaznadzey, A. D. (2018). Unusual Pipistrelle: Taxonomic Position of the Malayan Noctule (Pipistrellus stenopterus; Vespertilionidae; Chiroptera). *Zoological Studies* , *57*, e60.

399. Seamark, E. C. J. (2009). Observations. *African Bat Conservation News*, *21*, 4.

400. Kipson, M., Gazaryan, S., & Horáček, I. (2020). Savi's Pipistrelle Hypsugo savii (Bonaparte, 1837). In *Handbook of the Mammals of Europe* (pp. 1–18). Springer International Publishing.

401. Newman, B. A., Loeb, S. C., & Jachowski, D. S. (2021). Winter roosting ecology of tricolored bats (Perimyotis subflavus) in trees and bridges. *Journal of Mammalogy*, *102*(5), 1331–1341.

402. *Northern Pipistrelle*. (n.d.). The Australian Museum. Retrieved May 18, 2023, from https://australian.museum/learn/animals/bats/northern-pipistrelle/

403. Garbino, G. S. T., & Piñeros, C. A. C. (2022). First record of Platyrrhinus albericoi (Chiroptera: Phyllostomidae) roosting in Ficus americana (Moraceae). *Notas Sobre Mamíferos Sudamericanos*, *4*.

404. Ferrell, C. S., & Wilson, D. E. (1991). Platyrrhinus helleri. *Mammalian Species*, *373*, 1–5.

405. Velazco, P. M., Gardner, A. L., & Patterson, B. D. (2010). Systematics of the Platyrrhinus helleri species complex (Chiroptera: Phyllostomidae), with descriptions of two new species. *Zoological Journal of the Linnean Society*, *159*(3), 785–812.

406. Solari, S. (2016). Platyrrhinus ismaeli. *IUCN Red List of Threatened Species*, e.T136232A22002129.

407. Velazco, P. (2015). Platyrrhinus masu. *IUCN Red List of Threatened Species*, e.T136577A21998517.

408. Velazco, P. (2015). Platyrrhinus nigellus. *IUCN Red List of Threatened Species*, e.T136317A22020785.

409. Tavares, V. da C., & Velazco, P. M. (2010). Platyrrhinus recifinus (Chiroptera: Phyllostomidae). *Mammalian Species*, *42*(859), 119–123.

410. Velazco, P. M., Guevara, L., & Molinari, J. (2018). Systematics of the broad-nosed bats, Platyrrhinus umbratus (Lyon, 1902) and P. nigellus (Gardner and Carter, 1972) (Chiroptera: Phyllostomidae), based on genetic, morphometric, and ecological niche analyses. *Neotropical Biodiversity*, *4*(1), 119–133.

411. Aulagnier, S, Benda, & P. (2019). Plecotus christii. *IUCN Red List of Threatened Species*, e.T44931A22045680.

412. Kruskop, S. V., & Fukui, D. (2019). Plecotus ognevi. *IUCN Red List of Threatened Species*, e.T136598A21996784.

425. Pennay, M., Lavery, T. H., & Roberts, B. (2021). Pteropus capistratus. *IUCN Red List of Threatened Species*, e.T84891540A22012219.

426. Lavery, T. (2017). Pteropus cognatus. *IUCN Red List of Threatened Species*, e.T136397A22014516.

427. Tait, J., Perotto-Baldivieso, H. L., McKeown, A., & Westcott, D. A. (2014). Are flying-foxes coming to town? Urbanisation of the spectacled flying-fox (Pteropus conspicillatus) in Australia. *PloS One*, *9*(10), e109810.

428. Hahn, M. B., Epstein, J. H., Gurley, E. S., Islam, M. S., Luby, S. P., Daszak, P., & Patz, J. A. (2014). Roosting behaviour and habitat selection of Pteropus giganteus reveals potential links to Nipah virus epidemiology. *The Journal of Applied Ecology*, *51*(2), 376–387.

429. Fisher, D., Helgen, K., & Allison, A. (2021). Pteropus howensis. *IUCN Red List of Threatened Species*, e.T18728A22080900.

430. Jones, D. P., & Kunz, T. H. (2000). Pteropus hypomelanus. *Mammalian Species*, *639*, 1–6.

431. Kunz, T. H., & Jones, D. P. (2000). Pteropus vampyrus. *Mammalian Species*, *2000*(642), 1–6.

432. Granek, E. (2002). Conservation of Pteropus livingstonii based on roost site habitat characteristics on Anjouan and Moheli, Comoros islands. *Biological Conservation*, *108*(1), 93–100.

433. Chaiyes, A., Duengkae, P., Wacharapluesadee, S., Pongpattananurak, N., Olival, K. J., & Hemachudha, T. (2017). Assessing the distribution, roosting site characteristics, and population of Pteropus lylei in Thailand. *The Raffles Bulletin of Zoology*, *65*, 670–680.

434. Leary, T., & Helgen, K. (2020). Pteropus macrotis. *IUCN Red List of Threatened Species*, e.T18735A22082074.

435. Lavery, T. (2017). Pteropus mahaganus. *IUCN Red List of Threatened Species*, e.T18736A22082180.

436. Wiles, G. J., & Johnson, N. C. (2004). Population Size and Natural History of Mariana Fruit Bats (Chiroptera: Pteropodidae) on Sarigan, Mariana Islands. *Pacific Science*, *58*(4), 585–596.

437. Oleksy, R. Z., Ayady, C. L., Tatayah, V., Jones, C., Howey, P. W., Froidevaux, J. S. P., Racey, P. A., & Jones, G. (2019). The movement ecology of the Mauritian flying fox (Pteropus niger): a long-term study using solar-powered GSM/GPS tags. *Movement Ecology*, *7*, 12.

438. Buden, D. W., Helgen, K. M., & Wiles, G. J. (2013). Taxonomy, distribution, and natural history of flying foxes (Chiroptera, Pteropodidae) in the Mortlock Islands and Chuuk State, Caroline Islands. *ZooKeys*, *345*, 97–135.

439. Wiles, G. J., Engbring, J., & Otobed, D. (1997). Abundance, biology, and human exploitation of bats in the Palau Islands. *Journal of Zoology* , *241*(2), 203–227.

440. Mildenstein, T. (2016). Pteropus pohlei. *IUCN Red List of Threatened Species*, e.T18750A22085786.

450. Tsang, S. (2016). Pteropus temminckii. *IUCN Red List of Threatened Species*, e.T18762A22088270.

451. Miller, C. A., & Wilson, D. E. (1997). Pteropus tonganus. *Mammalian Species*, *552*, 1–6.

452. Hayes, F. E., & Engbring, J. (2020). Historic and recent status of the Kosrae flying fox (Pteropus ualanus) (Chiroptera: Pteropidae) on Kosrae, Micronesia. *Journal of Asia-Pacific Biodiversity*, *13*(2), 141–150.

453. Entwistle, A., & Nadia Corp. (1997). Status and distribution of the Pemba flying fox Pteropus voeltzkowi. *Oryx: The Journal of the Fauna Preservation Society*, *31*(2), 135–142.

454. Lavery, T. H., & Fisher, D. (2017). Pteropus woodfordi. *IUCN Red List of Threatened Species*, e.T18769A22089578.

455. Huang, J. C.-C., Jazdzyk, E. L., Nusalawo, M., Maryanto, I., Maharadatunkamsi, Wiantoro, S., & Kingston, T. (2014). A Recent Bat Survey Reveals Bukit Barisan Selatan Landscape as a Chiropteran Diversity Hotspot in Sumatra. *Acta Chiropterologica*, *16*(2), 413–449.

456. Burgin, C. (2019). Rhinolophidae. In D. E. Wilson & R. A. Mittermeier (Eds.), *Handbook of the Mammals of the World – Volume 9 Bats* (pp. 280–332). Barcelona: Lynx Edicions.

457. Ith, S., Bumrungsri, S., Furey, N. M., Bates, P. J., Wonglapsuwan, M., Khan, F. A. A., Thong, V. D., Soisook, P., Satasook, C., & Thomas, N. M. (2015). Taxonomic implications of geographical variation in Rhinolophus affinis (Chiroptera: Rhinolophidae) in mainland Southeast Asia. *Zoological Studies* , *54*, e31.

458. Nasri, N., Maulany, R. I., & Hamzah, A. S. (2021). Diversity of cave-dwelling bats in Leang Londrong, Bantimurung-Bulusaraung National Park: An initial field note. *IOP Conference Series: Earth and Environmental Science*, *886*(1), 012059.

459. Jayaraj, V. K. (2020). Rhinolophus borneensis. *IUCN Red List of Threatened Species*, e.T19527A21982599.

460. Ith, S., Soisook, P., Bumrungsri, S., Kingston, T., Puechmaille, S. J., Struebig, M. J., Bu, S. S. H., Thong, V. D., Furey, N. M., Thomas, N. M., & Bates, P. J. J. (2011). A Taxonomic Review of Rhinolophus coelophyllus Peters 1867 and R. shameli Tate 1943 (Chiroptera: Rhinolophidae) in Continental Southeast Asia. *Acta Chiropterologica*, *13*(1), 41–59.

461. Aul, B., Bates, P. J. J., Harrison, D. L., & Marimuthu, G. (2014). Diversity, distribution and status of bats on the Andaman and Nicobar Islands, India. *Oryx: The Journal of the Fauna Preservation Society*, *48*(2), 204–212.

462. Swanepoel, R., Smit, S. B., Rollin, P. E., Formenty, P., Leman, P. A., Kemp, A., Burt, F. J., Grobbelaar, A. A., Croft, J., Bausch, D. G., Zeller, H., Leirs, H., Braack, L. E. O., Libande, M. L., Zaki, S., Nichol, S. T., Ksiazek, T. G., Paweska, J. T., & International Scientific and Technical Committee for Marburg Hemorrhagic Fever Control in the Democratic Republic of Congo. (2007). Studies of reservoir hosts for Marburg virus. *Emerging Infectious Diseases*, *13*(12), 1847–1851.

463. UHrin, M., Gh, S. B., Bücs, S., Paunović, M., Miková, E., Juhász, M., Csősz, I., Estók, P., ULín, M. f., Gombkötő, P., Jére, C., Barti, L., Karapandža, B., Matis, Š., na Gy, Z. L., szodoray-parádi, F., & Benda, P. (2012). Revision of the occurrence of Rhinolophus euryale in the Carpathian region, Central Europe. *Vespertilio*, *16*, 289–328.

464. Armstrong, K. N., & Aplin, K. (2021). Rhinolophus euryotis (amended version of 2017 assessment). *IUCN Red List of Threatened Species*, e.T84372418A209537830.

465. Denys, C., Kadjo, B., Missoup, A. D., Monadjem, A., & Aniskine, V. (2013). New records of bats (Mammalia: Chiroptera) and karyotypes from Guinean Mount Nimba (West Africa). *Italian Journal of Zoology*, *80*(2), 279–290.

466. Flanders, J., Frick, W. F., Nziza, J., Nsengimana, O., Kaleme, P., Dusabe, M. C., Ndikubwimana, I., Twizeyimana, I., Kibiwot, S., Ntihemuka, P., Cheng, T. L., Muvunyi, R., & Webala, P. (2022). Rediscovery of the critically endangered Hill's horseshoe bat (Rhinolophushilli) and other new records of bat species in Rwanda. *Biodiversity Data Journal*, *10*, e83546.

467. Fahr, J., Djossa, B. A., & Vierhaus, H. (2006). Rapid assessment of bats (Chiroptera) in Déré, Diécké and Mt. Béro classified forests, southeastern Guinea; including a review of the distribution of bats in Guinée Forestière. In *A rapid biological assessment of three classified forests in southeastern Guinea* (pp. 168–180). Conservation International.

468. Wu, Y., Motokawa, M., & Harada, M. (2008). A new species of horseshoe bat of the genus Rhinolophus from China (Chiroptera: Rhinolophidae). *Zoological Science*, *25*(4), 438–443.

469. Abantas, A. D., & Nuneza, O. M. (2014). Species diversity of terrestrial vertebrates in Mighty Cave, Tagoloan, Lanao Del Norte, Philippines. *Journal of Biodiversity and Environmental Sciences*, *5*(6), 122–132.

470. Monadjem, A. (2016). Rhinolophus kahuzi. *IUCN Red List of Threatened Species*, e.T82347204A82347492.

471. Csorba, G., & Bates, P. J. J. (2016). Rhinolophus keyensis. *IUCN Red List of Threatened Species*, e.T19577A21992519.

472. Brown, K. M., & Dunlop, J. (1997). Rhinolophus landeri. *Mammalian Species*, *567*, 1–4.

473. Thong, V. D., Thanh, H. T., Soisook, P., & Csorba, G. (2019). Rhinolophus luctus. *IUCN Red List of Threatened Species*, e.T19548A21977086.

474. Fahr, J., Vierhaus, H., Hutterer, R., & Kock, D. (2002). A revision of the Rhinolophus maclaudi species group with the description of a new species from West Africa (Chiroptera: Rhinolophidae). *Myotis*, *40*(95), 126.

475. Liang, J., He, X., Peng, X., Xie, H., & Zhang, L. (2020). First record of existence of Rhinolophus malayanus (Chiroptera, Rhinolophidae) in China. *Mammalia*, *84*(4), 362–365.

476. Murphy, M. (2014). Roost caves of the Eastern Horseshoe Bat Rhinolophus megaphyllus Gray, 1834 (Chiroptera: Rhinolophidae) in the Pilliga forest in northern inland New South Wales, Australia. *The Australian Zoologist*, *37*(1), 117–126.

477. Benda, P., Ivanova, T., Gaisler, J., Gueorguieva, A., Petrov, B., & Others. (2003). Bats (Mammalia: Chiroptera) of the eastern Mediterranean. Part 3. Review of bat distribution in Bulgaria. *Acta Societatis Zoologicae Bohemicae*, *67*(4), 245–357.

478. Taylor, P. J., Stoffberg, S., Monadjem, A., Schoeman, M. C., Bayliss, J., & Cotterill, F. P. D. (2012). Four new bat species (Rhinolophus hildebrandtii complex) reflect Plio-Pleistocene divergence of dwarfs and giants across an Afromontane archipelago. *PloS One*, *7*(9), e41744.

479. Sun, K. (2020). Rhinolophus osgoodi. *IUCN Red List of Threatened Species*, e.T19557A21992735.

480. Eger, J. L., & Fenton, M. B. (2003). Rhinolophus paradoxolophus. *Mammalian Species*, *731*, 1–4.

481. Bates, P. J. J., Bumrungsri, S., Csorba, G., & Mao, X. G. (2019). Rhinolophus pearsonii. *IUCN Red List of Threatened Species*, e.T19559A21993105.

482. Sun, K. (2020). Rhinolophus rex. *IUCN Red List of Threatened Species*, e.T19562A21994639.

483. Peterhans, J. C. K., Fahr, J., Huhndorf, M. H., & Kaleme, P. (2013). Bats (Chiroptera) from the Albertine Rift, eastern Democratic Republic of Congo, with the description of two new species of the Rhinolophus maclaudi group. *Bonn Zool*, *62*, 186–202.

484. Cotterill, F. P. D. (2002). A new species of horseshoe bat (Microchiroptera: Rhinolophidae) from south−central Africa: with comments on its affinities and evolution, and the characterization of rhinolophid species. *Journal of Zoology* , *256*(2), 165–179.

485. Wu, Y., & Thong, V. D. (2011). A new species of Rhinolophus (Chiroptera: Rhinolophidae) from China. *Zoological Science*, *28*(3), 235–241.

486. Volleth, M., Khan, F. A. A., Müller, S., Baker, R. J., Arenas-Viveros, D., Stevens, R. D., Trifonov, V., Liehr, T., Heller, K.-G., & Sotero-Caio, C. G. (2021). Cytogenetic Investigations in Bornean Rhinolophoidea Revealed Cryptic Diversity in Rhinolophus sedulus Entailing Classification of Peninsular Malaysia Specimens as a New Species. *Acta Chiropterologica*, *23*(1), 1–20.

487. Molur, S., Marimuthu, G., Srinivasulu, C., Mistry, S., Hutson, A. M., Bates, P. J. J., Walker, S., Priya, K. P., & Priya, A. R. B. (2002). Status of South Asian Chiroptera. *Conservation Action Management Plan (CAMP) Workshop Report, Zoo Outreach Organisation, 320pp*.

488. Taylor, P. (2017). Rhinolophus smithersi. *IUCN Red List of Threatened Species*, e.T64588371A64589277.

489. Soisook, P., Bumrungsri, S., Satasook, C., Thong, V. D., Bu, S. S. H., Harrison, D. L., & Bates, P. J. J. (2008). A taxonomic review of Rhinolophus stheno and R. malayanus (Chiroptera: Rhinolophidae) from continental Southeast Asia: an evaluation of echolocation call frequency

in discriminating between cryptic species. *Acta Chiropterologica / Museum and Institute of Zoology, Polish Academy of Sciences*, *10*(2), 221–242.

490. Basumatary, S., & Bera, S. (2014). Modern pollen record on bat guano deposit from siju cave and its implication to palaeoecological study in south Garo hills of meghalaya, India. *Journal of Cave and Karst Studies: The National Speleological Society Bulletin*, *76*(3), 173–183.

491. Heaney, L., Balete, D., Dolar, L., & Ong, P. (01 1998). A Synopsis of the Mammalian Fauna of the Philippine Islands. *Fieldiana. Zoology*, *88*, 1–61.

492. Mutumi, G. L., Jacobs, D. S., & Winker, H. (2016). Sensory Drive Mediated by Climatic Gradients Partially Explains Divergence in Acoustic Signals in Two Horseshoe Bat Species, Rhinolophus swinnyi and Rhinolophus simulator. *PloS One*, *11*(1), e0148053.

493. Bates, P. J. J., Thi, M. M., Nwe, T., Bu, S. S. H., Mie, K. M., Nyo, N., Khaing, A. A., Aye, N. N., Oo, T., & Mackie, I. (2004). A review of Rhinolophus (Chiroptera: Rhinolophidae) from Myanmar, including three species new to the country. *Acta Chiropterologica / Museum and Institute of Zoology, Polish Academy of Sciences*, *6*(1), 23–48.

494. Thong, V. D., Limbert, H., & Limbert, D. (2022). First Records of Bats (Mammalia: Chiroptera) from the World’s Largest Cave in Vietnam. *Diversity*, *14*(7), 534.

495. Monadjem, A. (2020). Rhinolophus willardi. *IUCN Red List of Threatened Species*, e.T82346260A82347169.

496. Zhou, Z.-M., Guillén-Servent, A., Lim, B. K., Eger, J. L., Wang, Y.-X., & Jiang, X.-L. (2009). A New Species from Southwestern China in the Afro-Palearctic Lineage of the Horseshoe Bats (Rhinolophus). *Journal of Mammalogy*, *90*(1), 57–73.

497. Cooper-Bohannon, R., & Monadjem, A. (2020). Rhinolophus ziama. *IUCN Red List of Threatened Species*, e.T44786A22068674.

498. Rinehart, J. B., & Kunz, T. H. (2006). Rhinophylla pumilio. *Mammalian Species*, *791*, 1–5.

499. Benda, P., Reiter, A., Al-Jumaily, M., Nasher, A. K., & Hulva, P. (2009). A new species of mouse-tailed bat (Chiroptera: Rhinopomatidae: Rhinopoma) from Yemen. *Journal of the National Museum (Prague), Natural History Series*, *177*(6), 53–68.

500. Qumsiyeh, M. B., & Jones, J. K. (1986). Rhinopoma hardwickii and Rhinopoma muscatellum. *Mammalian Species*, *263*, 1–5.

501. Schlitter, D. A., & Qumsiyeh, M. B. (1996). Rhinopoma microphyllum. *Mammalian Species*, *542*, 1–5.

502. Srinivasulu, B., & Srinivasulu, C. (2019). Rhinopoma muscatellum. *IUCN Red List of Threatened Species*, e.T19602A21997131.

503. Soriano, P. J., Naranjo, M. E., & Fariñas, M. R. (2004). A new subspecies of the little desert bat (Rhogeessa minutilla) from a Venezuelan semiarid enclave. *Mammalian Biology = Zeitschrift Fur Saugetierkunde*, *69*(6), 439–443.

504. Téllez, H. L. A., Iñiguez-Davalos, L. I., Olvera-Vargas, M., Vargas-Contreras, J. A., & Herrera-Lizaola, O. A. (2018). Bats associated to caves in Jalisco, Mexico. *THERYA*, *9*(1).

505. Kwiecinski, G. G., & Griffiths, T. A. (1999). Rousettus egyptiacus. *Mammalian Species*, *611*, 1–9.

506. Nangoy, M., Ransaleleh, T., Lengkong, H., Koneri, R., Latinne, A., & Kyes, R. C. (2021). Diversity of fruit bats (Pteropodidae) and their ectoparasites in Batuputih Nature Tourism Park, Sulawesi, Indonesia. *Biodiversitas Journal of Biological Diversity*, *22*(6).

507. Bergmans, W. (1997). Taxonomy and biogeography of African fruit bats (Mammalia,Megachiroptera). 5. The genera Ussonycteris Andersen, 1912, Myonycteris

Matschie, 1899 and Megaloglossus Pagenstecher, 1885; general remarks and conclusions; annex: key to all species. *Beaufortia*, *47*(2), 11–90.

517. Tanshi, I. (2020). Scotonycteris occidentalis. *IUCN Red List of Threatened Species*, e.T84466273A84466694.

518. Obitte, B. (2021). Scotonycteris zenkeri. *IUCN Red List of Threatened Species*, e.T84464403A192236400.

519. Jacobs, D. S., & Barclay, R. M. R. (2009). Niche Differentiation in Two Sympatric Sibling Bat Species, Scotophilus dinganii and Scotophilus mhlanganii. *Journal of Mammalogy*, *90*(4), 879–887.

520. Srinivasulu, B., Srinivasulu, C., & Venkateshwarlu, P. (2010). First record of Lesser Yellow House Bat Scotophilus kuhlii Leach, 1821 from Secunderabad, Andhra Pradesh, India with a note on its diet. *Journal of Threatened Taxa*, 1234–1236.

521. Goodman, S. M., Ratrimomanarivo, F. H., & Randrianandrianina, F. H. (2006). A new species of Scotophilus (Chiroptera: Vespertilionidae) from western Madagascar. *Acta Chiropterologica*, *8*(1), 21–37.

522. Ratrimomanarivo, F. H., & Goodman, S. M. (2005). The first records of the synanthropic occurrence of Scotophilus spp. on Madagascar. *African Bat Conservation News*, *6*(3), 5.

523. Monadjem, A., Raabe, T., Dickerson, B., Silvy, N., & McCleery, R. (2010). Roost use by two sympatric species of Scotophilus in a natural environment. *South African Journal of Wildlife Research-24-Month Delayed Open Access*, *40*(1), 73–76.

524. Lumsden, L. F., Reardon, T. B., & Armstrong, K. N. (2021). Scotorepens orion (amended version of 2020 assessment). *IUCN Red List of Threatened Species*, e.T14945A209531493.

532. Carneiro, L., Monteiro, L. R., & Nogueira, M. R. (2022). Sturnira tildae (Chiroptera: Phyllostomidae). *Mammalian Species*, *54*(1015).

533. Law, B. S. (1993). Roosting and foraging ecology of the Queensland blossom bat (Syconycteris australis) in north-eastern New South Wales: flexibility in response to seasonal variation. *Wildlife Research* , *20*(4), 419–431.

534. Shah, T. A., Srinivasulu, C., Kaur, H., Srinivasulu, B., & Devender, G. (2014). New distribution records of Tadarida aegyptiaca E. Geoffroy, 1818 (Mammalia: Chiroptera: Molossidae) from Karnataka, India. *International Journal of Fauna and Biological Studies*, *1*, 41–43.

535. Turton, M., & Hoye, G. (2011). Note: The use of a building for breeding by the white-striped freetail-bat Tadarida australis at Newington, Sydney, New South Wales. In *The Biology and Conservation of Australasian Bats* (pp. 460–463). Royal Zoological Society of NSW Mosman.

536. Fukui, D., & Sano, A. (2019). Tadarida insignis. *IUCN Red List of Threatened Species*, e.T136716A22036641.

537. Armstrong, K. (2021). Austronomus kuboriensis. *IUCN Red List of Threatened Species*, e.T136201A22009294.

538. Monadjem, A., & Cotterill, W. (2017). Tadarida lobata. *IUCN Red List of Threatened Species*, e.T21317A22121550.

544. Monadjem, A., Fahr, J., Mickleburgh, S., Racey, P. A., Hutson, A. M., Ravino, J., & Bergmans, W. (2017). Taphozous mauritianus. *IUCN Red List of Threatened Species*, e.T21460A22111004.

545. Phelps, K., Csorba, G., Bumrungsri, S., Helgen, K., Francis, C., Bates, P., Gumal, M., Balete, D. S., Heaney, L., Molur, S., & Srinivasulu, C. (2019). Taphozous melanopogon. *IUCN Red List of Threatened Species*, e.T21461A22110277.

546. Monadjem, A., Molur, S., Hutson, A. M., Amr, Z. S. S., Kock, D., Mickleburgh, S., & Bergmans, W. (2020). Taphozous perforatus. *IUCN Red List of Threatened Species*, e.T21463A166505490.

547. Kyle Armstrong (South Australian Museum), Reardon, T., Andrew Burbidge (IUCN SSC Australasian Marsupial and Monotreme Specialist Group), & John Woinarski (Natural Resources, Environment and The Arts, NT). (2021). Taphozous troughtoni. *IUCN Red List of Threatened Species*, e.T21466A209539933.

548. Rosa, R. O. L., Silva, C. H. A., Oliveira, T. F., Silveira, M., & Aguiar, L. M. S. (2020). Type of shelter and first description of the echolocation call of disk-winged bat (*Thyroptera devivoi*). *Biota Neotropica*, *20*(2), e20190821.

554. Solari, S. (2018). Tonatia saurophila. *IUCN Red List of Threatened Species*, e.T41530A22004890.

555. Ramasindrazana, B., & Goodman, S. (2014). Documented record of Triaenops menamena (family Hipposideridae) in the central highlands of Madagascar. *African Bat Conservation News*, *25*.

566. Coroiu, I. (2016). Vespertilio murinus. *IUCN Red List of Threatened Species*, e.T22947A22071456.

567. Fukui, D., Sano, A., & Kruskop, S. V. (2019). Vespertilio sinensis. *IUCN Red List of Threatened Species*, e.T22949A22071812.

568. Gomes, L. A. C., Maas, A. C. S., Godoy, M. S. M., Martins, M. A., Pedrozo, A. R., & Peracchi, A. L. (2018). Ecological considerations on Xeronycteris vieirai: an endemic bat species from the Brazilian semiarid macroregion. *Mastozoologia Neotropical*, *25*(1), 81–88.
